# Genome assembly and annotation of the olive grass mouse *Abrothrix olivacea* reveal transcriptomic and cellular adaptations across contrasting biomes

**DOI:** 10.1101/2025.09.12.675871

**Authors:** M. Magallanes Alba, Baricalla Agustin, Feijoo Matias, D’Elia Guillermo, E. Naya Daniel, P. Lessa Enrique

## Abstract

*Abrothrix olivacea* (Waterhouse, 1837), the olive grass mouse, is a widely distributed sigmodontine rodent that inhabits a broad range of environments, from the hyper arid deserts of southernmost Perú and northern Chile to the Patagonian steppe to the humid temperate rainforests of southern South America. Its extensive ecological breadth, coupled with physiological adaptations to water scarcity, makes it an ideal model for studying environmental responses and phenotypic plasticity. Here, we present the first *de novo* scaffold-level genome assembly of *A. olivacea*, generated from short-read DNA sequencing. The 2.25 Gb assembly achieved a scaffold N50 of 123 Mb and a BUSCO completeness score of 98.61%, indicating high sequence completeness. Genome annotation identified 21,476 protein-coding genes, providing a valuable resource for evolutionary, ecological, and functional genomics. As a case study, we used this reference genome to explore gene expression and genetic divergence in kidney tissue from individuals inhabiting contrasting environments: the southern Andean rainforest and the Patagonian steppe. By integrating single-cell transcriptomic data from *Mus musculus*, we performed cell type deconvolution, revealing environment-specific expression patterns linked to renal function. This new genomic resource opens avenues for investigating local adaptation, population structure, and conservation genetics in one of South America’s most ecologically versatile and widely distributed rodents.

## Background

The olive grass mouse, *Abrothrix olivacea* (Waterhouse 1837), a sigmodontine of the tribe Abrotrichini, exhibits remarkable ecological adaptability, thriving in diverse habitats in southern South America, across various regions in southernmost Perú, Chile and Argentina, from hyper arid deserts in southernmost Peru and northern Chile to the Valdivian and Magellanic temperate rainforests in the south, as well as the Chilean Mediterranean region, the Patagonian steppes and Fuegian grasslands (Patton et al. 2015). This widespread distribution across heterogeneous landscapes offers valuable opportunities to explore how phenotypic variability is structured and whether it correlates with environmental features (Quiroga-Carmona and D’Elía 2022). *A. olivacea* exhibits exceptional urine concentration capability and a relative medullary thickness comparable to desert-adapted species (McNab 2002; Al‐kahtani et al. 2004). Higher tolerance to water shortage in populations from xeric habitat has already been demonstrated (Bozinovic et al. 2011). Based on variation in morphology, coloration patterns, and DNA sequence data (Mann 1978; Rodríguez-Serrano et al. 2006; Pizarro et al. 2012; Quiroga-Carmona et al. 2023), distinct subspecies of *A. olivacea* have been described and six divergent mitochondrial phylogeogroups have been found along its distribution (Lessa et al. 2010; Quiroga-Carmona et al. 2022). More recently, studies of both mitochondrial DNA and nuclear SNPs have further documented substantial geographical variation in this species, both latitudinally and across biomes (Giorello et al. 2021). All these characteristics make *A. olivacea* an excellent focal case for the study of geographical variation in response to environmental changes.

While a *de novo* transcriptome of *A. olivacea* has been previously been reported (Giorello et al. 2014; Giorello et al. 2018), a complete genome sequence for this species is not yet available. Therefore, we sequenced the entire genome of the olive mouse and subsequently performed its assembly and annotation. Here we report the first *de novo* scaffold-level genome assembly for the olive mouse based on short-read DNA sequencing. The genome assembly yielded a 2.25 Gb sequence with a scaffold N50 of 123 Mb. Benchmarking with Glires universal single-copy gene orthologs (BUSCOs) showed 98.8% completeness. Genome annotation resulted in the identification of 21,476 coding sequences. This breakthrough provides essential groundwork for numerous advancements. It enables genome-wide analyses, detection of structural variations, improves gene annotation accuracy, and facilitates comparative genomics. Furthermore, this genomic data is crucial for conservation efforts, aiding in the identification of genetically distinct populations, understanding their evolutionary potential, and developing strategies for their preservation amidst environmental challenges.

We leveraged the newly generated genome assembly to investigate gene expression differences in the kidneys of individuals from the southern Andean rainforests and the Patagonian steppe and to assess genetic divergence among populations. Furthermore, employing a deconvolution technique with the kidney single-cell profile of the house mouse (*Mus musculus*), we connected the observed gene expression variations to known kidney cell type functions. Also, beyond *A. olivacea*, other rodent models have been investigated for their physiological and transcriptomic responses to water limitation. In particular, the cactus mouse (*Peromyscus eremicus*), a desert specialist from North America, and the lesser Egyptian jerboa (*Jaculus jaculus*), native to the Saharan and Middle Eastern deserts, have been studied under experimental dehydration challenges (MacManes 2017; Gillard et al. 2023). The availability of genome assemblies and transcriptomic data for these species offers a valuable comparative framework to contextualize the responses observed in *A. olivacea*, enabling us to explore shared and lineage-specific mechanisms of kidney adaptation to arid environments.

## Methods

### Study site and sample collection

The individual used for genome sequencing was a male *A. olivacea* (field catalog number GD1411; housed at Colección de Mamíferos Universidad Austral de Chile under number UACH 8896). The specimen was collected on April 21, 2011, at Fundo San Martín, San José de la Mariquina, Los Ríos Region, Chile (39°38.954′ S, 73°11.553′ W).

Kidney expression profiles were sourced from earlier studies that aimed to characterize the kidney transcriptome and deepen insights into the molecular adjustments to water availability (Giorello et al. 2014; Giorello et al. 2018). Raw sequences are accessible at the BioProjects PRJNA471316 and PRJNA231324 in NCBI.

### Genomic DNA extraction, sequencing and assembly

High molecular weight genomic DNA was extracted from frozen liver tissue using a standard DNA purification kit (Qiagen) following the manufacturer’s recommendations. The extracted DNA was treated with RNase A to remove RNA contaminants. DNA quality and concentration were assessed with a NanoDrop spectrophotometer and a Qubit fluorometer, and integrity was confirmed by agarose gel electrophoresis. A whole genome shotgun approach on the Illumina HiSeq™ 2500 platform was employed to sequence the genome. This process yielded a total data of 34.5 Gb as paired-end library with insert sizes of about 400 base pairs (bp) and 29.9 Gb as two mate-paired libraries with insert sizes of 3 kb and 8 kb (**Table 1**).

**Table 1.**
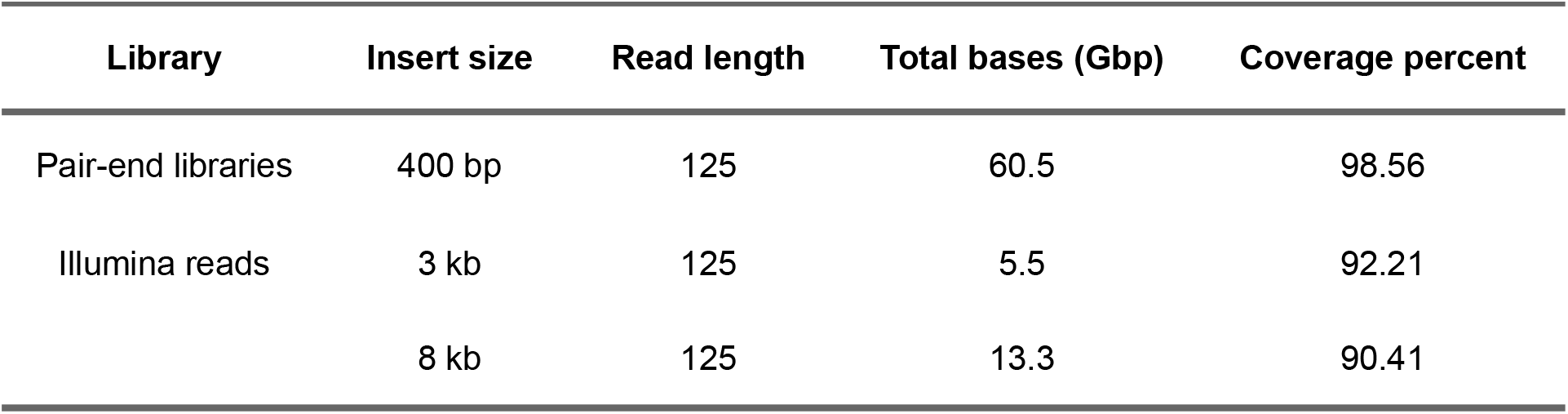
Statistics of genome sequencing of the Olive mouse *Abrothrix olivacea*.

| Library | Insert size | Read length | Total bases (Gbp) | Coverage percent |
| --- | --- | --- | --- | --- |
| Pair-end libraries | 400 bp | 125 | 60.5 | 98.56 |
| Illumina reads | 3 kb | 125 | 5.5 | 92.21 |
|  | 8 kb | 125 | 13.3 | 90.41 |

Initially, Jellyfish (Marçais and Kingsford 2011) and Genomescope 2.0 (Ranallo-Benavidez et al. 2020) were used to perform k-mer analysis by short insert size library reads before assembly; the genome size was estimated to be 2.25 Gb. Quality control on the raw sequence data was checked with FastQC v0.12.1 (Andrews et al. 2010), low-quality reads and adaptors were removed using Trim Galore v.0.6.10 (Krueger et al. 2023).

The initial assembly was created with MaSuRCA (Zimin et al. 2013). Genome assembly was further aided by the Redundans pipeline (Pryszcz and Gabaldón 2016). Scaffolding improvements were achieved using RagTag in scaffold mode (Alonge et al. 2022), leveraging the *Phyllotis vaccarum* reference genome (GCA_033318265.1) to guide structural organization. Finally, abyss-sealer (Jackman et al. 2017) was used to fill gaps in the scaffolds. **Figure 1** illustrates this process.

**Figure 1.**
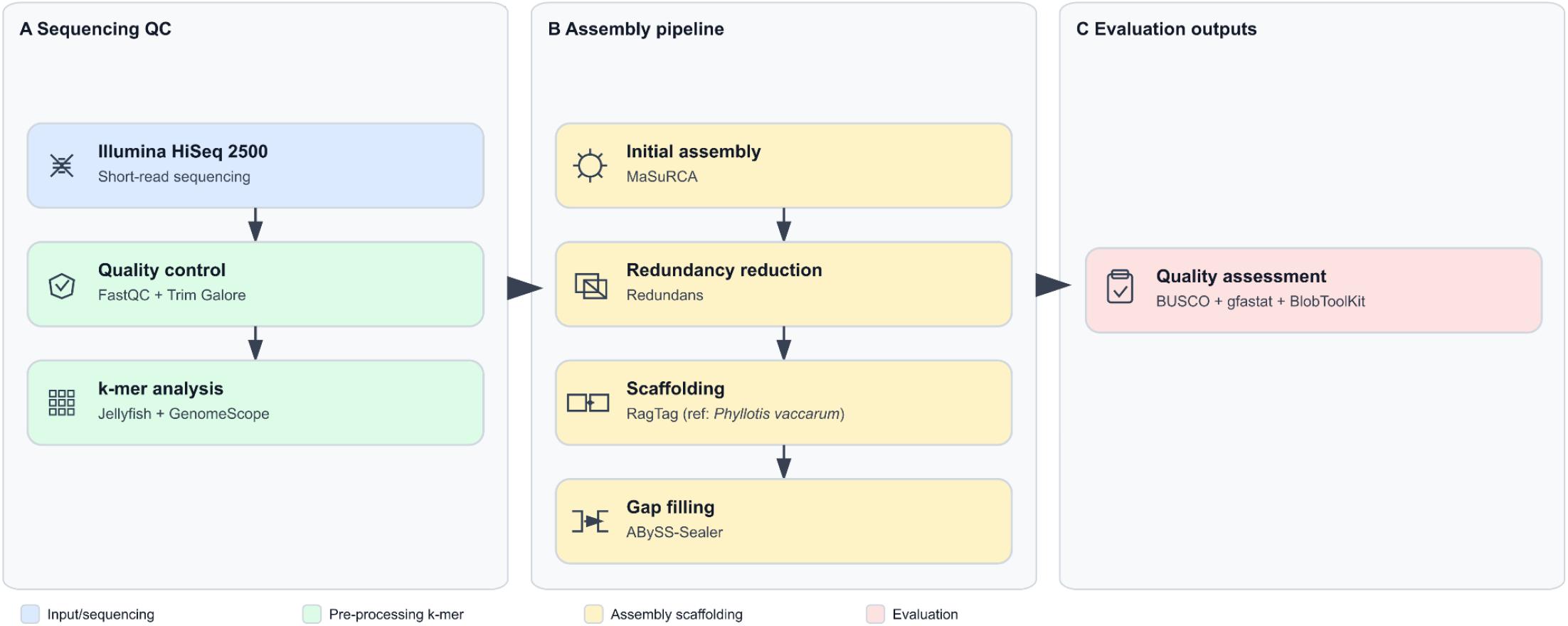
Workflow of the genome assembly and annotation of the olive mouse. Panel A shows sequencing and pre-processing steps, including Illumina short-read sequencing, quality control, and k-mer analysis. Panel B summarizes the assembly pipeline, consisting of initial assembly with MaSuRCA, redundancy reduction with Redundans, scaffolding with RagTag using *Phyllotis vaccarum* as reference, and gap filling with ABySS-Sealer. Panel C presents the evaluation stage, where assembly completeness and quality were assessed with BUSCO gfastats and BlobToolKit. Colors indicate different categories: input/sequencing (blue), pre-processing (green), assembly/scaffolding (yellow), and evaluation (pink).

Genome completeness was assessed using BUSCO v.5.8.0 (Simão et al. 2015; Tegenfeldt et al. 2025) and compleasm v.0.2.6 (Huang and Li 2023) with the Glires data set. Summary statistics were calculated with gfastats v1.3.6 (Formenti et al. 2022). To evaluate assembly quality, we analyzed the genome assembly with BlobToolKit v. 4.4.6 (Challis et al. 2020).

### Structural and functional annotation

Prior to genome annotation, repeat elements were identified and characterized using RepeatModeler v2.0.5 (Flynn et al. 2020) to generate a *de novo* repeat library. Subsequently, repetitive elements were masked using RepeatMasker v4.1.5 (Tarailo‐Graovac and Chen 2009), incorporating both the *de novo* library and known repeats from RepBase. Two independent gene annotations were generated. First, BRAKER3 v3.0.8 (Gabriel et al. 2023) was run using RNA-seq data aligned to the genome as transcriptomic evidence, together with protein homology evidence from the Glires RefSeq protein database. Second, gene prediction transfer was performed using LiftOn v1.0.5 (Chao et al. 2024), employing *Mus musculus* (GCF_000001635.27) as reference genome annotation. Finally, both annotations were merged using AGAT tools (agat_sp_merge.pl and agat_sp_fix_features_locations_duplicated.pl) (Jacques Dainat et al. 2024) to generate the final consensus annotation. The structural annotation obtained was also analyzed using BUSCO v.5.8.0.

### Differential expression

Kidney RNA-Seq data were initially quality-trimmed using TrimGalore v0.6.10. High-quality reads were subsequently aligned to the assembled genome using HISAT2 v2.2.1 (Kim et al. 2015). Gene-level read counts were obtained with FeatureCounts (Liao et al. 2014). To minimize noise and ensure robust results, only genes with counts per million (CPM) ≥ 1 in at least one experimental group were retained for downstream analyses. Consistency among biological replicates was evaluated using principal component analysis (PCA) implemented via the prcomp function in R (R Core Team 2025). Read counts were normalized using the TMM method, and differential expression analysis was conducted using the edgeR (Robinson et al. 2010) and limma (Ritchie et al. 2015) packages. The model included a single main factor (biome), comparing forest and steppe samples. Genes with adjusted p-values < 0.05 (Benjamini and Hochberg 1995) were considered significantly differentially expressed (DEGs).

In parallel, the publicly available datasets from *Peromyscus eremicus* (BioProject PRJEB19330; (MacManes 2017)) and *Jaculus jaculus* (BioProject PRJNA935758; (Gillard et al. 2023)) were reanalyzed but using the newly available genomes as reference (GCA_949786415.1 and, GCA_020740685.1, respectively), and all available samples from that study were processed for comparative purposes.

### Biological and functional analysis of DEGs

The ClusterProfiler v4.12.0 package (Yu et al. 2012) was utilized for gene set enrichment analysis (GSEA) of DEGs. This analysis was conducted on the complete set of genes using the Mouse MSigDB (Dolgalev 2022; Castanza et al. 2023). Gene Ontology and Kyoto Encyclopedia of Genes and Genomes (KEGG) were tested separately for each data set. Furthermore, to identify the source of DEGs at the cell type level from bulk RNA-seq data, a user-defined criterion of cell type signature gene set was created containing cluster marker genes for cell types identified in single-cell sequencing studies of mouse kidney (Park et al. 2018; Agarwal et al. 2021; Balzer et al. 2022). Normalized enrichment score (NES) was used to indicate the strength of enrichment. |NES| ≥ 1.0 and adjusted or nominal p-value < 0.05 were defined as statistical significance.

### Transcriptomic deconvolution of bulk kidney RNA-seq

To estimate the cellular composition of kidney bulk RNA-seq samples from *Abrothrix olivacea*, we performed transcriptomic deconvolution using CIBERSORTx (Newman et al. 2019). As a reference, we used the Tabula Muris Droplet dataset (Tabula Muris Consortium et al. 2018), a publicly available scRNA-seq atlas of the house mouse (*Mus musculus*). We filtered the dataset to retain only kidney-derived cells. To generate the cell-type signature matrix, we used the AggregateExpression function from Seurat v5 (Hao et al. 2024) to compute the average expression per annotated cell type, based on raw counts. This matrix was used as input for CIBERSORTx. Bulk RNA-seq counts were obtained using featureCounts (Liao et al. 2014) and preprocessed with edgeR (Robinson et al. 2010). Genes with zero counts across all samples were removed. Normalization to counts per million (CPM) was performed using edgeR. The resulting CPM matrix was used as input for CIBERSORTx along with the cell-type signature matrix derived from mouse kidney scRNA-seq data. Deconvolution was run with 100 permutations using default settings. Given the phylogenetic distance between *M. musculus* and *A. olivacea*, gene identifiers were manually curated to retain only orthologous genes shared across species, ensuring compatibility between bulk and single-cell matrices.

### Mitogenome assembly and annotation

The mitogenome was assembled, annotated and visualized using MitoZ v3.6 (Meng et al. 2019) The mitogenome of *Akodon montensis* was used as the starting sequence (NC_025746.1). The resulting circularised mitogenome was 16428 bp in length and contained the standard 37 vertebrate mitochondrial genes (13 protein-coding, 22 tRNAs, and 2 rRNAs) (Figure S1).

## Results and Discussion

### Genome assembly and annotation

We obtained a high-contiguity genome assembly for *Abrothrix olivacea*, consisting of 105 scaffolds with a total length of 2.25 Gb with a heterozygosity rate of 0.942% (**Figure 2**). The largest scaffold reached 200.1 Mb, and the assembly showed a scaffold N50 of 122.5 Mb with an L50 of 8, indicating strong assembly continuity. The assembly contains 630,661 gaps, totaling 7.78 Mb, which represents only 0.35% of the entire assembly. Overall, these metrics indicate a high-quality genome assembly with excellent scaffold continuity (**Table 2**). The genome assembly was compared to publicly available assemblies of related rodent species. Both the scaffold and contig N50 values of the *A. olivacea* genome fall within the mid-range of those observed across the selected taxa, indicating a comparable level of continuity despite differences in sequencing technologies and assembly approaches. This project has been deposited at NCBI under number PRJNA471316.

**Table 2.** Comparative genome assembly statistics for A. olivacea and related rodent species. Accession numbers (Acc. n^o^.) and references are provided where available. BUSCO completeness was assessed using the glires_odb12 database.

| Species | Reference | Acc_no | Year | Sequencing Technology | Genome Size_Gb | Scaffolds | Contigs | Scaffold_N50_Mb | Contig_N50_Mb | GC_percent | BUSCO_percent |
| --- | --- | --- | --- | --- | --- | --- | --- | --- | --- | --- | --- |
| *Abrothrix olivacea* | This study | xx | 2,025 | Illumina | 2.25 | 105 | 630,766 | 122.5 | 0.01 | 40.00 | 98.8 |
| *Sigmodon hispidus* | Colella et al. 2020 | GCA_004025045.1 | 2,019 | Illumina | 2.70 | 883,546 | 897,290 | 101.4 | 68.00 | 40.00 | 96.0 |
| *Phyllotis vaccharum* | Steppan et al. 2022 | GCA_033318265.1 | 2,023 | PacBio; Illumina | 2.70 | 304 | 437 | 147.9 | 23.20 | 42.50 | 97.3 |
| *Peromyscus eremicus* | Tigano et al. 2020 | GCA_949786415.1 | 2,023 | PacBio | 2.90 | 4,296 | 6,899 | 97.2 | 1.30 | 42.00 | 97.9 |
| *Jaculus jaculus* | Rhie et al. 2021 | GCA_020740685.1 | 2,021 | PacBio | 2.90 | 159 | 715 | 158.2 | 22.10 | 42.00 | 97.9 |
| *Peromyscus crinitus* | Tigano et al. 2022 | Dna Zoo | 2,023 | Illumina + 10X + Hi-C | — | 43,773 | 78,433 | 97.4 | 0.14 | 42.16 | — |

**Figure 2.**
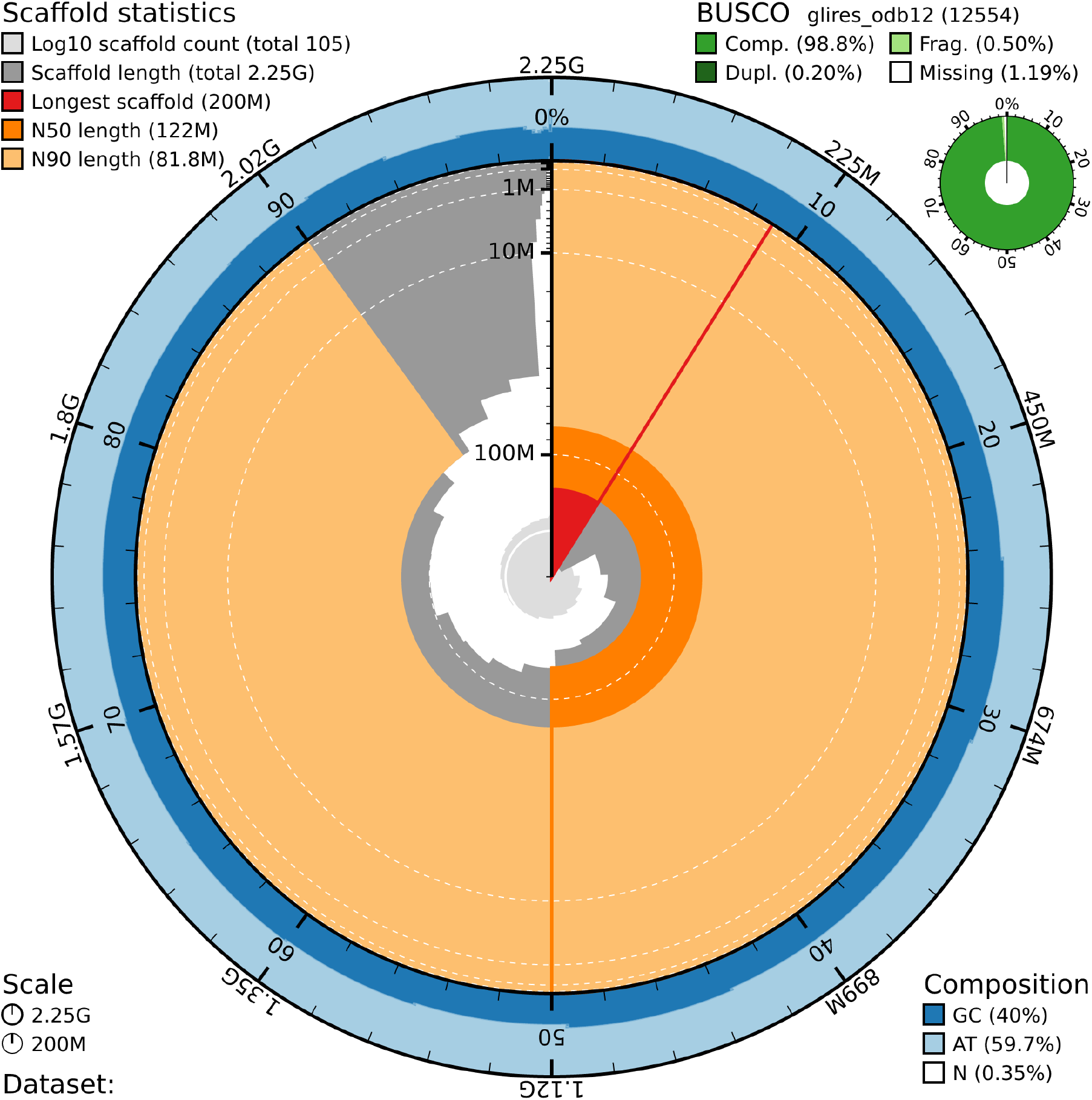
BlobToolKit snail plot summarizing key statistics of the genome assembly, including scaffold length distribution, N50, GC content, and completeness (BUSCO scores). This visualization provides an overview of the assembly quality and characteristics.

The resulting assembly exhibits a high level of sequence completeness, as assessed by BUSCO analysis, which indicated that 98.8% of the expected Glires single-copy orthologs were present in the genome and its annotation (**Figure 2 and Table 2**). In addition, RNA-seq read mapping confirmed that the assembled transcripts were well represented within the *A. olivacea* scaffolds. As an example, **Figure 3** shows RNA-seq read alignments across the *Atp1a2* gene, which is expressed at moderate levels and has a typical multiexonic structure. The uniform read coverage across exons and the presence of clear exon–intron boundaries support the quality and completeness of the genome assembly. A total of 21,476 protein-coding genes were predicted in the assembled genome with an average coding DNA sequence length of 1,711 bp (**Table 3**).

**Table 3.**
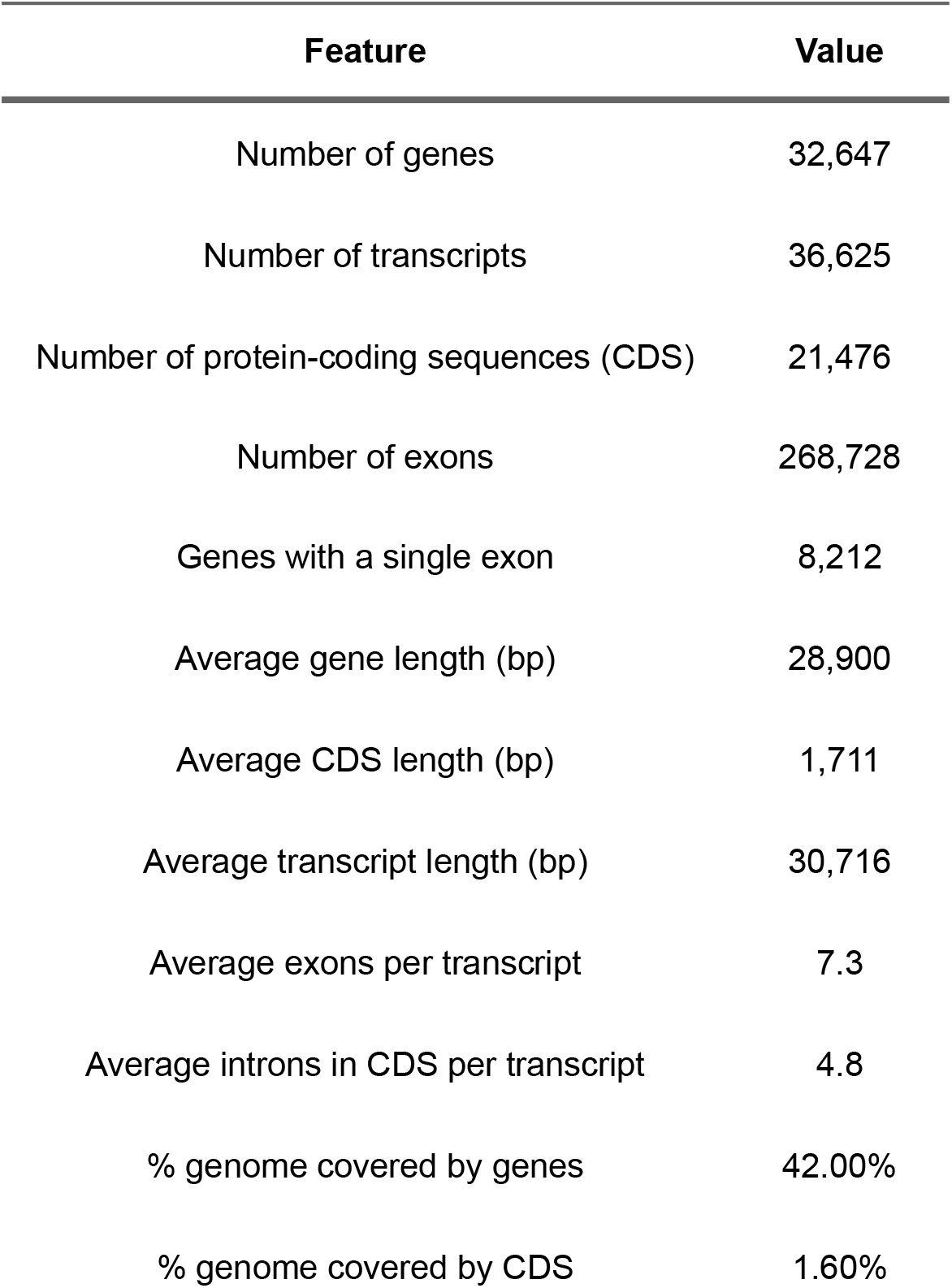
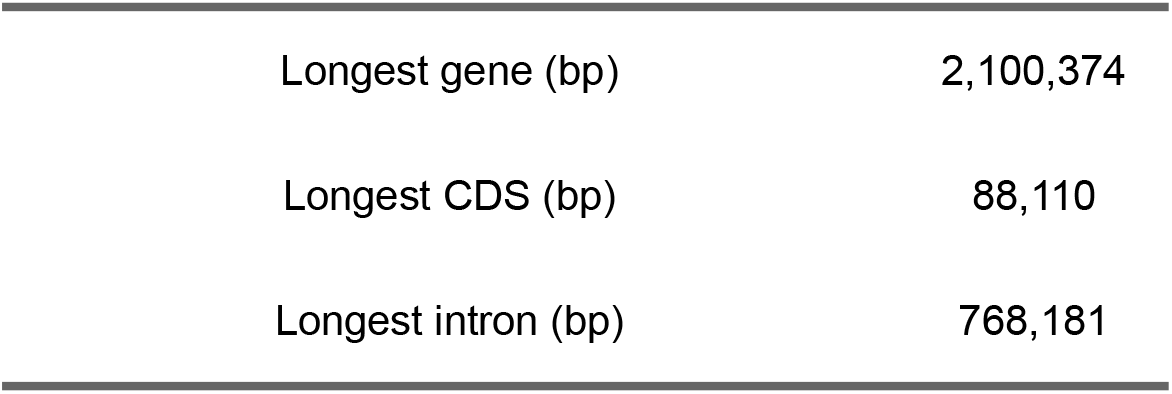
Summary of protein-coding gene annotation.

| Feature | Value |
| --- | --- |
| Number of genes | 32,647 |
| Number of transcripts | 36,625 |
| Number of protein-coding sequences (CDS) | 21,476 |
| Number of exons | 268,728 |
| Genes with a single exon | 8,212 |
| Average gene length (bp) | 28,900 |
| Average CDS length (bp) | 1,711 |
| Average transcript length (bp) | 30,716 |
| Average exons per transcript | 7.3 |
| Average introns in CDS per transcript | 4.8 |
| % genome covered by genes | 42.00% |
| % genome covered by CDS | 1.60% |
| Longest gene (bp) | 2,100,374 |
| Longest CDS (bp) | 88,110 |
| Longest intron (bp) | 768,181 |

**Figure 3.**
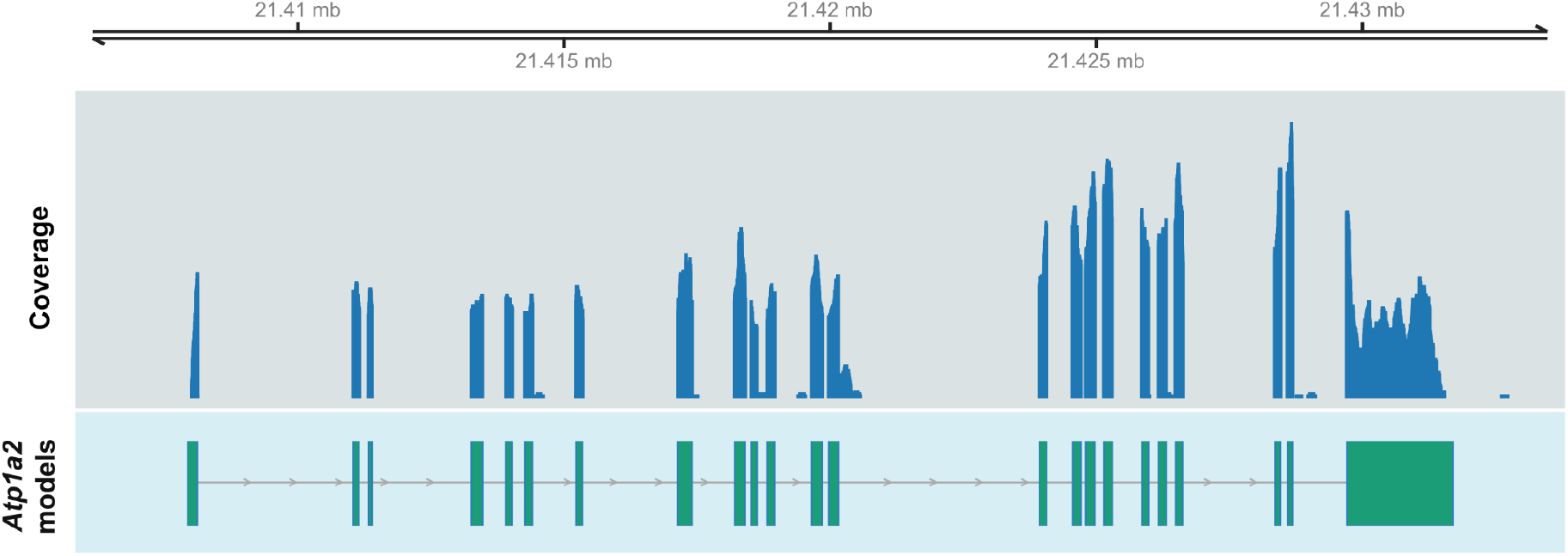
RNA-seq reads coverage across the *Atp1a2* gene in *A. olivacea*. The top panel shows the genomic region containing the *Atp1a2* locus, while the middle panel displays the RNA-seq read alignments. The bottom panel shows transcript models predicted from the RNA-seq data. Exon coverage and splice junctions are well supported, illustrating the completeness of the assembled genome and its utility for transcriptomic analyses.

### Dataset for ecotype associated targets

To assess the impact of using a complete reference genome on differential expression analyses, we revisited the data previously analyzed by Giorello et al. (2018), who employed a *de novo* transcriptome assembly for *A. olivacea* to assess differential expression profiles in forest and steppe samples. In our study, we used the newly assembled reference genome of the species for read mapping and quantification. Using the genome-based approach, we identified 985 differentially expressed genes (FDR < 0.05) between forest and steppe individuals, whereas the previous de novo transcriptome-based analysis reported 2,860 differentially expressed transcripts under the same significance threshold. Among these, 523 genes were shared between both approaches. The observed differences likely reflect the higher fragmentation and potential redundancy inherent to *de novo* assemblies, as well as the improved read assignment and gene annotation provided by the reference genome. These results highlight the importance of having a high-quality genome to optimize comparative transcriptomic analyses in non-model species (**Figure 4**). However, it is important to note that many genes were exclusively detected as differentially expressed in the previous analysis by Giorello et al. (2018). These may include lineage-specific or poorly annotated transcripts, isoform variants, or tissue-specific genes that are underrepresented or fragmented in the current genome annotation. Therefore, both strategies are complementary: the reference genome enhances confidence and interpretability, while transcriptome-based assemblies may still capture biologically relevant elements missed by automated annotation. Future annotation efforts integrating long-read sequencing and manual curation will be essential to reconcile these two views and fully capture the transcriptional complexity of *A. olivacea*.

**Figure 4.**
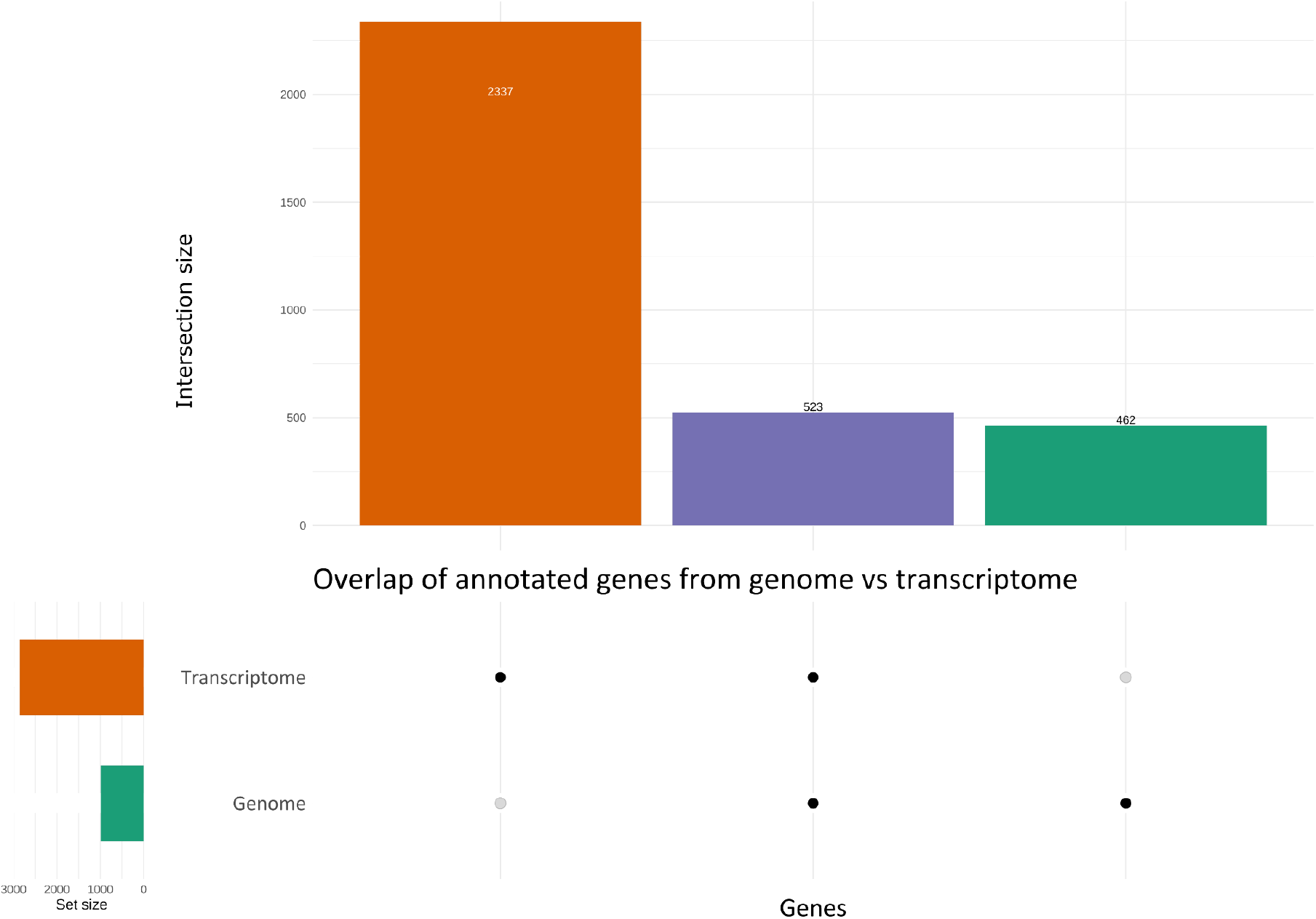
Overlap of annotated genes identified from genome (this study) and transcriptome analyses UpSet plot showing the distribution of gene symbols recovered from genome-based annotation (green) and transcriptome-based annotation (orange). The intersection (purple) highlights genes detected by both approaches, while the vertical bars indicate the size of each set or intersection. Unique sets correspond to genes identified exclusively by one of the approaches. This comparison highlights the complementarity of genome and transcriptome data for gene discovery.

To further investigate the sources and pathways associated with these DEGs, we conducted GSEA using the C5 ontology gene sets as references where 143 significant clusters were detected (**Table 4 and Table S1**). The analysis revealed that DEGs associated with the forest ecotype were significantly enriched in several immune-related processes, including *antigen processing and presentation, regulation of T cell mediated cytotoxicity*, and *response to interferon beta*. This suggests an upregulation of both innate and adaptive immune responses in forest-dwelling individuals, potentially reflecting increased exposure to pathogens in more humid environments or ecotype-specific immune strategies. In contrast, DEGs associated with the steppe ecotype (i.e., gene sets with negative enrichment scores) were predominantly enriched in developmental and metabolic processes such as *kidney morphogenesis, epithelial cell differentiation, lipid metabolism*, and *thermogenesis*. These functions are consistent with adaptive responses to arid and colder environments, where efficient water reabsorption and non-shivering thermogenesis may be particularly advantageous. Overall, the contrasting enrichment profiles between forest and steppe individuals underscore the ecological pressures shaping gene expression in each habitat, with the forest ecotype showing enhanced immune activity and the steppe ecotype exhibiting traits related to physiological adaptation to abiotic stress.

**Table 4.** Top 10 positively and negatively enriched gene sets identified by GSEA. To facilitate interpretation, only the 10 gene sets with the highest and lowest normalized enrichment scores (NES) are shown. Positive NES values indicate enrichment in genes upregulated in forest ecotype samples, while negative NES values correspond to genes upregulated in steppe ecotype samples. GSEA was performed using GO:BP terms from the C5 collection of MSigDB.

| Description | setSize | enrichmentScore | NES | pvalue | p.adjust |
| --- | --- | --- | --- | --- | --- |
| ANTIGEN PROCESSING AND PRESENTATION | 47 | 0.67 | 2.12 | 0 | 0.00 |
| ANTIGEN PROCESSING AND PRESENTATION OF PEPTIDE ANTIGEN | 25 | 0.70 | 1.92 | 0 | 0.02 |
| REGULATION OF T CELL MEDIATED CYTOTOXICITY | 16 | 0.77 | 1.91 | 0 | 0.03 |
| ANTIGEN PROCESSING AND PRESENTATION OF PEPTIDE ANTIGEN VIA MHC CLASS I | 16 | 0.76 | 1.86 | 0 | 0.03 |
| HORMONE BIOSYNTHETIC PROCESS | 23 | 0.69 | 1.85 | 0 | 0.03 |
| RESPONSE TO INTERFERON BETA | 16 | 0.75 | 1.84 | 0 | 0.04 |
| STEROID HORMONE BIOSYNTHETIC PROCESS | 20 | 0.70 | 1.83 | 0 | 0.05 |
| LEUKOCYTE MEDIATED CYTOTOXICITY | 51 | 0.57 | 1.82 | 0 | 0.04 |
| XENOBIOTIC METABOLIC PROCESS | 67 | 0.53 | 1.81 | 0 | 0.02 |
| LEUKOCYTE MEDIATED IMMUNITY | 163 | 0.46 | 1.81 | 0 | 0.01 |
| DETECTION OF ABIOTIC STIMULUS | 45 | -0.74 | -2.28 | 0 | 0.00 |
| DETECTION OF LIGHT STIMULUS | 17 | -0.86 | -2.21 | 0 | 0.00 |
| SENSORY PERCEPTION OF LIGHT STIMULUS | 71 | -0.65 | -2.17 | 0 | 0.00 |
| DETECTION OF STIMULUS | 71 | -0.64 | -2.16 | 0 | 0.00 |
| TRIGLYCERIDE METABOLIC PROCESS | 54 | -0.67 | -2.15 | 0 | 0.00 |
| SENSORY PERCEPTION | 191 | -0.53 | -2.11 | 0 | 0.00 |
| REGULATION OF PLASMA LIPOPROTEIN PARTICLE LEVELS | 44 | -0.68 | -2.10 | 0 | 0.00 |
| NEUTRAL LIPID BIOSYNTHETIC PROCESS | 31 | -0.72 | -2.08 | 0 | 0.01 |
| PROTEIN LIPID COMPLEX ASSEMBLY | 17 | -0.82 | -2.08 | 0 | 0.01 |
| REGULATION OF LIPID STORAGE | 27 | -0.72 | -2.04 | 0 | 0.01 |

### Comparison between A. olivacea and desert-adapted species DEGs

In order to compare our findings with those from another desert-adapted species, we conducted the DGE and GSEA on a dataset from the mouse *Peromyscus eremicus* exposed to experimental dehydration to examine the effects on desert-adapted animals (MacManes 2017). To reanalyze the *P. eremicus* dataset, we used the newly available reference genome instead of the *de novo* transcriptome assembly employed in the original study. This allowed us to assess how the choice of reference influences the detection of DEGs. A similar pattern to that observed in *A. olivacea* emerged: 455 DEGs were identified by both approaches, whereas 751 were uniquely detected using the *de novo* transcriptome and 395 were exclusive to the genome-guided analysis (Figure S2).

Strikingly, when comparing the DEGs identified between the natural ecotypes of *A. olivacea* and those obtained from the experimental dehydration treatments in *P. eremicus*, only 65 genes were shared (**Table 5**). This limited overlap suggests that the molecular responses distinguishing the forest and steppe ecotypes of *A. olivacea* likely cannot be explained solely by dehydration stress. Instead, these differences may reflect broader ecological, physiological, and evolutionary processes—including long-term adaptation to multiple abiotic factors such as temperature, food availability, and photoperiod, as well as biotic interactions. The contrasting DEG profiles underscore that environmental adaptation in natural populations may involve complex regulatory programs beyond those triggered by short-term experimental perturbations. Also, among those shared DEGs between *A. olivacea* ecotypes and dehydration-treated *P. eremicus*, we can highlight one of the most biologically meaningful gene; *Agt* that encodes angiotensinogen, the precursor of the renin–angiotensin–aldosterone system (RAAS). This pathway plays a central role in water and electrolyte balance, mediating renal reabsorption and blood pressure regulation. The consistent differential expression of *Agt* in steppe-dwelling *A. olivacea* and dehydrated *P. eremicus* highlights a convergent molecular signature of dehydration response. Whereas in *P. eremicus* this reflects acute transcriptomic plasticity to experimental stress, in *A. olivacea* it may represent a long-term evolutionary adjustment to xeric habitats. This parallel points to RAAS-related genes as core components of mammalian adaptation to water limitation. In addition to *Agt*, members of the solute carrier (Slc) family stand out due to their consistent expression patterns across both *A. olivacea* and *P. eremicus*. Specifically, *Slc15a2, Slc3a1*, and *Slc7a13* were upregulated in forest-dwelling *A. olivacea* and in hydrated *P. eremicus*, suggesting that their increased expression is associated with conditions of water availability. In contrast, *Slc2a1* showed the opposite pattern, being upregulated in steppe individuals and in dehydrated mice, pointing to a role in metabolic adjustments under water restriction, possibly by facilitating glucose uptake as an energy source under stress. These coherent cross-species signatures highlight solute transporters as critical components of renal and systemic responses to hydration status. Importantly, they illustrate how both acute dehydration stress and chronic adaptation to xeric habitats converge on similar molecular pathways, with Slc-mediated solute transport emerging as a central mechanism in mammalian water balance. Also, stress-related genes including *Fkbp5, Hspa2*, and *Dnaja1* indicate activation of cellular stress responses, while metabolic regulators (*Dgat2, Sqle, Rbp4*) point to adjustments in lipid metabolism that may contribute to water conservation through metabolic water production. Finally, genes associated with extracellular matrix and tissue remodeling (*Col15a1, Fermt1, Itgb6, Fst, Ptn*) suggest that structural changes in renal tissue could accompany physiological adaptation. Together, these shared DEGs reinforce the notion that, although overall transcriptomic overlap between *A. olivacea* and *P. eremicus* is limited, key pathways related to renal function, metabolism, and stress response are consistently recruited under xeric conditions.

**Table 5.**
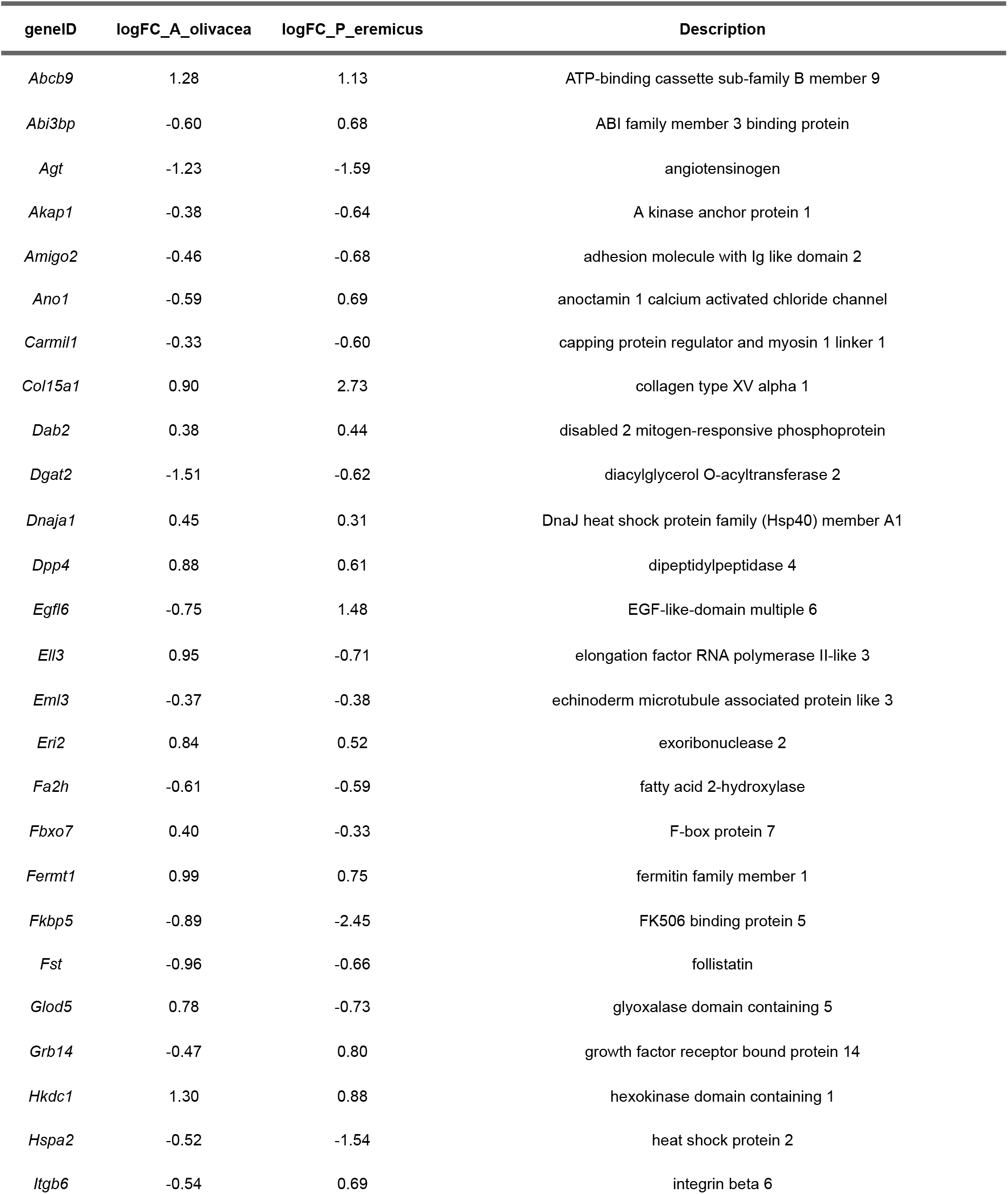
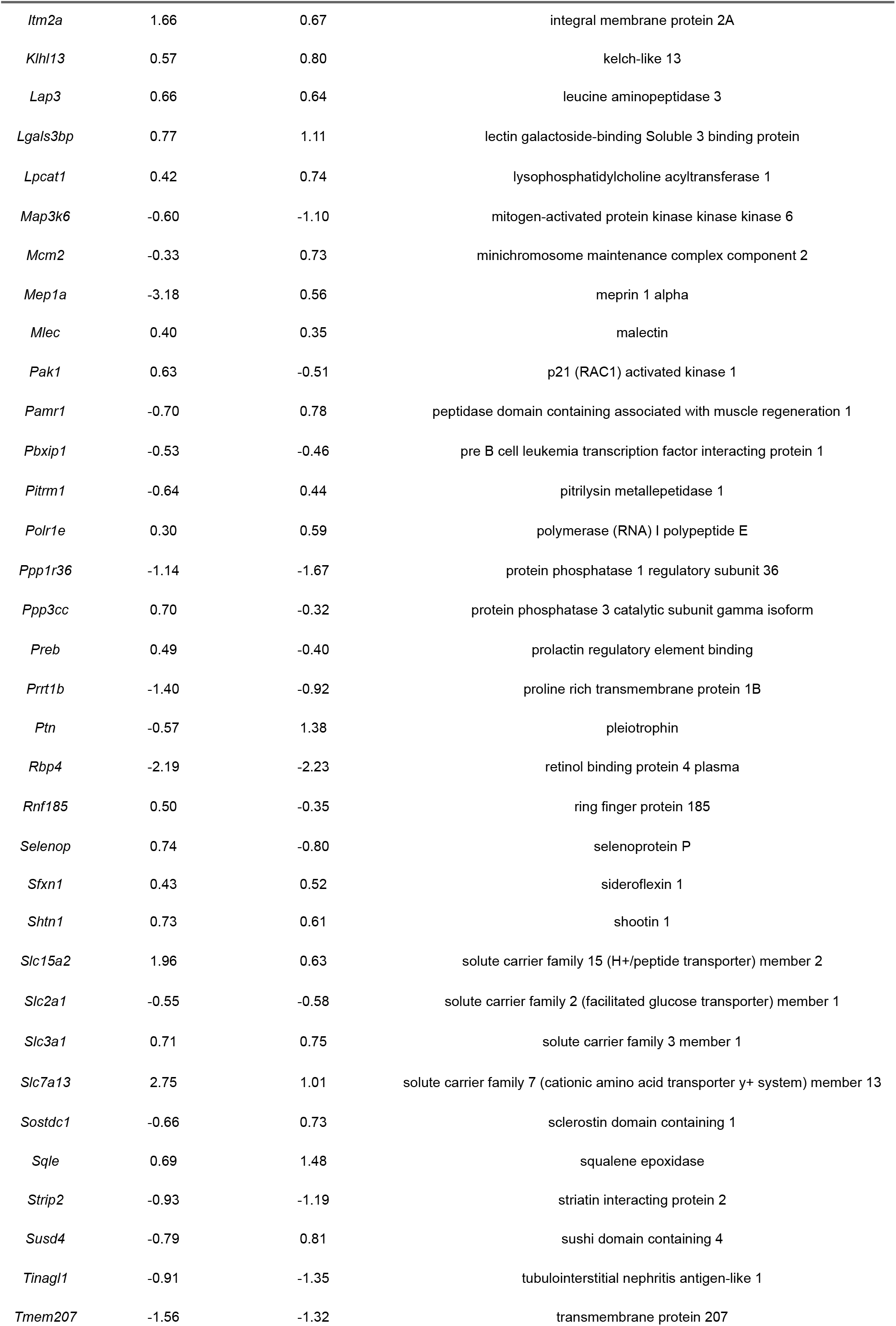
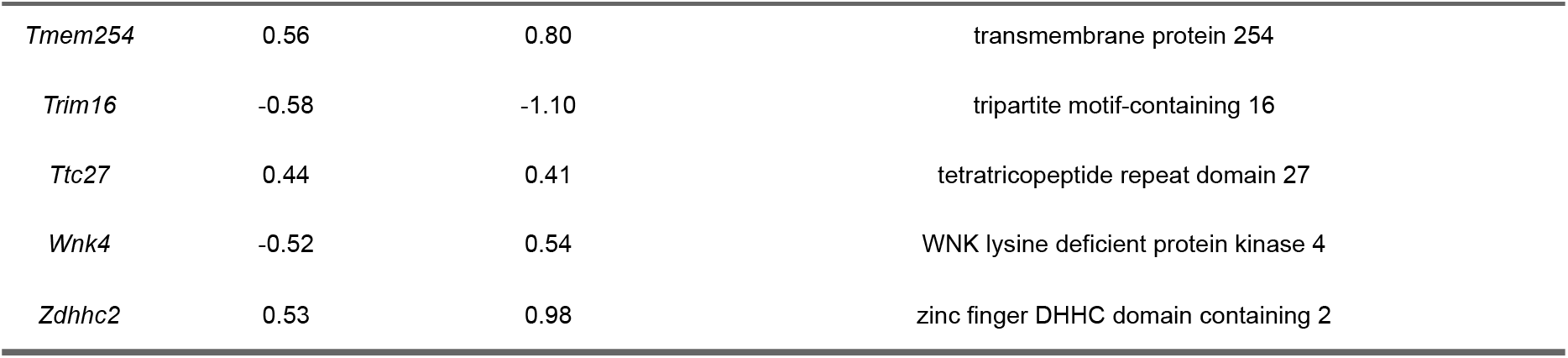
Differentially expressed genes (DEGs) shared between A. olivacea (forest vs. steppe ecotypes) and P. eremicus (hydrated vs. dehydrated treatments). For each gene, log fold change (logFC) values in both species are shown together with a brief functional description. Despite the limited overlap (65 DEGs), several candidates are functionally relevant for dehydration response and renal physiology, including Agt, solute carriers, stress-related proteins, and genes involved in tissue remodeling.

In addition to *P. eremicus*, data from the desert rodent *Jaculus jaculus* under experimental dehydration further reinforce the idea of partially convergent molecular strategies for coping with water limitation (Gillard et al. 2023). Several key genes overlapped with those identified in *A. olivacea*, including *Ren* (renin) and *Aqp2*, central components of the RAAS pathway and collecting duct water reabsorption, both upregulated under dehydration. Likewise, the glucose transporter *Slc2a1* showed consistent activation across species, highlighting its role in metabolic adjustments under limited water availability. Stress-related genes such as *Dnaja1*, together with solute transporters including *Slc5a2*, also exhibited dehydration-associated regulation, underscoring the importance of tubular transport and cellular stress responses. The recurrence of these genes across independent rodent lineages inhabiting xeric environments demonstrates that renal adaptation relies on a conserved molecular toolkit, where RAAS signaling, solute carrier activity, and stress response pathways repeatedly emerge as central mechanisms for maintaining water balance under arid conditions (Table S2).

### Deconvolution

A kidney cell type signature gene set was constructed and implemented to identify the source of DEGs at the cell type level from olive mouse bulk RNA-seq data across both ecotypes (**Figure 5**). The deconvolution analysis provided a complementary perspective on the renal transcriptomes of *A. olivacea*. As expected, epithelial cells from proximal tubules and the loop of Henle dominated the inferred cellular composition, reflecting their central roles in water and solute handling. Notably, forest individuals displayed higher relative contributions of proximal tubule cells, whereas steppe individuals showed increased signals from the loop of Henle and mesangial cells. Integrating deconvolution with differential expression analysis highlighted that the cellular basis of transcriptional divergence between forest and steppe ecotypes involves both renal and immune compartments. In the steppe, the overexpression of the collecting duct marker *Fxyd4* suggests a greater functional contribution of collecting duct epithelial cells, which may reflect adjustments in water and electrolyte handling under drier or more osmotically challenging conditions. In contrast, the forest exhibited higher expression of the proximal tubule marker *Enpep*, pointing to enhanced activity of proximal tubule epithelial cells, which play central roles in nutrient reabsorption and metabolic homeostasis. Additionally, the upregulation of macrophage-associated genes (*Csf1r* and *Cd68*) indicates an increased presence or activation of immune cells in the forest environment, potentially linked to differences in pathogen exposure or immune surveillance. Together, these results suggest that ecological differences between biomes shape kidney transcriptomes by modulating the relative abundance and activity of distinct renal and immune cell populations. These cellular shifts parallel the observed transcriptional differences in solute carriers and metabolic genes, underscoring that ecotypic divergence involves not only gene-level changes but also the engagement of distinct cellular programs within the kidney. Such variation may underlie differences in urine concentrating ability and water conservation strategies, highlighting the multi-layered nature of adaptation to extreme environments.

**Figure 5.**
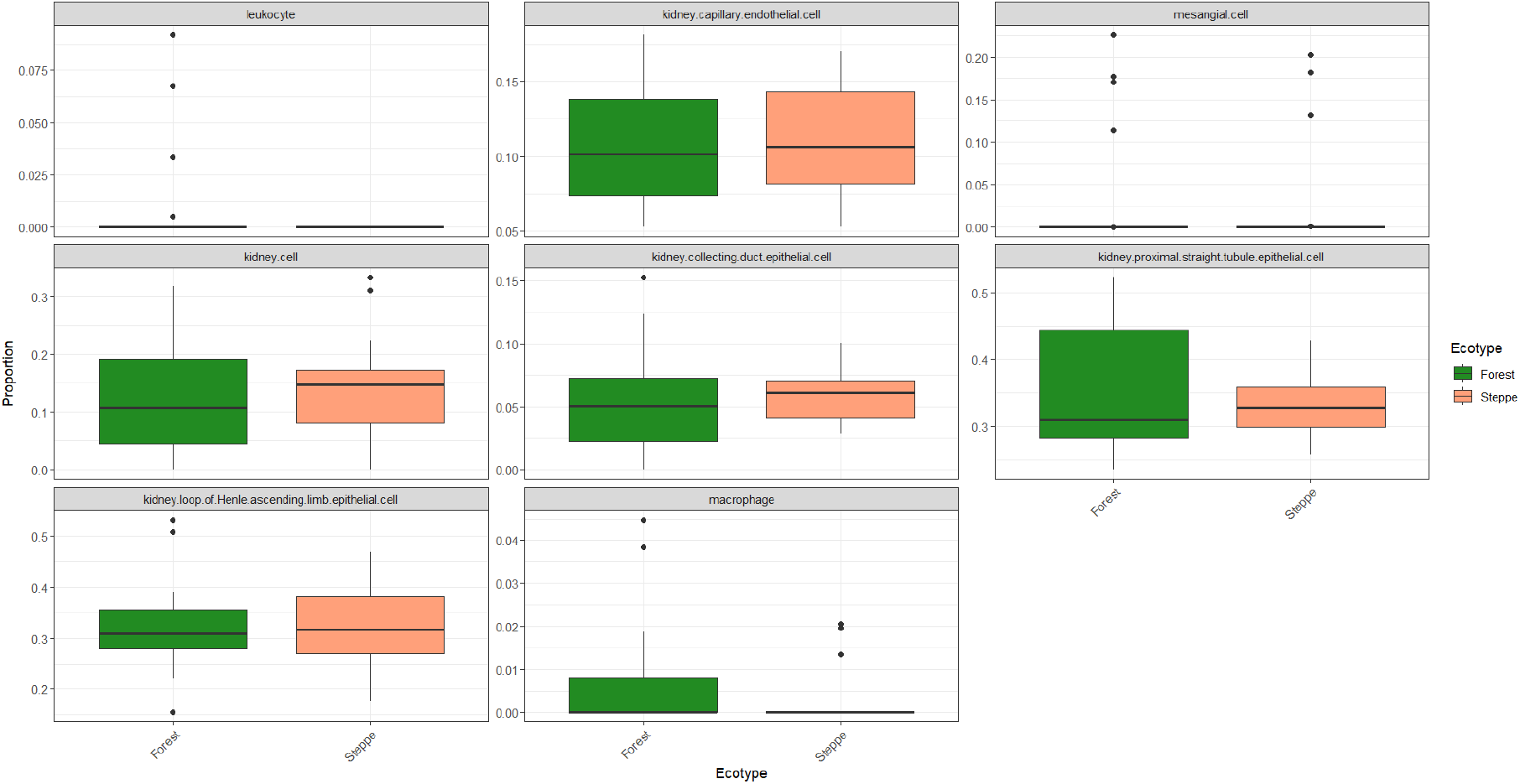
Cell type deconvolution of kidney transcriptomes from *A. olivacea* using CIBERSORTx. Stacked barplots show the relative proportions of inferred renal cell types in individuals from forest and steppe ecotypes. Major contributions were observed from the proximal tubule and loop of Henle epithelial cells, with differences between ecotypes suggesting variation in cell-type representation linked to water availability. These results integrate with differential expression findings, indicating that ecotypic divergence involves both transcriptional and cellular-level changes in renal physiology.

## Conclusion

This study presents the first high-quality genome assembly of the sigmodontine tribe Abrotrichini, the one that includes *Abrothrix olivacea*. This species thrives across contrasting biomes of southern South America. The newly generated genome represents a major resource for evolutionary and ecological genomics in sigmodontine rodents, providing a platform for studies of adaptation, physiology, and conservation. Beyond its potential value, we illustrated its utility through a case study of kidney transcriptomes of *A. olivacea* inhabiting forests and steppes.

Our analyses revealed striking ecological contrasts. Forest-dwelling individuals displayed strong enrichment for immune-related pathways, with upregulation of genes linked to macrophages, natural killer cells, and fibroblasts. Renal macrophages are known to play pivotal roles in immune surveillance and tissue homeostasis (Bell and Conway 2022), and their increased signature in forest animals may reflect ecological pressures such as higher pathogen exposure in humid environments. Interestingly, experimental models of salt-sensitive hypertension in rodents demonstrate that dietary protein source modulates kidney injury through immune–microbiota interactions (Mattson et al. 2023). As such, the forest-associated immune activation in *A. olivacea* could represent not pathology but an adaptive adjustment, potentially shaped by dietary and environmental inputs. Parallels with human studies are also striking; even brief exposure to forest environments enhances innate immune activity, increasing NK cell function and reducing stress hormones (Li et al. 2010). Together, these findings support the notion that forest habitats prime immunological states in mammals, a pattern now observed both in humans and in wild rodents.

Conversely, steppe individuals showed enrichment for developmental and metabolic pathways, including processes related to kidney morphogenesis, epithelial differentiation, lipid metabolism, and thermogenesis—traits consistent with adaptation to arid, cold environments where efficient water reabsorption and energy balance are critical. These results align with convergent responses observed in desert rodents such as *Peromyscus eremicus*, in which RAAS-related genes and solute carriers underlie responses to water stress.

Finally, evidence from other mammalian systems underscores the broader eco-physiological framework. In wild woodrats, populations feeding on toxic creosote bush host a unique functional microbiome adapted for detoxification (Stapleton et al. 2024). Analogously, environmental differences between forest and steppe may shape not only the renal transcriptome of *A. olivacea* but also its gut microbiome, selecting microbial consortia with metabolic capacities that support water and osmotic balance. Considering the interplay of microbiome function, immune cell activity, and renal gene expression highlights the complexity of adaptation in natural populations.

In sum, the genome of *A. olivacea* not only fills a critical gap in the genomic resources for sigmodontine rodents but also enables the dissection of multilevel adaptive processes, from gene regulation to cell-type composition and host–microbiome interactions. These findings exemplify how high-quality genomic resources in non-model species can illuminate the molecular architecture of adaptation to heterogeneous environments.

