## Supplemental files for "Genome assembly and annotation of the olive grass mouse *Abrothrix olivacea* reveal transcriptomic and cellular adaptations across contrasting biomes"

**Table S1. Complete results of Gene Set Enrichment Analysis (GSEA) contrasting kidney transcriptomes between forest and steppe ecotypes of *A. olivacea*.**
**The analysis was performed using the MSigDB C5 collection (GO: Biological Processes).**
**For each gene set, enrichment statistics are reported, including set size, enrichment score (ES), normalized enrichment score (NES), and associated significance values.**
**Positive NES values indicate enrichment in forest samples, while negative NES values correspond to enrichment in steppe samples.**

| ID | setSize | enrichmentScor | NES | pvalue | p.adjust | qvalue | rank | core_enrichmer | Ecotype |
|---|---|---|---|---|---|---|---|---|---|
| ANTIGEN PROCESSING AND PRESENTATION | 45 | 0.7047833259 | 2.311200454 | 1.73E-06 | 0.000882979876 | 0.000785490194 | 1665 | Wdfy4/Abcb9/Mp | \| |
| REGULATION OF T CELL MEDIATED CYTOTOXICIT | 16 | 0.8034987413 | 2.068726648 | 0.000202359901 | 0.01868950751 | 0.01662600165 | 933 | Ptprc/Pirb/Slc22a | Forest |
| ANTIGEN PROCESSING AND PRESENTATION OF I | 24 | 0.7149645942 | 2.001281385 | 0.000481851459 | 0.02783809874 | 0.02476449822 | 1665 | Abcb9/Mpeg1/Fc | Forest |
| STEROID HORMONE BIOSYNTHETIC PROCESS | 18 | 0.7446785544 | 1.948937373 | 0.000896661538 | 0.0401372969 | 0.03570574366 | 909 | Hsd17b1/Hsd3b1 | Forest |
| ANTIGEN PROCESSING AND PRESENTATION OF I | 15 | 0.7753212777 | 1.945242292 | 0.000712732787 | 0.03444572103 | 0.03064257388 | 1657 | Abcb9/Mpeg1/Fc | Forest |
| LEUKOCYTE MEDIATED CYTOTOXICITY | 47 | 0.5862565215 | 1.925566159 | 0.000810624895 | 0.03723558944 | 0.03312441332 | 954 | Ighg3/Ptprc/Pirb/ | Forest |
| REGULATION OF ANTIGEN PROCESSING AND PR | 7 | 0.9312042089 | 1.915190521 | 0.000184515654 | 0.01782802982 | 0.01585963958 | 362 | Ccl21a/Pirb/Cd68 | Forest |
| VITAMIN K METABOLIC PROCESS | 8 | 0.898006595 | 1.883605405 | 0.000291392762 | 0.02177286718 | 0.01936892801 | 419 | Cyp4f40/Cyp4f14 | Forest |
| DENDRITIC CELL ANTIGEN PROCESSING AND PR | 6 | 0.9544514276 | 1.869374863 | 4.36E-05 | 0.006426000319 | 0.005716506539 | 362 | Mpeg1/Ccl21a/C | Forest |
| LEUKOCYTE MEDIATED IMMUNITY | 156 | 0.4741322049 | 1.86530199 | 1.72E-05 | 0.003847671633 | 0.003422850756 | 1004 | Ptgds/Ighg3/S10 | Forest |
| NEGATIVE REGULATION OF CELL KILLING | 11 | 0.8202331401 | 1.864874709 | 0.001231957855 | 0.04664791772 | 0.04149752776 | 804 | Ptprc/Pirb/Fcgr2b | Forest |
| LYMPHOCYTE MEDIATED IMMUNITY | 114 | 0.4881996329 | 1.836170786 | 9.82E-05 | 0.01196050025 | 0.01063994312 | 970 | Ighg3/C1qc/Ptprc | Forest |
| NEGATIVE REGULATION OF INTERLEUKIN 10 PRC | 8 | 0.87140391 | 1.82780519 | 0.00104313401 | 0.04304255467 | 0.03829023234 | 667 | Pirb/Fcgr2b/Cd27 | Forest |
| XENOBIOTIC METABOLIC PROCESS | 60 | 0.5381661261 | 1.825664689 | 0.001327201369 | 0.04893181888 | 0.04352926372 | 725 | Gstm7/Ugt2b1/SI | Forest |
| DEFENSE RESPONSE TO VIRUS | 147 | 0.4618390505 | 1.81242241 | 0.000239275004 | 0.02060725116 | 0.01833200749 | 1531 | Rnase1/Gbp7/Sp | Forest |
| REGULATION OF DENDRITIC CELL ANTIGEN PRO | 5 | 0.9495802498 | 1.790848052 | 0.000445401138 | 0.0272254138 | 0.02421945974 | 362 | Ccl21a/Cd68/Fgl | Forest |
| NEGATIVE REGULATION OF DENDRITIC CELL DIF | 5 | 0.9489504675 | 1.789660322 | 0.000454353850 | 0.0272254138 | 0.02421945974 | 456 | Pirb/Fcgr2b/Tme | Forest |
| TYPE I INTERFERON PRODUCTION | 67 | 0.5180235887 | 1.780130339 | 0.00089412657 | 0.0401372969 | 0.03570574366 | 1437 | Gbp7/Isg15/Irgm | Forest |
| PROTEIN FOLDING | 140 | 0.4549310076 | 1.759298648 | 0.000288461421 | 0.02177286718 | 0.01936892801 | 2912 | Sdf2l1/Dnaja4/Ly | Forest |
| CELL ACTIVATION INVOLVED IN IMMUNE RESPON | 140 | 0.4519263286 | 1.747679023 | 0.000383818667 | 0.02591469655 | 0.02305345859 | 1004 | Sucnr1/Itm2a/Ptg | Forest |
| ADAPTIVE IMMUNE RESPONSE BASED ON SOMA | 131 | 0.4470376545 | 1.71945348 | 0.000650973468 | 0.03441750783 | 0.03061747569 | 970 | Ighg3/C1qc/Ptprc | Forest |
| CYTOPLASMIC TRANSLATION | 114 | 0.4489053827 | 1.688380928 | 0.001157583893 | 0.04548018328 | 0.04045872272 | 3812 | Rps20/Mcts1/Rp | Forest |
| IMMUNE EFFECTOR PROCESS | 268 | 0.3990570583 | 1.672141093 | 0.000127452045 | 0.01323351029 | 0.0117724003 | 1004 | Sucnr1/Itm2a/Ptg | Forest |
| RESPONSE TO VIRUS | 202 | 0.3925417421 | 1.60088809 | 0.000570188764 | 0.03195337838 | 0.02842541042 | 1720 | Rnase1/Gbp7/Sp | Forest |
| CYTOKINE PRODUCTION | 337 | 0.3497116823 | 1.506398872 | 0.000736490546 | 0.03497705952 | 0.03111524736 | 1754 | C1qtnf3/S100a13 | Forest |
| DEFENSE RESPONSE TO OTHER ORGANISM | 490 | 0.3362311422 | 1.494258306 | 0.000273410788 | 0.02128047302 | 0.01893089902 | 1437 | Rnase1/Slc15a2/ | Forest |
| POSITIVE REGULATION OF CELL DIFFERENTIATIC | 431 | -0.3306336671 | -1.43196341 | 0.001044573061 | 0.04304255467 | 0.03829023234 | 1921 | Gata3/Ripk1/Fzd | Steppe |
| CELL CELL ADHESION | 404 | -0.3324839245 | -1.432336597 | 0.001079103629 | 0.04414085208 | 0.03926726689 | 2520 | Itga6/Ppm1f/Ep3 | Steppe |
| GROWTH | 486 | -0.3306712729 | -1.446369009 | 0.001023452974 | 0.04296917378 | 0.03822495342 | 2678 | Zfyve27/Scnn1b/ | Steppe |
| ACTIN FILAMENT BASED PROCESS | 469 | -0.3382172522 | -1.479305625 | 0.000264639235 | 0.02118626109 | 0.01884708902 | 2957 | Kcnq1/Akap11/P | Steppe |
| HEAD DEVELOPMENT | 393 | -0.3456750959 | -1.479345348 | 0.001021585863 | 0.04296917378 | 0.03822495342 | 2568 | Atm/Setd1a/Map | Steppe |
| DEVELOPMENTAL GROWTH | 343 | -0.3634993606 | -1.537506945 | 0.000282597735 | 0.02169421522 | 0.01929896001 | 1524 | Cttn/Dag1/Smo/S | Steppe |
| SYNAPTIC SIGNALING | 288 | -0.3697333015 | -1.538461699 | 0.000627530058 | 0.03414250918 | 0.0303728396 | 2196 | Plp1/Cpeb3/Snca | Steppe |
| TISSUE MORPHOGENESIS | 330 | -0.3691293772 | -1.561062257 | 0.000362524986 | 0.0250813583 | 0.0223121291 | 2614 | Clasp1/Frs2/Hox | Steppe |
| CELL PROJECTION MORPHOGENESIS | 335 | -0.3695221783 | -1.561388655 | 0.000305932377 | 0.02255848744 | 0.02006780806 | 1921 | Gata3/Fzd3/Dvl1 | Steppe |
| MORPHOGENESIS OF AN EPITHELIUM | 280 | -0.3765469703 | -1.568299479 | 0.000415158260 | 0.02643803287 | 0.02351901342 | 2737 | Clasp1/Frs2/Hox | Steppe |
| MUSCLE SYSTEM PROCESS | 191 | -0.3981459945 | -1.582193546 | 0.001091436242 | 0.04432180216 | 0.03942823829 | 1524 | Cttn/Dag1/Camta | Steppe |
| BEHAVIOR | 263 | -0.3848075978 | -1.588271836 | 0.000255324319 | 0.02104172775 | 0.01871851359 | 1445 | Thra/Per3/Stat3/ | Steppe |
| BLOOD VESSEL MORPHOGENESIS | 340 | -0.379658157 | -1.599903382 | 0.000110495012 | 0.01238428101 | 0.01101693431 | 1734 | Klf4/Sulf1/Fzd8/C | Steppe |
| REGULATION OF HORMONE LEVELS | 236 | -0.3947068044 | -1.606632401 | 0.000310082623 | 0.0225675717 | 0.02007588932 | 796 | Adcy5/Hsd17b11 | Steppe |
| REGULATION OF SMALL GTPASE MEDIATED SIGN | 208 | -0.4028477417 | -1.618082252 | 0.000531736728 | 0.03018359226 | 0.02685102614 | 2614 | Tgfb2/Itpkb/Rabg | Steppe |
| REGULATION OF VASCULATURE DEVELOPMENT | 156 | -0.4175238836 | -1.620130675 | 0.001241505601 | 0.0466939422 | 0.04153847069 | 1832 | Tgfb2/Nos3/Sem | Steppe |
| HEART DEVELOPMENT | 320 | -0.3867730622 | -1.622495103 | 8.64E-05 | 0.01125913022 | 0.01001601126 | 2614 | Tgfb2/Nos3/Atm/ | Steppe |
| REGULATION OF EPITHELIAL CELL MIGRATION | 159 | -0.4170600224 | -1.628528604 | 0.001301214177 | 0.04829141886 | 0.04295957018 | 2614 | Tgfb2/Nos3/Bcas | Steppe |
| VASCULATURE DEVELOPMENT | 408 | -0.3783851496 | -1.632165013 | 1.49E-05 | 0.003562795079 | 0.003169427381 | 2463 | Aggf1/Ap2b1/Ptk | Steppe |
| MESENCHYME DEVELOPMENT | 172 | -0.414940402 | -1.6342416 | 0.000450998446 | 0.0272254138 | 0.02421945974 | 2614 | Tgfb2/Nos3/Map | Steppe |
| AMIDE TRANSPORT | 143 | -0.4313461469 | -1.65150989 | 0.000721866358 | 0.03457554765 | 0.03075806636 | 1057 | Irs2/Abcc1/Abca | Steppe |
| DEVELOPMENTAL GROWTH INVOLVED IN MORPH | 132 | -0.4373327695 | -1.652019229 | 0.000769139046 | 0.03591879349 | 0.03195300461 | 1689 | Slc9a6/Lzts2/Yap | Steppe |
| REGULATION OF ANATOMICAL STRUCTURE MOR | 452 | -0.3813905614 | -1.655349571 | 3.47E-06 | 0.001295957834 | 0.001152871314 | 1846 | Sdc2/Pak4/Inf2/L | Steppe |
| CELL MORPHOGENESIS INVOLVED IN NEURON D | 290 | -0.3997732736 | -1.664878585 | 0.000124873570 | 0.01323351029 | 0.0117724003 | 1921 | Gata3/Fzd3/Dvl1 | Steppe |
| HEART MORPHOGENESIS | 132 | -0.440854233 | -1.665321515 | 0.000635763974 | 0.03425789725 | 0.0304754877 | 2260 | Cldn5/Ccm2l/Asx | Steppe |
| ORGANIC HYDROXY COMPOUND TRANSPORT | 117 | -0.450009273 | -1.676784742 | 0.000713009214 | 0.03444572103 | 0.03064257388 | 1207 | Ldlrap1/Scarb1/S | Steppe |
| EPITHELIAL TUBE MORPHOGENESIS | 195 | -0.4248700454 | -1.692664367 | 0.000171453793 | 0.01685661505 | 0.01499547858 | 2265 | Tie1/Plxnb2/Ptch | Steppe |
| TISSUE MIGRATION | 207 | -0.4213545231 | -1.693401037 | 0.000260189096 | 0.02113187964 | 0.01879871182 | 2614 | Tgfb2/Nos3/Bcas | Steppe |
| LIPID LOCALIZATION | 234 | -0.4161979943 | -1.696552855 | 0.000107917250 | 0.01234220961 | 0.01097950801 | 1207 | Ldlrap1/Scarb1/V | Steppe |
| RESPIRATORY SYSTEM DEVELOPMENT | 131 | -0.4495696484 | -1.699176913 | 0.000651009248 | 0.03441750783 | 0.03061747569 | 2597 | Nos3/Mapk3/Fgf | Steppe |
| TUBE MORPHOGENESIS | 478 | -0.3891615228 | -1.699746718 | 5.51E-07 | 0.00035563534 | 0.000316369692 | 2463 | Aggf1/Ddr1/Ptk7 | Steppe |
| EPITHELIAL CELL DIFFERENTIATION | 362 | -0.4010398585 | -1.705557512 | 8.88E-06 | 0.002370623338 | 0.002108883153 | 1982 | Spry2/Jag1/Gstm | Steppe |
| ENDOTHELIAL CELL MIGRATION | 146 | -0.4444975978 | -1.708657053 | 0.000402168191 | 0.02626402833 | 0.0233642207 | 2415 | Klf4/Pik3r3/Card | Steppe |
| CELL CELL ADHESION VIA PLASMA MEMBRANE A | 95 | -0.4754465219 | -1.718481119 | 0.001166034266 | 0.04548018328 | 0.04045872272 | 2485 | Dchs1/Ptk7/Ptpn | Steppe |
| EPITHELIAL CELL DEVELOPMENT | 140 | -0.4496443513 | -1.724667793 | 0.000407739197 | 0.02626402833 | 0.0233642207 | 1959 | Vezf1/Ezr/Cldn5/ | Steppe |
| REGULATION OF PROTEIN SECRETION | 122 | -0.4591120586 | -1.728747763 | 0.000684659444 | 0.03444572103 | 0.03064257388 | 2124 | Rhbdf1/Golph3/N | Steppe |
| CELL CELL SIGNALING | 485 | -0.3963972491 | -1.732857598 | 2.95E-07 | 0.000236531565 | 0.000210416149 | 2004 | Nr1d1/Efnb1/Git | Steppe |
| HORMONE TRANSPORT | 144 | -0.4541268965 | -1.741295813 | 0.000127771989 | 0.01323351029 | 0.0117724003 | 949 | Gabbr1/Irs1/Adcy | Steppe |
| REGULATION OF SYSTEM PROCESS | 220 | -0.432110738 | -1.743562721 | 4.11E-05 | 0.006221203009 | 0.005534320871 | 1524 | Cttn/Dag1/Thra/ | Steppe |
| CELLULAR RESPONSE TO COLD | 5 | -0.9352141838 | -1.747196374 | 0.000907688003 | 0.0401372969 | 0.03570574366 | 5 | Prkaca/Ucp1 | Steppe |
| REGULATION OF INSULIN SECRETION | 75 | -0.5059643393 | -1.748484861 | 0.000657237807 | 0.03442206235 | 0.03062152735 | 2004 | Nr1d1/Slc25a22/ | Steppe |
| CARDIAC CELL FATE COMMITMENT | 5 | -0.9407139979 | -1.757471299 | 0.000612807733 | 0.03392700828 | 0.03018113213 | 384 | Tbx3/Tenm4/Sox | Steppe |
| RENAL SYSTEM DEVELOPMENT | 211 | -0.4383236597 | -1.760131346 | 2.48E-05 | 0.004628294867 | 0.00411728549 | 2544 | Tbc1d32/Sfrp1/F | Steppe |
| RESPONSE TO LIGHT STIMULUS | 169 | -0.4479626226 | -1.763967815 | 0.000105735864 | 0.01234220961 | 0.01097950801 | 1307 | Egfr/Col6a2/Map | Steppe |
| EMBRYONIC MORPHOGENESIS | 323 | -0.4189270616 | -1.764995356 | 2.15E-06 | 0.001001866384 | 0.000891250459 | 2614 | Tgfb2/Cobl/Hoxb | Steppe |
| PATTERN SPECIFICATION PROCESS | 194 | -0.4424650357 | -1.765910124 | 5.97E-05 | 0.008164664566 | 0.007263205114 | 2730 | Celsr2/Frs2/Hoxt | Steppe |
| EMBRYONIC ORGAN DEVELOPMENT | 222 | -0.437671913 | -1.7694523 | 1.33E-05 | 0.003397663678 | 0.003022528115 | 1921 | Gata3/Fzd3/Esrrl | Steppe |
| ANIMAL ORGAN MORPHOGENESIS | 494 | -0.404459343 | -1.770772639 | 2.27E-08 | 6.37E-05 | 5.66E-05 | 2597 | Nos3/Slc40a1/Hc | Steppe |
| TEMPERATURE HOMEOSTASIS | 101 | -0.486742421 | -1.770780537 | 0.000692369119 | 0.03444572103 | 0.03064257388 | 2004 | Nr1d1/Tlr4/Flcn/ | Steppe |
| AMEBOIDAL TYPE CELL MIGRATION | 277 | -0.4245607747 | -1.771837806 | 6.62E-06 | 0.001953727761 | 0.001738016957 | 2614 | Tgfb2/Nos3/Bcas | Steppe |
| POSITIVE REGULATION OF COA TRANSFERASE A | 5 | -0.9534027896 | -1.781176896 | 0.00020872915 | 0.01868950751 | 0.01662600165 | 92 | Agt/Apoa1 | Steppe |
| REGULATION OF COA TRANSFERASE ACTIVITY | 5 | -0.9534027896 | -1.781176896 | 0.00020872915 | 0.01868950751 | 0.01662600165 | 92 | Agt/Apoa1 | Steppe |
| BRANCHING MORPHOGENESIS OF AN EPITHELIA | 87 | -0.5024498895 | -1.78170546 | 0.000705522017 | 0.03444572103 | 0.03064257388 | 2265 | Tie1/Ptch1/Dlg5/ | Steppe |
| STRIATED MUSCLE TISSUE DEVELOPMENT | 121 | -0.4771029457 | -1.790572565 | 0.000242697818 | 0.02060725116 | 0.01833200749 | 2170 | Mtor/Ski/Mbd1/N | Steppe |
| NEPHRON EPITHELIUM DEVELOPMENT | 71 | -0.5208847825 | -1.792110852 | 0.001196211984 | 0.04621878072 | 0.04111577172 | 2247 | Asxl1/Ptch1/Cd2 | Steppe |

| | | | | | | | | | |
|---|---|---|---|---|---|---|---|---|---|
| NEPHRON DEVELOPMENT | 95 | -0.4983534795 | -1.801277337 | 0.000403227738 | 0.02626402833 | 0.0233642207 | 2485 | Dchs1/Plce1/Sul | Steppe |
| RENAL TUBULE DEVELOPMENT | 59 | -0.5460211693 | -1.80934554 | 0.001123883971 | 0.04531112069 | 0.04030832631 | 2213 | Ptch1/Cd24a/Hn | Steppe |
| UREA TRANSPORT | 6 | -0.916428317 | -1.811343177 | 0.00129507963 | 0.04829141886 | 0.04295957018 | 525 | Aqp3/Upk3a/Slc | Steppe |
| EPIDERMAL CELL DIFFERENTIATION | 92 | -0.5064644001 | -1.819872755 | 0.000223852982 | 0.01960112681 | 0.01743696919 | 1125 | Bcr/Hoxa7/Esrp1 | Steppe |
| KIDNEY MORPHOGENESIS | 57 | -0.5530170319 | -1.820318805 | 0.000909606835 | 0.0401372969 | 0.03570574366 | 2247 | Asxl1/Ptch1/Hnf1 | Steppe |
| KIDNEY EPITHELIUM DEVELOPMENT | 97 | -0.5024501877 | -1.821296588 | 0.000459723654 | 0.0272254138 | 0.02421945974 | 2247 | Asxl1/Ptch1/Cd2 | Steppe |
| REGULATION OF MIRNA METABOLIC PROCESS | 53 | -0.5626969397 | -1.8272972 | 0.001354572487 | 0.04961453737 | 0.04413660336 | 2663 | Esr1/Tgfb2/Drosl | Steppe |
| EMBRYONIC ORGAN MORPHOGENESIS | 137 | -0.4830448296 | -1.84024334 | 0.000127771989 | 0.01323351029 | 0.0117724003 | 2097 | Hnf1b/Pkd2/Spry | Steppe |
| EPITHELIAL CELL PROLIFERATION | 232 | -0.453614426 | -1.844083457 | 3.25E-06 | 0.0012959578340 | 0.001152871314 | 2004 | Nr1d1/Jag1/Cflar | Steppe |
| VENTRICULAR SEPTUM DEVELOPMENT | 48 | -0.5790956213 | -1.845725177 | 0.001025864982 | 0.04296917378 | 0.03822495342 | 2724 | Heg1/Sall1/Frs2/ | Steppe |
| SENSORY ORGAN MORPHOGENESIS | 118 | -0.4930162766 | -1.849140049 | 9.27E-05 | 0.01154964337 | 0.01027444889 | 1982 | Spry2/Jag1/Gata | Steppe |
| BROWN FAT CELL DIFFERENTIATION | 30 | -0.6385063521 | -1.855851211 | 0.001168655673 | 0.04548018328 | 0.04045872272 | 1396 | Ppargc1a/Cebpb | Steppe |
| REGULATION OF HORMONE SECRETION | 109 | -0.5036726543 | -1.858705682 | 0.000268465488 | 0.0211898676 | 0.01885029734 | 1307 | Egfr/Tfap2b/Pax | Steppe |
| VERY LOW DENSITY LIPOPROTEIN PARTICLE AS | 7 | -0.916108744 | -1.859856885 | 0.000450670779 | 0.0272254138 | 0.02421945974 | 130 | Soat2/Mfsd2a/Ap | Steppe |
| ISOPRENOID METABOLIC PROCESS | 64 | -0.551506236 | -1.861406165 | 0.000469568971 | 0.02741108868 | 0.02438463429 | 370 | Rdh10/Cyp1b1/F | Steppe |
| KERATINOCYTE DIFFERENTIATION | 63 | -0.5533673695 | -1.863206566 | 0.000783866403 | 0.03630402748 | 0.03229570497 | 1959 | Jag1/Cflar/Dsp/Y | Steppe |
| RESPONSE TO RETINOIC ACID | 53 | -0.5759067438 | -1.870194604 | 0.000745369932 | 0.03510128656 | 0.03122575851 | 2942 | Ptk2b/Gsk3b/Ph | Steppe |
| REGULATION OF BRANCHING INVOLVED IN URET | 10 | -0.8330643462 | -1.870345185 | 0.000533221919 | 0.03018359226 | 0.02685102614 | 1789 | Lhx1/Smo/Vegfa | Steppe |
| EMBRYONIC SKELETAL SYSTEM DEVELOPMENT | 64 | -0.5558720582 | -1.87614139 | 0.000376030819 | 0.02569849646 | 0.02286112913 | 1871 | Etl4/Ift140/Lhx1/ | Steppe |
| REGULATION OF MIRNA TRANSCRIPTION | 44 | -0.6019582604 | -1.879283486 | 0.000707171685 | 0.03444572103 | 0.03064257388 | 1998 | Smad6/Gata3/M | Steppe |
| PHOTOTRANSDUCTION | 9 | -0.8572940665 | -1.882520192 | 0.000934965740 | 0.04061665125 | 0.03613217257 | 6 | Plekhb1/Abca4/C | Steppe |
| NEGATIVE REGULATION OF CARTILAGE DEVELO | 10 | -0.8397539843 | -1.885364352 | 0.000461530034 | 0.0272254138 | 0.02421945974 | 1357 | Ltbp3/Rarg/Ccn4 | Steppe |
| SKIN DEVELOPMENT | 141 | -0.4919075507 | -1.888221583 | 2.10E-05 | 0.004349369495 | 0.003869156228 | 1959 | Jag1/Cflar/Ldb1/ | Steppe |
| NEGATIVE REGULATION OF CHONDROCYTE DIFF | 9 | -0.8603384265 | -1.88920526 | 0.000880681432 | 0.04012470526 | 0.03569454226 | 1357 | Ltbp3/Rarg/Ccn4 | Steppe |
| CELL MIGRATION INVOLVED IN SPROUTING ANGI | 35 | -0.62746418 | -1.892500964 | 0.001210041024 | 0.04621878072 | 0.04111577172 | 1298 | Klf4/Pik3r3/Card | Steppe |
| REGULATION OF EPITHELIAL CELL PROLIFERATI | 189 | -0.4753224467 | -1.89262195 | 4.31E-06 | 0.001510176754 | 0.001343438353 | 2004 | Nr1d1/Jag1/Cflar | Steppe |
| LENS DEVELOPMENT IN CAMERA TYPE EYE | 40 | -0.6195917706 | -1.898007967 | 0.000669201124 | 0.03444572103 | 0.03064257388 | 2169 | Ski/Fzr1/Vim/Spr | Steppe |
| LIPID STORAGE | 53 | -0.5850393711 | -1.899851819 | 0.000450797264 | 0.0272254138 | 0.02421945974 | 967 | Scarb1/Abca1/Fi | Steppe |
| PLASMA LIPOPROTEIN PARTICLE CLEARANCE | 24 | -0.6878653472 | -1.902833116 | 0.000971273970 | 0.04186937947 | 0.03724658724 | 1207 | Ldlrap1/Scarb1/ | Steppe |
| GLAND DEVELOPMENT | 236 | -0.4681533715 | -1.905592624 | 5.71E-07 | 0.00035563534 | 0.000316369692 | 2665 | Zbtb1/Esr1/Tgfb | Steppe |
| SENSORY ORGAN DEVELOPMENT | 263 | -0.4624780475 | -1.908852273 | 2.25E-07 | 0.000236531565 | 0.000210416149 | 1982 | Spry2/Jag1/Gata | Steppe |
| REGULATION OF CHONDROCYTE DIFFERENTIATI | 22 | -0.6966888437 | -1.915259466 | 0.001168655673 | 0.04548018328 | 0.04045872272 | 1529 | Zfp219/Zbtb16/P | Steppe |
| POSITIVE REGULATION OF PEPTIDE SECRETION | 47 | -0.6018384963 | -1.915403697 | 0.000353489428 | 0.02489098192 | 0.02214277215 | 1867 | Prkce/Orai1/Pfkf | Steppe |
| EMBRYONIC CAMERA TYPE EYE DEVELOPMENT | 22 | -0.6978009754 | -1.918316815 | 0.001150537346 | 0.04548018328 | 0.04045872272 | 1299 | Tbx2/Aldh1a3/Hi | Steppe |
| DITERPENOID METABOLIC PROCESS | 34 | -0.6405830407 | -1.920437031 | 0.000918560161 | 0.04021571205 | 0.03577550101 | 370 | Rdh10/Cyp1b1/F | Steppe |
| NEGATIVE REGULATION OF LIPID CATABOLIC PR | 13 | -0.7901973554 | -1.924082731 | 0.00121237701 | 0.04621878072 | 0.04111577172 | 570 | Acacb/Adora1/Pi | Steppe |
| RETINA MORPHOGENESIS IN CAMERA TYPE EYE | 29 | -0.6609230424 | -1.924269922 | 0.001027457046 | 0.04296917378 | 0.03822495342 | 1367 | Stat3/Ush1c/Ntrk | Steppe |
| NEGATIVE REGULATION OF HORMONE SECRETI | 23 | -0.6949101461 | -1.924707077 | 0.000680222293 | 0.03444572103 | 0.03064257388 | 870 | Irs1/Midn/Ffar2/ | Steppe |
| MAMMARY GLAND DEVELOPMENT | 80 | -0.5505315492 | -1.929233522 | 8.89E-05 | 0.0113238544 | 0.01007358925 | 1550 | Robo1/Stat5a/Pt | Steppe |
| POSITIVE REGULATION OF COLD INDUCED THER | 57 | -0.5875871982 | -1.934110461 | 0.000251924675 | 0.02104172775 | 0.01871851359 | 1714 | Stat6/Ogt/Elovl6/ | Steppe |
| CELL FATE COMMITMENT | 105 | -0.5287262483 | -1.937930827 | 6.61E-05 | 0.008814874203 | 0.007841625198 | 2517 | Sfrp1/Ep300/Paf | Steppe |
| FORMATION OF PRIMARY GERM LAYER | 69 | -0.5678018388 | -1.939758008 | 0.000210106883 | 0.01868950751 | 0.01662600165 | 2097 | Hnf1b/Axin1/Leo | Steppe |
| NERVOUS SYSTEM PROCESS | 401 | -0.4564541972 | -1.963340814 | 2.02E-10 | 1.13E-06 | 1.01E-06 | 1383 | Usp53/Itga3/Mbp | Steppe |
| TERPENOID METABOLIC PROCESS | 42 | -0.6339899814 | -1.967768501 | 0.000355331647 | 0.02489098192 | 0.02214277215 | 370 | Rdh10/Cyp1b1/F | Steppe |
| CAMERA TYPE EYE MORPHOGENESIS | 63 | -0.584949929 | -1.969546107 | 0.000169927056 | 0.01685661505 | 0.01499547858 | 1789 | Lhx1/Nf1/Abi2/F | Steppe |
| METANEPHRIC NEPHRON DEVELOPMENT | 24 | -0.7135710538 | -1.973942484 | 0.000393236128 | 0.02623446744 | 0.02333792362 | 2043 | Pkd2/Lhx1/Smo/ | Steppe |
| REGULATION OF LIPID STORAGE | 28 | -0.6825864252 | -1.97637342 | 0.000689426458 | 0.03444572103 | 0.03064257388 | 1184 | Scarb1/Abca1/Fi | Steppe |
| EYE MORPHOGENESIS | 76 | -0.5699650841 | -1.978887981 | 3.73E-05 | 0.005810063222 | 0.005168574969 | 1789 | Lhx1/Nf1/Abi2/F | Steppe |
| BLOOD VESSEL ENDOTHELIAL CELL MIGRATION | 80 | -0.5650231141 | -1.980016466 | 3.28E-05 | 0.00541772762 | 0.004819557085 | 2415 | Klf4/Pik3r3/Card | Steppe |
| REGULATION OF TRIGLYCERIDE BIOSYNTHETIC | 15 | -0.7889243534 | -1.985273161 | 0.000334151480 | 0.02400749868 | 0.02135683416 | 1184 | Scarb1/Rgn/Fitm | Steppe |
| EPIDERMIS DEVELOPMENT | 156 | -0.5136458783 | -1.993115786 | 1.01E-06 | 0.000565364615 | 0.000502942788 | 1316 | Jag1/Cflar/Ldb1/ | Steppe |
| RETINA DEVELOPMENT IN CAMERA TYPE EYE | 83 | -0.5642269166 | -1.994783432 | 3.73E-05 | 0.005810063222 | 0.005168574969 | 1452 | Mdm1/Smarca4/ | Steppe |
| FLUID TRANSPORT | 16 | -0.7816861179 | -1.99756893 | 0.000191985267 | 0.0182353464 | 0.0162219844 | 825 | Mllt6/Slc4a11/Aq | Steppe |
| REGULATION OF KIDNEY DEVELOPMENT | 18 | -0.7564270708 | -2.003215038 | 0.00061751514 | 0.03392700828 | 0.03018113213 | 2097 | Hnf1b/Gata3/Lhx | Steppe |
| DETECTION OF VISIBLE LIGHT | 9 | -0.9175873213 | -2.014917317 | 2.46E-05 | 0.004628294867 | 0.00411728549 | 314 | Sema5b/Reep6/ | Steppe |
| NEUTRAL LIPID METABOLIC PROCESS | 71 | -0.5877994768 | -2.022331726 | 4.87E-05 | 0.006825405915 | 0.00607181382 | 1390 | Lpin2/Abhd6/Sca | Steppe |
| REGULATION OF PEPTIDE TRANSPORT | 84 | -0.5774704484 | -2.036104224 | 2.34E-05 | 0.004628294867 | 0.00411728549 | 1895 | Slc25a22/Prkce/ | Steppe |
| SENSORY SYSTEM DEVELOPMENT | 194 | -0.5146026724 | -2.053816676 | 9.05E-08 | 0.000126858884 | 0.000112852413 | 1816 | Ift140/Mitf/Lhx1/ | Steppe |
| WATER TRANSPORT | 13 | -0.8454650618 | -2.058656251 | 0.000104663491 | 0.01234220961 | 0.01097950801 | 825 | Mllt6/Slc4a11/Aq | Steppe |
| TRIGLYCERIDE BIOSYNTHETIC PROCESS | 30 | -0.7155185837 | -2.079691182 | 3.29E-05 | 0.00541772762 | 0.004819557085 | 1390 | Lpin2/Scarb1/Rg | Steppe |
| SENSORY PERCEPTION | 185 | -0.5250346956 | -2.090317755 | 5.73E-08 | 0.000107081213 | 9.53E-05 | 1383 | Usp53/Mbp/Ush1 | Steppe |
| PROTEIN LIPID COMPLEX ORGANIZATION | 28 | -0.7221954232 | -2.091058049 | 0.000129879205 | 0.01323351029 | 0.0117724003 | 1184 | Scarb1/Abca1/Z | Steppe |
| DETECTION OF STIMULUS | 64 | -0.6254641949 | -2.111024015 | 6.11E-06 | 0.001902061937 | 0.00169205555 | 1251 | Pkd1/Jup/Scarb1 | Steppe |
| DETECTION OF LIGHT STIMULUS | 14 | -0.8498197436 | -2.114132176 | 2.98E-05 | 0.005210833286 | 0.004635505925 | 314 | Plekhb1/Sema5 | Steppe |
| PROTEIN LIPID COMPLEX ASSEMBLY | 17 | -0.8150153622 | -2.115905963 | 4.62E-05 | 0.006640732375 | 0.005907530059 | 967 | Abca1/Zdhhc8/S | Steppe |
| REGULATION OF PLASMA LIPOPROTEIN PARTICL | 41 | -0.6891501892 | -2.119391591 | 2.90E-05 | 0.005210833286 | 0.004635505925 | 1207 | Ldlrap1/Scarb1/ | Steppe |
| NEUTRAL LIPID BIOSYNTHETIC PROCESS | 34 | -0.7221032949 | -2.164830818 | 2.03E-05 | 0.004349369495 | 0.003869156228 | 1390 | Lpin2/Scarb1/Pla | Steppe |
| TRIGLYCERIDE METABOLIC PROCESS | 57 | -0.6628948062 | -2.181994065 | 2.81E-06 | 0.001211365382 | 0.001077618702 | 817 | Rgn/Fitm2/Gk5/F | Steppe |
| CELLULAR RESPONSE TO FATTY ACID | 20 | -0.8036833929 | -2.187757643 | 1.53E-05 | 0.003562795079 | 0.003169427381 | 870 | Irs1/Ffar2/Cpt1a/ | Steppe |
| SENSORY PERCEPTION OF LIGHT STIMULUS | 72 | -0.6424282503 | -2.210409554 | 2.68E-07 | 0.000236531565 | 0.000210416149 | 1316 | Ush1c/Ablim1/Cr | Steppe |
| RESPONSE TO FATTY ACID | 33 | -0.7403750099 | -2.214484598 | 8.40E-06 | 0.002354140621 | 0.00209422029 | 490 | Foxo3/Irs1/Ffar2 | Steppe |
| DETECTION OF ABIOTIC STIMULUS | 39 | -0.7245565769 | -2.227457379 | 4.92E-06 | 0.001622912838 | 0.001443727263 | 798 | Whrn/Plekhb1/Ki | Steppe |

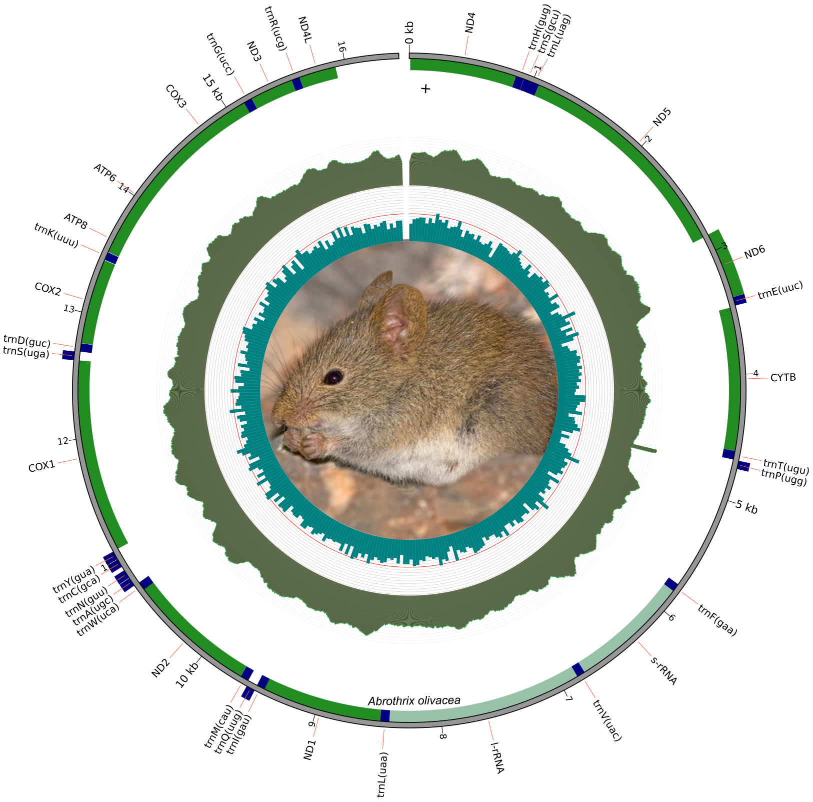


**Figure S1. Assembly and annotation of the mitochondrial genome of *A. olivacea*.**
**The complete circularized mitogenome was reconstructed from Illumina reads and annotated using standard mitochondrial gene models.**
**Protein-coding genes, rRNAs, and tRNAs are indicated with their respective orientation along the circular genome.**

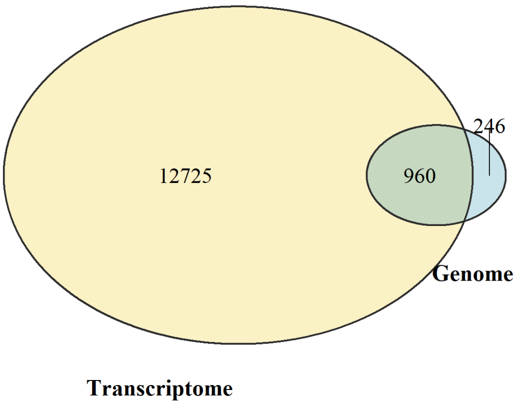


**Figure S2. Venn diagram comparing genes identified in Peromyscus eremicus through two complementary approaches: (i) detection based on the reference genome (Genome) and (ii) de novo transcriptome assembly (Transcriptome).**
**The diagram highlights the number of genes uniquely detected by each method and those found in both, illustrating differences in gene recovery between the two strategies.**

**Table S2. List of differentially expressed genes (DEGs) detected in Jaculus jaculus in response to contrasting hydration states.**
**For each gene, identifiers, log fold change values, adjusted p-values, and brief functional descriptions are provided.**
**This dataset complements the main text by detailing the transcriptional response of J. jaculus to dehydration.**

| **geneID** | **logFC** | **AveExpr** | **t** | **P.Value** | **adj.P.Val** | **B** |
|---|---|---|---|---|---|---|
| Kcnj15 | 1.93305431 | 10.12599071 | 17.23070638 | 2.71E-10 | 4.03E-06 | 12.94598157 |
| Grem2 | -3.242114229 | 4.334037896 | -12.80103896 | 1.03E-08 | 7.67E-05 | 9.955192345 |
| Slco4a1 | -1.668679704 | 6.144097867 | -9.491591056 | 3.46E-07 | 0.001717269328 | 7.079057667 |
| Cnnm1 | -2.904067471 | 3.113165679 | -8.823646682 | 7.89E-07 | 0.001768356799 | 5.523995995 |
| Slc6a12 | -1.538007766 | 5.417287257 | -8.955218208 | 6.68E-07 | 0.001768356799 | 6.437146827 |
| Crispld2 | -1.280377913 | 3.079370022 | -8.373238474 | 1.41E-06 | 0.001768356799 | 5.39192336 |
| Itga2 | -1.173190894 | 5.399517024 | -8.306750189 | 1.54E-06 | 0.001768356799 | 5.619462718 |
| Cryab | -1.09988595 | 8.659840768 | -8.432300723 | 1.31E-06 | 0.001768356799 | 5.784427338 |
| Vstm4 | -1.006900825 | 2.934938663 | -8.346416942 | 1.46E-06 | 0.001768356799 | 5.368319987 |
| Tnfaip2 | -0.9644876139 | 5.660364387 | -8.485531843 | 1.22E-06 | 0.001768356799 | 5.851322588 |
| LOC101611632 | 0.8021746823 | 5.552823226 | 8.55748907 | 1.11E-06 | 0.001768356799 | 5.941757451 |
| Clcn5 | 0.9252906071 | 8.075180924 | 8.316569929 | 1.52E-06 | 0.001768356799 | 5.635882764 |
| Nox4 | 1.457113845 | 8.86884974 | 8.772287356 | 8.42E-07 | 0.001768356799 | 6.207847611 |
| Fst | -2.231362201 | 3.892507555 | -7.970806351 | 2.42E-06 | 0.002212242304 | 5.163341266 |
| LOC101594411 | -0.9688608098 | 5.117306088 | -7.97661206 | 2.40E-06 | 0.002212242304 | 5.186388908 |
| Kiaa0040 | -0.9296430063 | 4.724988364 | -7.941353957 | 2.52E-06 | 0.002212242304 | 5.144346687 |
| Esrrb | -0.8270156733 | 7.035182531 | -7.942216565 | 2.52E-06 | 0.002212242304 | 5.124483871 |
| Ptgs2 | -3.21087266 | 1.387333269 | -7.749761862 | 3.29E-06 | 0.002504399214 | 3.558317199 |
| Cldn15 | -1.101543855 | 4.464289326 | -7.752138585 | 3.28E-06 | 0.002504399214 | 4.884031077 |
| Arhgap28 | -0.8109302935 | 4.355605224 | -7.733811049 | 3.36E-06 | 0.002504399214 | 4.863948113 |
| Hoxd3 | -1.037475921 | 4.568227575 | -7.688581872 | 3.58E-06 | 0.002540588759 | 4.788163128 |

| | | | | | | |
|---|---|---|---|---|---|---|
| Ctsa | 0.9461787147 | 8.217320434 | 7.579712507 | 4.17E-06 | 0.002826032853 | 4.644390007 |
| Slc4a7 | -0.9744412379 | 5.3799816 | -7.534008643 | 4.45E-06 | 0.002883732622 | 4.577671283 |
| Ngrn | -0.4619203893 | 6.153832908 | -7.36265286 | 5.68E-06 | 0.003529931087 | 4.310990396 |
| Wsb1 | -1.052474456 | 10.07123703 | -7.098907469 | 8.35E-06 | 0.004975013157 | 3.985908655 |
| Rasl11b | -0.9006238197 | 5.213444285 | -7.051876678 | 8.95E-06 | 0.005127488984 | 3.897758765 |
| Aqp3 | -1.736416577 | 6.839111477 | -6.984804799 | 9.88E-06 | 0.005259266934 | 3.769762229 |
| Steap2 | 1.497183344 | 3.668720213 | 6.990603214 | 9.80E-06 | 0.005259266934 | 3.832899125 |
| Cflar | 1.579080687 | 4.802562634 | 6.935823879 | 1.06E-05 | 0.005462586218 | 3.720405714 |
| LOC123459222 | -1.322823035 | 3.867754997 | -6.882143154 | 1.15E-05 | 0.005722533059 | 3.67189123 |
| Slc14a2 | -1.105316374 | 8.636290602 | -6.807500065 | 1.29E-05 | 0.006026856838 | 3.53768443 |
| Sowahb | -1.053557415 | 3.682550273 | -6.796196561 | 1.31E-05 | 0.006026856838 | 3.530341375 |
| Lrrc8a | 0.533606231 | 5.429905107 | 6.784552902 | 1.33E-05 | 0.006026856838 | 3.469615982 |
| Trim56 | 0.7132674055 | 3.902876043 | 6.744544725 | 1.42E-05 | 0.00621554671 | 3.474889695 |
| Tbx3 | -0.714841463 | 4.708793102 | -6.671077808 | 1.59E-05 | 0.00675343862 | 3.356719771 |
| Bdnf | -3.051738103 | 0.2261073964 | -6.497961083 | 2.07E-05 | 0.006782521665 | 1.449783565 |
| Tfap2b | -0.7754227699 | 6.00076968 | -6.499459626 | 2.07E-05 | 0.006782521665 | 3.02419288 |
| Cir1 | -0.6205884847 | 4.770306525 | -6.57675489 | 1.83E-05 | 0.006782521665 | 3.21580467 |
| Kctd17 | -0.5174972423 | 6.128156892 | -6.583259864 | 1.81E-05 | 0.006782521665 | 3.148512314 |
| Smcr8 | 0.5849555256 | 5.778233847 | 6.532525397 | 1.96E-05 | 0.006782521665 | 3.075709265 |
| Stk35 | 0.6835323981 | 5.47770513 | 6.490985111 | 2.09E-05 | 0.006782521665 | 3.02694453 |
| Tnrc6b | 0.7026493601 | 5.955084636 | 6.59739506 | 1.78E-05 | 0.006782521665 | 3.16661602 |
| LOC101613156 | 0.7492531052 | 8.554527071 | 6.56164495 | 1.88E-05 | 0.006782521665 | 3.171232184 |
| Fam13b | 0.7561800101 | 5.736533878 | 6.582836727 | 1.82E-05 | 0.006782521665 | 3.152731344 |
| Hook3 | 0.8725226494 | 4.101195843 | 6.5326262 | 1.96E-05 | 0.006782521665 | 3.158245611 |
| LOC101595540 | 1.034692514 | 3.937924103 | 6.512932089 | 2.02E-05 | 0.006782521665 | 3.133036819 |
| LOC123455959 | -4.43519378 | -1.43445886 | -6.238028057 | 3.11E-05 | 0.007060980369 | 0.5417952989 |
| Arhgap22 | -2.095748743 | 1.659400911 | -6.285068966 | 2.89E-05 | 0.007060980369 | 2.312745122 |
| Nr4a1 | -1.882115383 | 2.491042915 | -6.304711049 | 2.80E-05 | 0.007060980369 | 2.593938405 |
| Slc38a3 | -1.471707611 | 8.009313612 | -6.233405839 | 3.14E-05 | 0.007060980369 | 2.638414128 |
| LOC123458479 | -1.291287845 | 3.991061526 | -6.219433769 | 3.21E-05 | 0.007060980369 | 2.694997193 |
| Gal3st1 | -1.127243911 | 4.290570198 | -6.189706295 | 3.36E-05 | 0.007060980369 | 2.643695682 |
| Nos1 | -1.116174885 | 3.929664565 | -6.380425164 | 2.49E-05 | 0.007060980369 | 2.936806846 |
| Slc2a4 | -1.031517931 | 4.164659525 | -6.193353139 | 3.34E-05 | 0.007060980369 | 2.655953576 |
| Hoxd9 | -0.7587606791 | 5.771363703 | -6.206010168 | 3.28E-05 | 0.007060980369 | 2.577661183 |
| Dennd6b | -0.7573529064 | 4.68497821 | -6.232973464 | 3.14E-05 | 0.007060980369 | 2.6986441 |
| Fhl1 | -0.7441738217 | 5.76450477 | -6.264987153 | 2.98E-05 | 0.007060980369 | 2.661606679 |
| Cdc42ep3 | -0.7440759099 | 4.442386105 | -6.222853713 | 3.19E-05 | 0.007060980369 | 2.680774415 |
| Cldn4 | -0.608023915 | 6.0494422 | -6.32540319 | 2.71E-05 | 0.007060980369 | 2.753492028 |
| Naa80 | -0.5798488487 | 4.959025418 | -6.366425622 | 2.54E-05 | 0.007060980369 | 2.878336998 |
| Gata3 | -0.5360572581 | 5.50740273 | -6.34677869 | 2.62E-05 | 0.007060980369 | 2.809366317 |
| LOC101602550 | -0.5203469666 | 5.932608457 | -6.384315881 | 2.47E-05 | 0.007060980369 | 2.845638486 |
| Mga | 0.5602056509 | 6.109866885 | 6.296486066 | 2.84E-05 | 0.007060980369 | 2.693302636 |
| Smad7 | 0.5920657964 | 5.399799789 | 6.278512804 | 2.92E-05 | 0.007060980369 | 2.715889773 |
| Gxylt1 | 0.6915030309 | 2.859937671 | 6.208161805 | 3.27E-05 | 0.007060980369 | 2.661855385 |
| Etnk1 | 0.7332640049 | 6.523660371 | 6.292481298 | 2.86E-05 | 0.007060980369 | 2.688894582 |
| Osgin1 | 0.8167126235 | 7.706648739 | 6.399966871 | 2.41E-05 | 0.007060980369 | 2.879158643 |
| Ccdc117 | 1.135810153 | 4.497638207 | 6.20771427 | 3.27E-05 | 0.007060980369 | 2.639880723 |
| Tinag | 1.141218975 | 7.542755349 | 6.390639299 | 2.45E-05 | 0.007060980369 | 2.87035925 |
| LOC123456021 | 1.409674187 | 4.617317209 | 6.432871982 | 2.29E-05 | 0.007060980369 | 2.973123324 |
| Gadd45b | 1.972980279 | 4.965416329 | 6.40082891 | 2.41E-05 | 0.007060980369 | 2.935736122 |
| Cry2 | -1.063930329 | 5.182251836 | -6.138702218 | 3.65E-05 | 0.007350197757 | 2.499115785 |
| Cd9 | -0.7132868778 | 6.485342155 | -6.142727843 | 3.63E-05 | 0.007350197757 | 2.446789964 |
| Dnajb1 | 2.354443448 | 6.356087159 | 6.155706497 | 3.55E-05 | 0.007350197757 | 2.498769431 |
| Hbegf | -1.597199486 | 1.77398381 | -6.095553597 | 3.91E-05 | 0.00757035775 | 2.134061932 |
| Prox1 | -0.9126354699 | 5.300247798 | -6.099154344 | 3.89E-05 | 0.00757035775 | 2.422038664 |
| Bcl9l | 0.868462139 | 5.842559926 | 6.106114339 | 3.85E-05 | 0.00757035775 | 2.397607929 |
| Fez1 | -1.394108758 | 2.724852168 | -6.083407737 | 3.99E-05 | 0.00762077154 | 2.412583183 |
| Znf609 | 0.6077897319 | 6.287271816 | 6.067857254 | 4.09E-05 | 0.007715155889 | 2.323935894 |
| LOC101611263 | -0.5516166205 | 5.271777216 | -6.042115518 | 4.26E-05 | 0.007843797146 | 2.331698282 |
| Tmem245 | -0.4635110403 | 6.36219167 | -6.049018653 | 4.22E-05 | 0.007843797146 | 2.291724741 |
| LOC123454182 | -1.729332022 | 3.064204863 | -6.021392999 | 4.41E-05 | 0.007854872856 | 2.33453625 |
| C1qtnf1 | -1.016663979 | 4.165136598 | -6.017865305 | 4.43E-05 | 0.007854872856 | 2.374030869 |
| Mfsd4a | -0.5773125471 | 7.935530709 | -6.011443842 | 4.48E-05 | 0.007854872856 | 2.268478406 |
| Nr2f2 | -0.4698400073 | 6.830716855 | -6.026217848 | 4.37E-05 | 0.007854872856 | 2.255794949 |
| Tmigd1 | 0.7288510239 | 7.301964275 | 5.992200519 | 4.62E-05 | 0.007875582209 | 2.220647562 |
| Hspa5 | 1.17761643 | 9.359849671 | 5.99152823 | 4.63E-05 | 0.007875582209 | 2.29773049 |
| Bcl6 | 1.412449615 | 6.535870122 | 5.98842606 | 4.65E-05 | 0.007875582209 | 2.206063474 |
| Fam43b | -1.87391239 | 0.3833630206 | -5.971491096 | 4.78E-05 | 0.00788253743 | 1.510106072 |
| Atp1b3 | -0.8752073324 | 5.65528781 | -5.967722036 | 4.81E-05 | 0.00788253743 | 2.186982165 |
| Fa2h | -0.839328237 | 4.80440886 | -5.958358251 | 4.88E-05 | 0.00788253743 | 2.229414232 |
| Mpp7 | -0.7077879295 | 5.625869504 | -5.961752275 | 4.86E-05 | 0.00788253743 | 2.169813181 |
| Rfwd3 | 0.5560593389 | 4.692022584 | 5.953881507 | 4.92E-05 | 0.00788253743 | 2.2151515 |
| Fam20c | -0.6925756465 | 5.592223472 | -5.887481415 | 5.48E-05 | 0.008261422992 | 2.056730672 |
| Hnrnpdl | -0.5543533052 | 6.430079049 | -5.909599547 | 5.29E-05 | 0.008261422992 | 2.064151895 |
| Mtrfr | 0.6936349419 | 3.31975642 | 5.893185062 | 5.43E-05 | 0.008261422992 | 2.193644391 |
| Slc9a2 | 1.030214141 | 2.894040821 | 5.886822013 | 5.49E-05 | 0.008261422992 | 2.157632744 |
| Ptar1 | 1.05880317 | 2.616080146 | 5.907747427 | 5.30E-05 | 0.008261422992 | 2.19779076 |
| Tob2 | 1.080324294 | 7.305190398 | 5.895350231 | 5.41E-05 | 0.008261422992 | 2.061572872 |
| LOC101615653 | -1.495333824 | 3.262604718 | -5.847811763 | 5.85E-05 | 0.008492480533 | 2.101441037 |
| Hip1 | -0.9950765285 | 6.145440147 | -5.817780634 | 6.15E-05 | 0.008492480533 | 1.918788243 |
| Htra1 | -0.6986561894 | 6.826384772 | -5.828817497 | 6.04E-05 | 0.008492480533 | 1.933090805 |
| Slc12a9 | -0.6014988824 | 4.426017984 | -5.84794366 | 5.85E-05 | 0.008492480533 | 2.097712938 |
| Tasor2 | 0.4465592564 | 6.180392696 | 5.854755908 | 5.78E-05 | 0.008492480533 | 1.97262367 |
| Phkb | 0.4893034479 | 6.411871978 | 5.828829675 | 6.04E-05 | 0.008492480533 | 1.929887002 |
| Zc3h12c | 1.034991852 | 3.120462924 | 5.826766186 | 6.06E-05 | 0.008492480533 | 2.091517452 |
| Slc2a2 | 1.34703249 | 3.83461656 | 5.835582922 | 5.97E-05 | 0.008492480533 | 2.066881373 |

| | | | | | | |
|---|---|---|---|---|---|---|
| Gjb6 | 1.626226498 | 6.274958766 | 5.817029442 | 6.15E-05 | 0.008492480533 | 1.928737846 |
| Rnf182 | -2.450624653 | 1.940261755 | -5.783023545 | 6.51E-05 | 0.00866156968 | 1.683991428 |
| Vgf | -2.291130334 | 2.821941196 | -5.783759058 | 6.50E-05 | 0.00866156968 | 1.960981751 |
| Eva1b | -0.637304471 | 4.273740733 | -5.79004035 | 6.43E-05 | 0.00866156968 | 2.016407596 |
| LOC123462731 | 1.204703625 | 2.184563132 | 5.792849509 | 6.40E-05 | 0.00866156968 | 1.993416975 |
| Fyco1 | 0.3738217712 | 6.970985024 | 5.765989952 | 6.70E-05 | 0.008830040073 | 1.833782159 |
| Kctd12 | -0.7534739877 | 8.397141152 | -5.759534052 | 6.77E-05 | 0.008846568218 | 1.878772578 |
| Ddi2 | 0.744669252 | 3.405537068 | 5.753298326 | 6.84E-05 | 0.008860630241 | 1.971093454 |
| Hirip3 | -0.5569417141 | 5.285376119 | -5.711952592 | 7.32E-05 | 0.009170606226 | 1.797276367 |
| Pikfyve | 0.9010159823 | 5.505663127 | 5.719580326 | 7.23E-05 | 0.009170606226 | 1.767543202 |
| Mapk10 | 1.526969241 | 4.415495796 | 5.715282409 | 7.28E-05 | 0.009170606226 | 1.887792744 |
| Chl1 | 2.357936194 | 1.720695253 | 5.719317571 | 7.23E-05 | 0.009170606226 | 1.80080409 |
| Btbd8 | 0.7597315366 | 3.352554533 | 5.705621898 | 7.40E-05 | 0.009190372354 | 1.902499816 |
| Fabp4 | -1.63769103 | 6.663541453 | -5.68495044 | 7.66E-05 | 0.00935603069 | 1.709104331 |
| LOC123454581 | -0.8613449368 | 4.70531818 | -5.689462849 | 7.60E-05 | 0.00935603069 | 1.83090946 |
| Cep350 | 0.475583145 | 6.479830673 | 5.666046835 | 7.90E-05 | 0.009576983298 | 1.658310743 |
| Fabp5 | -1.109887493 | 4.806077561 | -5.65371168 | 8.07E-05 | 0.009592231264 | 1.769170638 |
| Slx4 | -0.7432105742 | 5.655782388 | -5.650655323 | 8.11E-05 | 0.009592231264 | 1.650433499 |
| Smg1 | 0.4911843862 | 6.271501138 | 5.657440174 | 8.02E-05 | 0.009592231264 | 1.643016974 |
| Nuak1 | -0.6795582809 | 4.285902345 | -5.615621059 | 8.60E-05 | 0.009907705147 | 1.710011178 |
| Hdac7 | -0.5934556881 | 6.010858222 | -5.599402375 | 8.84E-05 | 0.009907705147 | 1.560003229 |
| Nsun5 | -0.5880076104 | 4.65205507 | -5.611942202 | 8.65E-05 | 0.009907705147 | 1.686362548 |
| Fnip1 | 0.6008878231 | 6.539507391 | 5.607800392 | 8.71E-05 | 0.009907705147 | 1.562384355 |
| Tgoln2 | 0.7175406483 | 7.415374961 | 5.599016775 | 8.84E-05 | 0.009907705147 | 1.580497263 |
| Abcc2 | 1.588344459 | 8.032149232 | 5.611987015 | 8.65E-05 | 0.009907705147 | 1.625128901 |
| Ackr3 | 1.702533985 | 5.297905509 | 5.613162467 | 8.63E-05 | 0.009907705147 | 1.651060256 |
| Slf2 | 0.7826066139 | 5.560628886 | 5.580642671 | 9.12E-05 | 0.01011733631 | 1.53867248 |
| Pank3 | 0.8284790935 | 3.845214116 | 5.577653998 | 9.16E-05 | 0.01011733631 | 1.647182017 |
| Specc1 | -0.5996926714 | 5.205303535 | -5.565489917 | 9.35E-05 | 0.01025042866 | 1.546525826 |
| LOC101600794 | -1.652942186 | 7.089312973 | -5.545641909 | 9.67E-05 | 0.01044508781 | 1.473305852 |
| Tbc1d16 | -0.6997449766 | 7.325914488 | -5.549507933 | 9.61E-05 | 0.01044508781 | 1.479453831 |
| Fam174b | -0.6995813429 | 6.815513837 | -5.513894514 | 0.000102041435 | 0.01082187287 | 1.402254514 |
| Ehd4 | -0.5705807418 | 6.717831756 | -5.511889091 | 0.000102387712 | 0.01082187287 | 1.397376922 |
| Rala | -0.4740785026 | 6.279940456 | -5.516609955 | 0.000101574516 | 0.01082187287 | 1.405529854 |
| Pten | 0.6024787783 | 7.235288695 | 5.504323889 | 0.000103705130 | 0.01088392645 | 1.40646065 |
| Hk1 | -0.5127327592 | 7.564444779 | -5.496919146 | 0.00010501183 | 0.01094399512 | 1.394410566 |
| Map4k2 | -0.8666390377 | 4.521135078 | -5.487441169 | 0.000106709630 | 0.01104370569 | 1.499798908 |
| Mycbp2 | 0.4713567867 | 7.101003074 | 5.469462138 | 0.000110009795 | 0.01130673092 | 1.346662545 |
| Rrad | -1.130514356 | 3.604499468 | -5.46238682 | 0.000111337699 | 0.01136483384 | 1.515421571 |
| Fpgs | -0.5016287018 | 5.650183787 | -5.45066423 | 0.000113574920 | 0.01151433363 | 1.328705273 |
| Itpr2 | 0.5336613203 | 6.824231234 | 5.443596311 | 0.000114946538 | 0.01157465038 | 1.285443449 |
| Wnk4 | -0.6349708431 | 7.977370714 | -5.427714053 | 0.000118092411 | 0.01173287468 | 1.291527358 |
| Ass1 | 1.330589058 | 10.37625313 | 5.428726074 | 0.000117889288 | 0.01173287468 | 1.4019573 |
| Adamts16 | -1.979824217 | 2.136603098 | -5.414117237 | 0.000120857271 | 0.01191079886 | 1.388112085 |
| Disp1 | -0.4469847404 | 4.575397985 | -5.411091711 | 0.000121481676 | 0.01191079886 | 1.357915611 |
| Washc5 | 0.5017765133 | 6.145905557 | 5.403459438 | 0.000123071881 | 0.01198784476 | 1.211323773 |
| Mmd | -0.6493609318 | 3.964802563 | -5.393342994 | 0.000125213354 | 0.01211723781 | 1.380564865 |
| Foxc2 | -0.9154805188 | 3.377909011 | -5.378419343 | 0.000128443968 | 0.01229141526 | 1.382882649 |
| Vasn | -0.4669011658 | 4.821519757 | -5.374539765 | 0.000129298022 | 0.01229141526 | 1.274575315 |
| Atg14 | 0.528898571 | 4.931378728 | 5.373682696 | 0.000129487498 | 0.01229141526 | 1.242041029 |
| LOC101596230 | -0.6916299736 | 4.370194297 | -5.369259407 | 0.000130470013 | 0.01230629497 | 1.313744728 |
| Setbp1 | 0.8440667771 | 3.865081071 | 5.36353306 | 0.000131753577 | 0.01234920478 | 1.296081543 |
| Tor4a | -0.8408073296 | 2.879380244 | -5.339751059 | 0.000137227510 | 0.01244456665 | 1.312341845 |
| Als2cl | -0.6902973394 | 4.960773867 | -5.338148887 | 0.000137604744 | 0.01244456665 | 1.208880316 |
| Cntln | 0.4767338308 | 3.752784714 | 5.353814337 | 0.000133962393 | 0.01244456665 | 1.319511462 |
| Pggt1b | 0.5085554448 | 4.466765249 | 5.33740095 | 0.000137781218 | 0.01244456665 | 1.221242104 |
| Chd9 | 0.5748754915 | 6.57716657 | 5.346051095 | 0.000135754625 | 0.01244456665 | 1.113907502 |
| Znf189 | 0.6093992913 | 4.652321395 | 5.337623291 | 0.000137728732 | 0.01244456665 | 1.204530852 |
| Upk2 | -1.288973716 | 3.171578819 | -5.30520696 | 0.000145606426 | 0.01244664896 | 1.253589548 |
| Sncg | -0.9981742754 | 4.760579011 | -5.295503497 | 0.000148055177 | 0.01244664896 | 1.163706784 |
| Rbm3 | -0.9328398747 | 5.133730286 | -5.332304161 | 0.000138990123 | 0.01244664896 | 1.187017927 |
| Pmm1 | -0.891347703 | 6.103176393 | -5.327838505 | 0.000140058458 | 0.01244664896 | 1.091102715 |
| P3h3 | -0.7158160922 | 4.658497897 | -5.305237386 | 0.000145598815 | 0.01244664896 | 1.181753948 |
| LOC101599363 | -0.6591901492 | 8.67232978 | -5.294433771 | 0.000148327757 | 0.01244664896 | 1.0991077 |
| Eme2 | -0.6222251935 | 4.591999331 | -5.311173608 | 0.000144121771 | 0.01244664896 | 1.185456952 |
| Mecom | -0.4900510509 | 6.886123948 | -5.305903108 | 0.000145432384 | 0.01244664896 | 1.046620308 |
| Hspg2 | -0.4668609413 | 6.534558156 | -5.291779197 | 0.000149006447 | 0.01244664896 | 1.016225321 |
| Pawr | 0.4999092165 | 6.01932054 | 5.292370367 | 0.000148855023 | 0.01244664896 | 1.025548085 |
| Ptprd | 1.002451259 | 6.807157398 | 5.289869304 | 0.000149496756 | 0.01244664896 | 1.037184062 |
| Pah | 1.188872628 | 9.67729109 | 5.306205617 | 0.000145356823 | 0.01244664896 | 1.178633847 |
| Eepd1 | 1.288779978 | 4.465398982 | 5.294314586 | 0.000148358159 | 0.01244664896 | 1.195622481 |
| Klf9 | 1.482573033 | 4.564933788 | 5.30666196 | 0.000145242915 | 0.01244664896 | 1.149699251 |
| Bnc2 | 0.6182153409 | 4.228174131 | 5.284747133 | 0.000150820077 | 0.01248706456 | 1.151747461 |
| Mapre3 | -0.4797998608 | 5.099442372 | -5.281251968 | 0.000151730091 | 0.01249300302 | 1.093947344 |
| Psca | -1.161326591 | 3.616083996 | -5.263476999 | 0.000156447776 | 0.01281066603 | 1.194734063 |
| Mbtps1 | 0.4323944607 | 7.056518861 | 5.258186747 | 0.000157881316 | 0.0128574058 | 0.9721338261 |
| Cd24 | -0.7765815195 | 8.178096834 | -5.254489756 | 0.000158891271 | 0.01286932942 | 1.010599341 |
| Foxq1 | -1.224458192 | 2.391046198 | -5.247255812 | 0.000160887059 | 0.01296053973 | 1.114521869 |
| Adcy4 | -0.5543829774 | 5.123458675 | -5.242812786 | 0.000162125839 | 0.01299011498 | 1.020782118 |
| Acer2 | -0.9107471372 | 7.181376706 | -5.235431029 | 0.000164206086 | 0.01301682608 | 0.9399173862 |
| Arl15 | -0.5744981603 | 6.474153801 | -5.23838747 | 0.000163369602 | 0.01301682608 | 0.9241320488 |
| Slc33a1 | 0.4548355144 | 6.85148218 | 5.230415003 | 0.000165635564 | 0.01303384363 | 0.917940872 |
| Znf827 | 0.6105503767 | 3.095462182 | 5.228551575 | 0.000166169918 | 0.01303384363 | 1.134056573 |
| Tmem131l | -0.6088142497 | 5.550287873 | -5.209749759 | 0.000171663461 | 0.01305255391 | 0.9147155955 |
| LOC101593628 | -0.5447742931 | 4.099456613 | -5.223827612 | 0.000167532649 | 0.01305255391 | 1.088108993 |
| Marchf7 | -0.4013924241 | 6.289591367 | -5.210707829 | 0.000171378989 | 0.01305255391 | 0.8762275947 |

| | | | | | | |
|---|---|---|---|---|---|---|
| Tnks2 | 0.4551465535 | 6.322332283 | 5.216251351 | 0.000169742643 | 0.01305255391 | 0.8851492832 |
| Tmtc3 | 0.6366392713 | 5.344263462 | 5.210513449 | 0.000171436668 | 0.01305255391 | 0.8953157422 |
| LOC105944257 | 2.29929988 | -0.3194380672 | 5.210265742 | 0.000171510193 | 0.01305255391 | 0.2421568178 |
| Abcb4 | 0.7256272068 | 4.772584328 | 5.198469285 | 0.000175050257 | 0.01317562621 | 0.9589010996 |
| Plagl2 | 1.02571231 | 4.991954882 | 5.199114002 | 0.000174854821 | 0.01317562621 | 0.9404814447 |
| Casr | 0.7688861899 | 4.191904291 | 5.194057572 | 0.000176393747 | 0.01321003027 | 0.9786620915 |
| Zbtb33 | 0.4217246884 | 4.660004809 | 5.184224709 | 0.000179427062 | 0.01337000759 | 0.9467287391 |
| LOC101601970 | -0.5031364879 | 4.571933727 | -5.181091361 | 0.000180405078 | 0.01337600437 | 0.9911319501 |
| Dnajb4 | 1.13747073 | 5.903943465 | 5.175847987 | 0.000182054182 | 0.01343145287 | 0.8328311366 |
| Zdhhc20 | 0.6117190872 | 5.233188595 | 5.169883272 | 0.000183949330 | 0.0135044181 | 0.8539625954 |
| Hs3st3a1 | -1.611099133 | 2.493228545 | -5.155967759 | 0.000188451263 | 0.01360273865 | 1.000366044 |
| Ptprr | -1.172016865 | 2.986796196 | -5.154479226 | 0.000188939602 | 0.01360273865 | 1.021803691 |
| Trib2 | -0.7637105482 | 5.663241274 | -5.156420118 | 0.000188303120 | 0.01360273865 | 0.813358637 |
| Mrtfa | -0.5422799759 | 5.247000977 | -5.15499348 | 0.000188770743 | 0.01360273865 | 0.8550582176 |
| Oga | 0.3767650189 | 7.486802748 | 5.151149747 | 0.000190036690 | 0.01361594613 | 0.7987349345 |
| Abca2 | -0.6393749177 | 5.985565967 | -5.141943165 | 0.000193105116 | 0.01370402641 | 0.7653526552 |
| Dnaja1 | 1.405501223 | 6.669147344 | 5.14227297 | 0.000192994307 | 0.01370402641 | 0.7720518819 |
| Lamb2 | -0.6080472911 | 7.545842244 | -5.135005788 | 0.000195451422 | 0.01380479879 | 0.7693924105 |
| LOC101600843 | -1.309537494 | 8.363807301 | -5.124466536 | 0.000199073082 | 0.0139942743 | 0.7990551879 |
| Pappa2 | -1.990696341 | 6.351192484 | -5.098109525 | 0.000208440513 | 0.01402640892 | 0.7202411108 |
| LOC123453632 | -1.079653296 | 2.268766404 | -5.098861381 | 0.000208167017 | 0.01402640892 | 0.8829157071 |
| Arhgef25 | -0.8535168821 | 3.381702711 | -5.104036355 | 0.000206294699 | 0.01402640892 | 0.9385235564 |
| Large2 | -0.816525298 | 3.454440504 | -5.11477317 | 0.000202465974 | 0.01402640892 | 0.9549603299 |
| Itm2c | -0.7373060254 | 9.907186818 | -5.103006627 | 0.000206665850 | 0.01402640892 | 0.8338792632 |
| Amigo3 | -0.6793401761 | 2.701548337 | -5.098060778 | 0.000208458259 | 0.01402640892 | 0.9102868908 |
| Eng | -0.6735884456 | 6.626901875 | -5.095435187 | 0.000209416380 | 0.01402640892 | 0.674481898 |
| Irx1 | -0.6474117754 | 4.24595555 | -5.105674424 | 0.000205705713 | 0.01402640892 | 0.8781610768 |
| Irx2 | -0.5262540062 | 4.854455636 | -5.098388473 | 0.000208338999 | 0.01402640892 | 0.7953891541 |
| Trim13 | 0.5764735764 | 2.834061211 | 5.101397763 | 0.000207247140 | 0.01402640892 | 0.93502036 |
| Eya3 | 0.5949609825 | 3.659789969 | 5.094160503 | 0.000209883190 | 0.01402640892 | 0.868327449 |
| Ube2o | -0.4095971929 | 5.131980419 | -5.083825495 | 0.000213708396 | 0.01421828675 | 0.7476373576 |
| LOC123456016 | 1.61018586 | 1.113956377 | 5.078199738 | 0.000215821138 | 0.01429503299 | 0.6825766925 |
| Tmem170a | 0.5042796603 | 5.110231934 | 5.072171104 | 0.000218109402 | 0.01438267446 | 0.7006884488 |
| Marveld1 | -0.994882948 | 3.313733273 | -5.058925409 | 0.000223226429 | 0.01465525758 | 0.8613863856 |
| Fhl2 | -0.6206610554 | 5.234627892 | -5.052450461 | 0.000225773258 | 0.01471211983 | 0.6710271179 |
| Vps35 | 0.3931111136 | 6.97050902 | 5.050615794 | 0.000226500399 | 0.01471211983 | 0.6051497976 |
| Nrip1 | 0.9208357254 | 4.749626992 | 5.049222805 | 0.000227054120 | 0.01471211983 | 0.6733596452 |
| Plekhh3 | -0.6940439831 | 4.812329223 | -5.042907479 | 0.000229582279 | 0.01474769274 | 0.7183295589 |
| Xpo7 | 0.4524969164 | 7.169395911 | 5.043858447 | 0.000229199716 | 0.01474769274 | 0.6023758976 |
| Tnc | -1.263674618 | 5.271939127 | -5.029820892 | 0.000234915081 | 0.0148501655 | 0.644820785 |
| Scx | -0.9219384482 | 2.807953563 | -5.027781276 | 0.000235757800 | 0.0148501655 | 0.7999562657 |
| Glud1 | -0.8135608652 | 10.05115949 | -5.033152686 | 0.000233545220 | 0.0148501655 | 0.7279300179 |
| Dpp4 | 0.6347940811 | 8.79036766 | 5.033250828 | 0.000233504996 | 0.0148501655 | 0.6666821347 |
| Gpr146 | 0.7963755446 | 5.28267696 | 5.026811049 | 0.000236159781 | 0.0148501655 | 0.6242745031 |
| Arl5b | 0.613497419 | 5.511098866 | 5.019011722 | 0.000239417263 | 0.01499174571 | 0.5632518688 |
| Tjp3 | -0.6947182515 | 4.15349466 | -5.005652359 | 0.000245106334 | 0.01521599844 | 0.7106131712 |
| N4bp2 | 0.4782861309 | 6.144799433 | 5.00344097 | 0.000246061573 | 0.01521599844 | 0.5122990071 |
| Pik3ca | 0.5795421082 | 7.018697849 | 5.004251563 | 0.000245710976 | 0.01521599844 | 0.5275937782 |
| Ssh2 | 0.515954913 | 5.167400475 | 4.99583884 | 0.000249375143 | 0.01529398259 | 0.542979587 |
| Znf697 | 1.335820579 | 3.457195895 | 4.99769085 | 0.000248563635 | 0.01529398259 | 0.7179654073 |
| Sepsecs | 0.3019337446 | 5.778743058 | 4.993373257 | 0.000250459795 | 0.01529755057 | 0.4996492564 |
| Gem | -1.538216423 | 2.097402546 | -4.980926585 | 0.000256010910 | 0.01544216355 | 0.6704418344 |
| Megf9 | -0.4851021398 | 7.255731621 | -4.98398661 | 0.000254634375 | 0.01544216355 | 0.4960394936 |
| Endod1 | -0.4422596849 | 6.847145463 | -4.977988132 | 0.000257340062 | 0.01544216355 | 0.4694548234 |
| Pura | 0.4349642227 | 4.876679568 | 4.983160932 | 0.000255005039 | 0.01544216355 | 0.5681272052 |
| Ttc9c | 0.6386232003 | 2.748873748 | 4.976516633 | 0.000258008369 | 0.01544216355 | 0.7315505144 |
| Aqp2 | -1.410478581 | 7.750853055 | -4.963660552 | 0.000263924723 | 0.01565138036 | 0.5074625254 |
| Col6a1 | -0.7826281429 | 4.485593743 | -4.962095358 | 0.000264654623 | 0.01565138036 | 0.6272456606 |
| LOC101608112 | -0.6416300001 | 2.813339725 | -4.965345343 | 0.000263141396 | 0.01565138036 | 0.7100965577 |
| Acat2 | 0.6699344418 | 7.492234009 | 4.957899708 | 0.000266621589 | 0.0157053816 | 0.4553581699 |
| Epor | -0.7345263119 | 2.976542882 | -4.954440299 | 0.00026825486 | 0.01573937865 | 0.6956641114 |
| Ptgs1 | -1.346818278 | 4.172898176 | -4.946817059 | 0.000271890900 | 0.01582808628 | 0.6264220268 |
| Prpf40b | -0.4354410238 | 4.83571791 | -4.948670367 | 0.000271002230 | 0.01582808628 | 0.5484497904 |
| Hoxd4 | -0.6822842481 | 5.097839444 | -4.940134776 | 0.000275120388 | 0.015953771 | 0.5189550693 |
| Marf1 | 0.4231672612 | 6.412258782 | 4.928719449 | 0.000280730133 | 0.01615336364 | 0.3773133936 |
| Sytl2 | 0.7676773986 | 7.205658839 | 4.929011296 | 0.000280585238 | 0.01615336364 | 0.4007281119 |
| Cntf | -1.372164393 | 2.592044706 | -4.92556382 | 0.000282301818 | 0.0161813231 | 0.640409735 |
| Tppp3 | -1.238365981 | 1.916400584 | -4.914051864 | 0.000288113616 | 0.01641317577 | 0.5256649936 |
| Dnajb5 | 0.7890086879 | 4.74812077 | 4.913198398 | 0.000288549423 | 0.01641317577 | 0.5037227866 |
| Aloxe3 | -2.729917111 | -0.02648678047 | -4.906407405 | 0.000292041684 | 0.01648733844 | -0.1239759894 |
| Fbxl6 | -0.6883346318 | 6.097486744 | -4.90636071 | 0.000292065849 | 0.01648733844 | 0.3642364497 |
| Mcam | -0.4971624956 | 4.669533063 | -4.901683129 | 0.000294497091 | 0.01656184962 | 0.4758689918 |
| Atp6ap2 | 0.5526381362 | 8.384509744 | 4.89814188 | 0.000296351726 | 0.01660349539 | 0.4033213668 |
| LOC123456671 | -0.6726672167 | 6.637918393 | -4.892805139 | 0.000299169720 | 0.01665481965 | 0.3160454303 |
| LOC101596437 | 0.4149844161 | 7.049680709 | 4.892177632 | 0.000299502896 | 0.01665481965 | 0.3249766503 |
| Samhd1 | 0.7472108838 | 4.257652981 | 4.88839558 | 0.000301519185 | 0.01670461123 | 0.445729591 |
| Tle2 | -0.9172194998 | 5.267755829 | -4.884501172 | 0.000303610154 | 0.01675815605 | 0.4149218948 |
| Pde3b | -0.514701601 | 8.66824281 | -4.873090133 | 0.000309824328 | 0.01697860811 | 0.3765199927 |
| Rnasek | -0.4834008928 | 7.642384362 | -4.8720323 | 0.000310407069 | 0.01697860811 | 0.3063505448 |
| Skil | 0.8689542562 | 4.657984827 | 4.870918377 | 0.000311021942 | 0.01697860811 | 0.3782929702 |
| Slc4a11 | -1.016400886 | 6.306412966 | -4.867634414 | 0.000312842046 | 0.01698387279 | 0.2796991453 |
| Lratd1 | 0.6895715322 | 2.734423557 | 4.864593623 | 0.000314537267 | 0.01698387279 | 0.5481255468 |
| Rhou | 0.6957431696 | 4.949509193 | 4.866151373 | 0.000313667640 | 0.01698387279 | 0.3877977015 |
| Ybx3 | -0.6061348516 | 5.739741607 | -4.862533218 | 0.000315691362 | 0.01698465118 | 0.296384238 |
| Apcs | -1.239029566 | 7.848377202 | -4.857628072 | 0.000318456664 | 0.01707179738 | 0.2954613359 |
| Ifrd1 | -0.6555714899 | 5.425076526 | -4.853236963 | 0.000320953571 | 0.01708125769 | 0.2918782171 |

| | | | | | | |
|---|---|---|---|---|---|---|
| Wdr72 | -0.5878905341 | 7.073037579 | -4.85125128 | 0.000322089370 | 0.01708125769 | 0.2526341438 |
| Gcc2 | 0.5160067659 | 6.311276282 | 4.849285712 | 0.000323217786 | 0.01708125769 | 0.2354512083 |
| Slc7a8 | 1.233786664 | 9.121444112 | 4.855216981 | 0.000319825163 | 0.01708125769 | 0.3693803333 |
| Gper1 | -1.609321708 | 2.202166332 | -4.843029131 | 0.000326837116 | 0.01714798494 | 0.5014613571 |
| Zfp69 | -1.450188607 | 1.448734992 | -4.839182958 | 0.000329082982 | 0.01714798494 | 0.3032945975 |
| Isca1 | 0.3084372221 | 6.813109072 | 4.840842479 | 0.000328111987 | 0.01714798494 | 0.2229814795 |
| Phf24 | 1.489621426 | 3.177686276 | 4.840657514 | 0.000328220064 | 0.01714798494 | 0.4802440193 |
| LOC101595853 | -0.5649312316 | 4.606677603 | -4.835629188 | 0.000331172389 | 0.01719673213 | 0.354519576 |
| Grin2c | -1.847849076 | 3.256769297 | -4.821834755 | 0.000339414425 | 0.01723880191 | 0.4776773588 |
| Hoxb2 | -0.6088539753 | 5.65193548 | -4.822655306 | 0.000338918232 | 0.01723880191 | 0.2422309622 |
| Tnpo1 | 0.4690716548 | 7.584807027 | 4.83143228 | 0.000333657718 | 0.01723880191 | 0.2485073554 |
| Ccdc6 | 0.5950661324 | 6.169858612 | 4.822625266 | 0.000338936384 | 0.01723880191 | 0.1905581279 |
| Rmnd5a | 0.7252215471 | 4.948283825 | 4.823708784 | 0.000338282293 | 0.01723880191 | 0.23500414 |
| LOC123458818 | 0.8061042956 | 2.400814891 | 4.825932416 | 0.000336944057 | 0.01723880191 | 0.4791311327 |
| Megf10 | 1.326551536 | 5.181506 | 4.820736553 | 0.000340079699 | 0.01723880191 | 0.2383139596 |
| Znf518a | 0.5924167531 | 4.813688323 | 4.814171788 | 0.000344084936 | 0.01738270445 | 0.2600047279 |
| Sgms2 | 0.5991277789 | 3.372337548 | 4.810883803 | 0.000346109393 | 0.01742590641 | 0.3881461249 |
| Ppt2 | -0.4703111746 | 5.750296723 | -4.808618259 | 0.000347511528 | 0.01743759029 | 0.1980898003 |
| LOC101609267 | 1.46338527 | 8.714259495 | 4.804663329 | 0.000349973401 | 0.01750219328 | 0.2552051153 |
| Per1 | 1.508632707 | 5.83544354 | 4.79549573 | 0.000355750104 | 0.01773158462 | 0.1841886412 |
| Setd6 | -0.4986280528 | 4.463781365 | -4.793431121 | 0.000357064692 | 0.01773778371 | 0.3214760497 |
| Sbno1 | 0.5983141144 | 6.123984644 | 4.78923244 | 0.000359753709 | 0.01775301168 | 0.1284211984 |
| Zfand2a | 0.8740921353 | 3.065257595 | 4.789900308 | 0.000359324572 | 0.01775301168 | 0.4196763582 |
| Znf503 | -0.3851925002 | 6.102668105 | -4.784203486 | 0.000363002219 | 0.01785419825 | 0.1296272956 |
| Ccnt1 | 0.6776765508 | 2.903296011 | 4.779870699 | 0.000365825499 | 0.0179338731 | 0.3792107767 |
| Sybu | -1.113204572 | 2.208215823 | -4.768234239 | 0.000373521521 | 0.01801128764 | 0.3651857792 |
| Obsl1 | -0.4914469453 | 6.313464569 | -4.766148083 | 0.000374918949 | 0.01801128764 | 0.09168809284 |
| LOC101604209 | -0.3669481541 | 4.726572806 | -4.762947908 | 0.000377073189 | 0.01801128764 | 0.2375041465 |
| Fktn | 0.5160093058 | 3.738981425 | 4.764495657 | 0.000376029699 | 0.01801128764 | 0.3036589956 |
| Rfx7 | 0.5659455443 | 4.54323334 | 4.764937366 | 0.000375732450 | 0.01801128764 | 0.2126329079 |
| Mcl1 | 0.7675293353 | 8.040488259 | 4.775284404 | 0.000368838862 | 0.01801128764 | 0.1644443683 |
| Rad51b | 1.234603278 | 0.7108106506 | 4.770154753 | 0.000372239837 | 0.01801128764 | 0.04457099029 |
| Trim25 | 1.45865681 | 3.84898273 | 4.769390133 | 0.000372749568 | 0.01801128764 | 0.2783685312 |
| Baiap3 | 2.649682791 | 1.600496152 | 4.75997666 | 0.000379084849 | 0.01804952558 | 0.08926373367 |
| Jazf1 | -0.7557514052 | 4.409579981 | -4.754819327 | 0.000382603119 | 0.01810137868 | 0.2380917682 |
| Bicc1 | 0.37180507 | 8.239404635 | 4.755919396 | 0.000381849826 | 0.01810137868 | 0.1409946713 |
| Lmf2 | -0.3133375006 | 6.208564046 | -4.74915676 | 0.000386505185 | 0.01817062073 | 0.05884200968 |
| Sox6 | 0.7452735802 | 3.478322422 | 4.750389557 | 0.000385652157 | 0.01817062073 | 0.299591165 |
| Slc9a3r2 | -0.6925170905 | 6.510333923 | -4.73694495 | 0.000395061807 | 0.01848142397 | 0.03441732925 |
| Nras | 0.4702660242 | 4.795909403 | 4.736191038 | 0.000395596473 | 0.01848142397 | 0.1208028048 |
| LOC123463455 | -1.292525451 | 5.248894996 | -4.730681524 | 0.000399526666 | 0.01852809681 | 0.1166914093 |
| Sec31b | -1.006716535 | 3.151910943 | -4.729568926 | 0.000400325248 | 0.01852809681 | 0.3190619679 |
| Zfhx3 | 0.5425786386 | 3.877066781 | 4.730727509 | 0.000399493696 | 0.01852809681 | 0.2333494175 |
| Smpd2 | -0.4470318188 | 4.541064862 | -4.727289587 | 0.000401966458 | 0.01854645859 | 0.1973525235 |
| Hoxc4 | -0.7108705609 | 3.382470833 | -4.723590883 | 0.000404644563 | 0.018612401 | 0.3064777873 |
| Lrrc4c | -1.306889198 | 4.387689768 | -4.706352425 | 0.000417373171 | 0.01908009928 | 0.1516780325 |
| Cebpd | 0.8964994214 | 5.121094146 | 4.706410368 | 0.000417329696 | 0.01908009928 | 0.05720719917 |
| Klf6 | 0.7340301672 | 4.197283356 | 4.704366087 | 0.000418866354 | 0.01908980209 | 0.1094251946 |
| LOC123463481 | -1.061263445 | 0.29846423 | -4.69897775 | 0.000422944926 | 0.01920599214 | -0.2376433799 |
| Tmem25 | -0.6002767781 | 5.331887127 | -4.697601495 | 0.000423993250 | 0.01920599214 | 0.02493614851 |
| Gga3 | -0.5021338932 | 5.759342624 | -4.693367429 | 0.000427235393 | 0.0192942093 | -0.00481472924 |
| Pitpnm1 | -0.4671678599 | 6.449217366 | -4.685645594 | 0.000433214706 | 0.01950513222 | -0.06089979195 |
| Lclat1 | -0.5646898321 | 4.907101795 | -4.681196965 | 0.000436698852 | 0.01951614978 | 0.04543980949 |
| Slc16a10 | 0.6731416599 | 4.250673687 | 4.681300866 | 0.000436617146 | 0.01951614978 | 0.0997971991 |
| Manf | 0.8265166739 | 5.939270159 | 4.680321359 | 0.000437388044 | 0.01951614978 | -0.045022305 |
| Ust | -0.7925482954 | 3.052339845 | -4.674770013 | 0.000441783893 | 0.01959495646 | 0.2281504851 |
| LOC123461591 | 1.105410925 | 2.062046141 | 4.674989035 | 0.000441609593 | 0.01959495646 | 0.2348323321 |
| Ophn1 | 0.7064978553 | 4.457868132 | 4.668621379 | 0.000446706304 | 0.01975449274 | 0.04274420011 |
| LOC101607881 | 0.4301573127 | 7.742438311 | 4.665829718 | 0.000448959991 | 0.01979541644 | -0.05591938311 |
| Tent5a | 0.7674465704 | 3.9217183 | 4.655493068 | 0.000457408014 | 0.02010841194 | 0.08849324252 |
| Ctdsp2 | 0.5881116245 | 7.835072685 | 4.650889633 | 0.000461223312 | 0.02021650301 | -0.06661372832 |
| Nceh1 | 0.5542014898 | 6.462820512 | 4.649215911 | 0.00046261866 | 0.02021819909 | -0.1245463344 |
| Selenoi | 0.5821377116 | 3.666526829 | 4.644789998 | 0.000466329610 | 0.02032079002 | 0.08289228364 |
| LOC123455313 | -1.629929449 | 0.8583953096 | -4.639913315 | 0.000470454319 | 0.02033151406 | -0.1163133626 |
| Sox18 | -1.13294414 | 3.966438422 | -4.639661324 | 0.000470668479 | 0.02033151406 | 0.1163119063 |
| Cd36 | -0.9764875102 | 4.354335323 | -4.640694664 | 0.000469790913 | 0.02033151406 | 0.04087773569 |
| Egfl7 | -0.7182829647 | 5.280574678 | -4.636052313 | 0.000473746815 | 0.02034653832 | -0.05242438283 |
| Bdh1 | -0.5704865669 | 7.049612954 | -4.637273561 | 0.000472702809 | 0.02034653832 | -0.1403958852 |
| Upk1a | -1.370920809 | 2.675029643 | -4.627179794 | 0.000481403873 | 0.02037440667 | 0.1515429851 |
| LOC101605572 | -1.004439613 | 4.950475127 | -4.626857877 | 0.000481684093 | 0.02037440667 | -0.05893291104 |
| Rwdd2b | -0.4975065964 | 5.558299689 | -4.6258595 | 0.000482554230 | 0.02037440667 | -0.1078763623 |
| LOC101603871 | 0.3974037696 | 3.947441861 | 4.632696959 | 0.000476627541 | 0.02037440667 | 0.05190465438 |
| Mideas | 0.606118166 | 4.542918197 | 4.625808747 | 0.000482598507 | 0.02037440667 | -0.05495718973 |
| LOC101606456 | 0.8733028209 | 3.79561688 | 4.625932524 | 0.000482490530 | 0.02037440667 | 0.04005528939 |
| LOC105944219 | 0.420244785 | 4.150953018 | 4.623591557 | 0.000484536927 | 0.02039845713 | 0.01197769966 |
| Foxl1 | -1.119728812 | 0.8269946288 | -4.617470205 | 0.000489930693 | 0.02056742852 | -0.1537778523 |
| Lrp6 | 0.5860925845 | 4.680115432 | 4.614878877 | 0.000492232751 | 0.02060602443 | -0.1091753011 |
| Ephb3 | -0.7617852151 | 3.668518555 | -4.612285074 | 0.000494548250 | 0.02064496521 | 0.08603225435 |
| LOC101595240 | -2.83897864 | 0.2608027709 | -4.599467228 | 0.000506157856 | 0.02082351454 | -0.2723685831 |
| LOC123455739 | -1.211600501 | 5.863753725 | -4.598321779 | 0.000507209003 | 0.02082351454 | -0.158394588 |
| Ccdc80 | -0.7700960868 | 4.230293015 | -4.601716754 | 0.000504100100 | 0.02082351454 | -0.00749291609 |
| St13 | 0.4438008513 | 8.226648284 | 4.602113953 | 0.000503737664 | 0.02082351454 | -0.1339941092 |
| Patj | 0.4739825511 | 7.055264879 | 4.605243762 | 0.000500891220 | 0.02082351454 | -0.1899977001 |
| Amer1 | 0.5968866134 | 4.431927147 | 4.598505669 | 0.000507040099 | 0.02082351454 | -0.0681914624 |
| P3h2 | -0.7440646461 | 6.263296836 | -4.594375892 | 0.000510847405 | 0.02085796955 | -0.2211853238 |
| LOC101598792 | 1.809426967 | 9.378521723 | 4.595526522 | 0.000509783651 | 0.02085796955 | -0.0898568007 |

| | | | | | | |
|---|---|---|---|---|---|---|
| Mrps6 | -0.568932342 | 6.586347424 | -4.589538828 | 0.000515344508 | 0.02098409619 | -0.235073421 |
| Grem1 | -3.347000474 | -0.5671661389 | -4.578495666 | 0.000525766092 | 0.02115867336 | -0.717331371 |
| Foxc1 | -0.5556815514 | 5.02215155 | -4.579909853 | 0.000524419390 | 0.02115867336 | -0.1565755228 |
| Crip2 | -0.4398489678 | 5.794797438 | -4.581511978 | 0.000522898040 | 0.02115867336 | -0.2231491423 |
| Fhod3 | -0.3911197035 | 6.088250171 | -4.57861024 | 0.000525656852 | 0.02115867336 | -0.2527108399 |
| Dcaf1 | 0.3022372725 | 5.773370014 | 4.573419736 | 0.000530629411 | 0.02115867336 | -0.2489660878 |
| LOC101599483 | 0.3644039798 | 5.685238903 | 4.571251621 | 0.000532720908 | 0.02115867336 | -0.2440735977 |
| Lcor | 0.4752744519 | 6.66966154 | 4.57298891 | 0.000531044332 | 0.02115867336 | -0.2617093878 |
| C4H3orf38 | 0.5401870435 | 2.910349941 | 4.571887577 | 0.000532106541 | 0.02115867336 | 0.04702075492 |
| Hnf4g | 1.011563981 | 6.709837252 | 4.570105986 | 0.000533829509 | 0.02115867336 | -0.249337273 |
| Ar | 1.317822753 | 2.713517083 | 4.575815745 | 0.000528327974 | 0.02115867336 | 0.03496656396 |
| LOC101611863 | 1.062184623 | 6.418113124 | 4.566326474 | 0.000537503838 | 0.02124779761 | -0.2691202613 |
| Plat | -0.7391762597 | 7.520542472 | -4.557660486 | 0.000546027987 | 0.02147085776 | -0.2617472443 |
| Hsp90b1 | 0.5866124024 | 9.34695138 | 4.558002988 | 0.000545688444 | 0.02147085776 | -0.1568468444 |
| Samd8 | 0.790488573 | 4.20579883 | 4.554103631 | 0.000549567082 | 0.02155315325 | -0.1526811774 |
| Sbf2 | 0.3793144634 | 5.841322993 | 4.549408553 | 0.000554275167 | 0.0216807423 | -0.3023991235 |
| Zim3 | 0.6782616608 | 3.648834986 | 4.547372881 | 0.000556329454 | 0.02170413053 | -0.06871830519 |
| Ptk7 | -0.5996907922 | 4.45442404 | -4.5412109 | 0.000562596021 | 0.02176867362 | -0.1292957492 |
| Dynll2 | 0.4077106425 | 6.431651091 | 4.540009206 | 0.000563826613 | 0.02176867362 | -0.3264069053 |
| Plekhm3 | 0.8324820104 | 2.903272385 | 4.542576312 | 0.000561201152 | 0.02176867362 | -0.02180894948 |
| Prune2 | 1.629588299 | 6.837898176 | 4.542283586 | 0.000561499891 | 0.02176867362 | -0.294515401 |
| Slc49a3 | 0.7559475567 | 4.611667904 | 4.537901914 | 0.000565991320 | 0.02179578461 | -0.1623515002 |
| A1cf | 0.9726785572 | 5.120524835 | 4.534221591 | 0.000569792600 | 0.0218856163 | -0.286554297 |
| Hoxb3 | -0.580070875 | 4.013770818 | -4.532572792 | 0.000571504161 | 0.0218949268 | -0.08418991015 |
| Agpat2 | -0.6214921543 | 4.64958654 | -4.518196194 | 0.000586655676 | 0.02241776806 | -0.1957260191 |
| Fndc10 | -1.550027837 | 0.09891119102 | -4.51101918 | 0.000594374711 | 0.02265464532 | -0.5714961394 |
| LOC101611810 | 0.5549333311 | 8.068172959 | 4.509476194 | 0.000596047925 | 0.02266046489 | -0.3165828531 |
| Plin1 | -1.978273458 | 3.089940438 | -4.505106981 | 0.000600812422 | 0.02276925435 | -0.05365828839 |
| Gdpd2 | -1.071830896 | 2.721344262 | -4.504055307 | 0.000601965122 | 0.02276925435 | -0.05113803485 |
| Phc3 | 1.068952777 | 2.704344844 | 4.501577923 | 0.000604689551 | 0.02281440096 | -0.07074197788 |
| Tbc1d25 | -0.5121629506 | 4.417320177 | -4.495468616 | 0.000611462800 | 0.02301169222 | -0.2039006764 |
| Tnxb | -0.7715841314 | 5.185159396 | -4.487666265 | 0.000620227522 | 0.02305050067 | -0.3208557542 |
| Hmgcr | 0.5699758148 | 7.003643707 | 4.488568586 | 0.000619207287 | 0.02305050067 | -0.4038964332 |
| Samd15 | 0.7203956043 | 3.159830232 | 4.489453354 | 0.000618208585 | 0.02305050067 | -0.09898684177 |
| Idh1 | 0.7225358448 | 9.036921045 | 4.492653546 | 0.000614610204 | 0.02305050067 | -0.2862216825 |
| Ypel2 | 1.302949705 | 3.975246566 | 4.490583069 | 0.000616935818 | 0.02305050067 | -0.1938645263 |
| Chpf2 | -0.6293109177 | 3.964387562 | -4.480112061 | 0.000628837500 | 0.0232512956 | -0.1795371493 |
| Lmna | -0.5916043576 | 5.41266419 | -4.478830067 | 0.000630310905 | 0.0232512956 | -0.3610404597 |
| Hspa8 | 1.235513183 | 10.91133691 | 4.479685008 | 0.000629327918 | 0.0232512956 | -0.2077027647 |
| Emilin1 | -0.7576236853 | 5.102317554 | -4.474898191 | 0.000634852173 | 0.02336082227 | -0.3693554428 |
| Gcnt1 | -0.6356528724 | 6.283250459 | -4.473551749 | 0.000636415073 | 0.02336082227 | -0.4467810678 |
| Scarf2 | -1.06583302 | 2.328181175 | -4.468179168 | 0.000642691134 | 0.02345892804 | -0.12147772 |
| Tmem184b | -0.4503547606 | 5.834285302 | -4.466666821 | 0.000644469337 | 0.02345892804 | -0.4388438925 |
| Atp11b | 0.3939648122 | 6.623171887 | 4.465890434 | 0.000645384184 | 0.02345892804 | -0.4567411859 |
| Rasal2 | 0.4808802103 | 3.743569958 | 4.468431406 | 0.000642395050 | 0.02345892804 | -0.2227732351 |
| LOC101617211 | -1.496716457 | 6.410422938 | -4.461858595 | 0.000650156717 | 0.02346073987 | -0.4492570771 |
| Hnrnph3 | -0.4760472623 | 7.39224091 | -4.462589386 | 0.000649288968 | 0.02346073987 | -0.4505447852 |
| LOC101595102 | 0.3394835824 | 5.855222679 | 4.463638059 | 0.000648045857 | 0.02346073987 | -0.4533075563 |
| Myoz1 | -1.127693091 | 1.336575143 | -4.451970352 | 0.000662016844 | 0.02371643518 | -0.297227222 |
| Slc17a5 | 0.2671662875 | 5.333569976 | 4.453560602 | 0.000660094464 | 0.02371643518 | -0.4414326647 |
| Slc7a7 | 0.6835411635 | 7.552164807 | 4.452503702 | 0.000661371458 | 0.02371643518 | -0.4475909373 |
| Lysmd3 | 0.5459604545 | 5.273126103 | 4.450592759 | 0.000663686838 | 0.02371924448 | -0.4382582582 |
| Crbn | 0.3515017944 | 4.208064268 | 4.449095141 | 0.000665507279 | 0.02372740429 | -0.310469126 |
| C5H21orf91 | 0.739333712 | 2.497216746 | 4.446079461 | 0.000669188709 | 0.02374504605 | -0.1536044186 |
| Pdk4 | 2.967422856 | 4.735904251 | 4.447357178 | 0.000667626354 | 0.02374504605 | -0.3537655567 |
| Ly6g5b | -1.059708773 | 1.267838408 | -4.438734213 | 0.000678243964 | 0.02391638159 | -0.3050142212 |
| Slc6a9 | -0.7321476222 | 3.258397995 | -4.435684087 | 0.000682041345 | 0.02391638159 | -0.192217671 |
| Dcaf15 | -0.560444289 | 5.422046496 | -4.440783368 | 0.000675705076 | 0.02391638159 | -0.4214316643 |
| Atf2 | 0.382007352 | 6.148236712 | 4.436874307 | 0.000680556920 | 0.02391638159 | -0.514295386 |
| Lzic | 0.4204589058 | 4.757056679 | 4.437426274 | 0.000679869651 | 0.02391638159 | -0.4215062252 |
| Anks1a | -0.3417440151 | 6.42246741 | -4.433069891 | 0.000685313501 | 0.02397471154 | -0.5230642235 |
| LOC123461645 | -1.735815667 | 0.6999000469 | -4.42441557 | 0.000696262375 | 0.02430069832 | -0.606126998 |
| Ckap4 | -0.7559712025 | 3.553339174 | -4.420748357 | 0.000700956331 | 0.02439096497 | -0.2403610271 |
| Nr2f1 | -0.7243334494 | 4.268337977 | -4.419841611 | 0.000702121987 | 0.02439096497 | -0.3284378414 |
| LOC123455308 | -0.957068315 | 3.704915096 | -4.417854206 | 0.000704683890 | 0.02442303259 | -0.2838576126 |
| Spon1 | -0.5872423855 | 4.757981122 | -4.416284062 | 0.000706714750 | 0.02443658915 | -0.422947628 |
| Rasgrp2 | -0.7590869218 | 2.489838488 | -4.413399791 | 0.000710461122 | 0.02450398528 | -0.20311675 |
| Dnajc13 | 0.3809205033 | 7.517415333 | 4.411567722 | 0.000712851455 | 0.02450398528 | -0.519331437 |
| Mtmr3 | 0.4869755693 | 5.970346696 | 4.410997931 | 0.000713596565 | 0.02450398528 | -0.5596248868 |
| Timp3 | -0.4323383045 | 9.929345531 | -4.408508694 | 0.000716861178 | 0.02450927903 | -0.3948015093 |
| Ciao2a | 0.3610142558 | 6.379808662 | 4.408372758 | 0.000717039901 | 0.02450927903 | -0.5680209497 |
| Plekhf2 | 0.8040700103 | 2.203210318 | 4.407042702 | 0.000718791037 | 0.02451291265 | -0.2113081523 |
| Arfgap2 | -0.3427909478 | 6.779471209 | -4.400729097 | 0.000727163984 | 0.02468547805 | -0.5812190505 |
| Arhgap42 | 0.6502305138 | 5.221365196 | 4.40109028 | 0.000726682283 | 0.02468547805 | -0.5329649022 |
| Fam126b | 1.387365674 | 5.184378571 | 4.399353711 | 0.000729001329 | 0.0246916064 | -0.5409126918 |
| Pkm | -0.4365361164 | 8.698382298 | -4.398044733 | 0.000730754419 | 0.02469485968 | -0.4915936335 |
| Cacul1 | 0.3812141452 | 6.140188521 | 4.395806994 | 0.000733761486 | 0.0247403788 | -0.5895667865 |
| Slc12a6 | 0.7420108137 | 7.115064647 | 4.394160489 | 0.000735982224 | 0.02475923947 | -0.5668899266 |
| Lrrc28 | -0.4063794866 | 4.000846544 | -4.392156193 | 0.000738694916 | 0.02479452779 | -0.3645022934 |
| Mcm8 | -0.4987019123 | 3.204103463 | -4.388285172 | 0.000743963409 | 0.02480377337 | -0.2821373878 |
| Kif16b | 0.4418116039 | 6.055703991 | 4.389119505 | 0.000742824600 | 0.02480377337 | -0.6026291543 |
| Jun | 0.851747073 | 5.506330683 | 4.389588402 | 0.000742185377 | 0.02480377337 | -0.5476359962 |
| Lpcat3 | 0.4060460772 | 7.487576349 | 4.383181258 | 0.000750969323 | 0.02498146389 | -0.5782517796 |
| Ift122 | -0.5084582153 | 5.412844409 | -4.376875446 | 0.000759719316 | 0.02508789653 | -0.5710155097 |
| Pex16 | -0.5038039056 | 4.713112578 | -4.376507311 | 0.000760233388 | 0.02508789653 | -0.4772896061 |
| Gprc5c | -0.438449357 | 6.883316711 | -4.378939639 | 0.000756843482 | 0.02508789653 | -0.6178268297 |

| | | | | | | |
|---|---|---|---|---|---|---|
| Clcn4 | 0.664022894 | 5.152620151 | 4.37602857 | 0.000760902451 | 0.02508789653 | -0.5345607651 |
| Akap8 | -0.6563559283 | 4.814141623 | -4.37268841 | 0.000765587442 | 0.02518664383 | -0.4691760466 |
| Scnm1 | -0.4199648868 | 3.630523087 | -4.368033817 | 0.000772165869 | 0.02534711002 | -0.3315960984 |
| LOC101598862 | -2.000918618 | 0.03474075237 | -4.361731644 | 0.000781166076 | 0.02553008342 | -0.7081305098 |
| LOC123456015 | 1.168088273 | 4.258294095 | 4.361923182 | 0.000780890946 | 0.02553008342 | -0.4493074376 |
| Edem1 | 0.8088380658 | 4.572093031 | 4.360104121 | 0.000783507926 | 0.02555058782 | -0.5572355078 |
| Pak3 | -1.867080012 | 0.7547471852 | -4.356339215 | 0.000788953113 | 0.02567198309 | -0.4943124544 |
| Snrnp70 | -0.5222681112 | 7.94641375 | -4.353016534 | 0.000793791191 | 0.02571710898 | -0.6323521452 |
| Trps1 | 0.7552789336 | 4.418854557 | 4.353916789 | 0.000792477331 | 0.02571710898 | -0.5110637893 |
| LOC101613101 | -0.5067888774 | 8.403602802 | -4.345784154 | 0.000804428364 | 0.0260051972 | -0.5958522964 |
| Thbs2 | -1.246291353 | 2.547472192 | -4.342123479 | 0.000809868402 | 0.02612439135 | -0.3256305454 |
| Ppargc1a | 0.5276144731 | 7.37208336 | 4.340538063 | 0.000812236232 | 0.02614418266 | -0.6646137587 |
| Atp1b1 | -0.4214742485 | 11.68506292 | -4.332916915 | 0.000823718680 | 0.02643079487 | -0.4446816593 |
| Tnrc6a | 0.4594055779 | 6.251506993 | 4.332278836 | 0.000824687621 | 0.02643079487 | -0.7070092535 |
| Osbpl11 | 0.4202082878 | 4.493308153 | 4.330539193 | 0.000827335302 | 0.02645875109 | -0.5666731812 |
| Cand1 | 0.3244746707 | 6.609342802 | 4.329230527 | 0.000829332835 | 0.02646583991 | -0.7128140574 |
| Itpkb | -0.5639370964 | 5.534183697 | -4.327946752 | 0.000831297216 | 0.02647184278 | -0.6815784133 |
| Fnip2 | 0.5382560501 | 6.205949079 | 4.323222384 | 0.000838567729 | 0.02664642828 | -0.7215385285 |
| LOC123462787 | -1.002384932 | 1.51401448 | -4.320084846 | 0.000843432466 | 0.02674398735 | -0.4378359671 |
| Mmp28 | -1.110974156 | 2.875240184 | -4.313332658 | 0.000854000797 | 0.02686153506 | -0.3683571469 |
| Abca7 | -0.8530627871 | 3.460707575 | -4.313168784 | 0.000854258982 | 0.02686153506 | -0.417323353 |
| Zbtb8a | -0.7560313799 | 2.337208384 | -4.31311147 | 0.000854349300 | 0.02686153506 | -0.3703566612 |
| Ephb2 | 1.111407944 | 2.98647841 | 4.314320073 | 0.000852446829 | 0.02686153506 | -0.4263823313 |
| Rusc2 | 0.6556835131 | 4.545304458 | 4.311082074 | 0.000857553652 | 0.02690552018 | -0.5790878035 |
| Znf81 | 0.524291101 | 2.611686269 | 4.30829884 | 0.000861968484 | 0.02693064218 | -0.3937889361 |
| Atp7a | 0.6262277391 | 5.395151895 | 4.308319992 | 0.000861934843 | 0.02693064218 | -0.7128970166 |
| Nipbl | 0.4586989187 | 6.048519894 | 4.306202088 | 0.000865309894 | 0.02694053191 | -0.7560028194 |
| Klf7 | 0.8436045765 | 1.268972791 | 4.305832365 | 0.000865900475 | 0.02694053191 | -0.4000524962 |
| LOC123459232 | -0.6205971536 | 3.957615861 | -4.290099613 | 0.000891420821 | 0.02709080726 | -0.522471903 |
| Lpin1 | -0.5987877234 | 4.511521379 | -4.296645693 | 0.000880709291 | 0.02709080726 | -0.609263372 |
| Anxa2 | -0.5856761298 | 8.208415061 | -4.296384082 | 0.000881134814 | 0.02709080726 | -0.7075551083 |
| Rex1bd | -0.5132293452 | 4.207918008 | -4.296108854 | 0.000881582714 | 0.02709080726 | -0.535035234 |
| Sim2 | -0.4460942019 | 4.697172257 | -4.290204198 | 0.000891248633 | 0.02709080726 | -0.6493803081 |
| Ufc1 | -0.2807714659 | 6.447448381 | -4.294327334 | 0.000884487611 | 0.02709080726 | -0.7804732418 |
| Ppp4r2 | 0.3331963815 | 6.490864757 | 4.29429344 | 0.000884542972 | 0.02709080726 | -0.7795712608 |
| LOC101612114 | 0.3597819837 | 6.654059607 | 4.290638954 | 0.000890533221 | 0.02709080726 | -0.7837685283 |
| Bbx | 0.5855627384 | 5.114192797 | 4.300414426 | 0.000874602758 | 0.02709080726 | -0.7176524331 |
| Amd1 | 0.5925913999 | 6.184971597 | 4.2905777 | 0.000890633981 | 0.02709080726 | -0.7842164321 |
| Mertk | 0.8291558293 | 7.153320047 | 4.289417791 | 0.000892544210 | 0.02709080726 | -0.7636486344 |
| LOC101612958 | 1.566050206 | 2.209213486 | 4.289449742 | 0.000892491534 | 0.02709080726 | -0.4257891552 |
| Slc2a1 | -0.5103704762 | 6.325855678 | -4.286944156 | 0.000896632121 | 0.02714341369 | -0.7878640004 |
| Cipc | 0.366628833 | 5.915692779 | 4.286052213 | 0.000898110876 | 0.02714341369 | -0.7823561094 |
| Pls1 | 0.6320306194 | 5.936788247 | 4.285070471 | 0.000899741418 | 0.02714341369 | -0.7919641989 |
| Acp6 | -1.304818905 | 4.306909359 | -4.281316485 | 0.000906004514 | 0.02727714197 | -0.568397022 |
| Faim2 | -2.104191665 | 0.7615187392 | -4.277999299 | 0.000911576296 | 0.02730952781 | -0.6637441621 |
| St3gal1 | -0.6497396052 | 5.462644956 | -4.277668439 | 0.000912133969 | 0.02730952781 | -0.7537533324 |
| Mark1 | -0.6242558787 | 4.150678907 | -4.276320173 | 0.000914410144 | 0.02730952781 | -0.5735687502 |
| Srsf2 | -0.3202005121 | 7.351141589 | -4.276492509 | 0.000914118875 | 0.02730952781 | -0.7995399104 |
| Foxk1 | 0.5454130825 | 3.690481939 | 4.272269418 | 0.000921284044 | 0.02745979223 | -0.5661204243 |
| Gja4 | -0.5998442431 | 3.67215204 | -4.268751518 | 0.000927296971 | 0.02752889793 | -0.5212481222 |
| Atm | 0.3845911497 | 4.783417639 | 4.269815785 | 0.000925473626 | 0.02752889793 | -0.7109456828 |
| LOC101603562 | -0.6679459051 | 3.307701714 | -4.265715605 | 0.000932518608 | 0.02762887639 | -0.5130915265 |
| Pde10a | -0.7084714959 | 2.657249631 | -4.2637786 | 0.000935866015 | 0.02767303815 | -0.465707277 |
| Mfap4 | -0.9396156553 | 4.79678782 | -4.262366732 | 0.000938313723 | 0.0276904741 | -0.6874907877 |
| Nectin4 | -0.7215984569 | 2.145171724 | -4.259054899 | 0.000944081281 | 0.02775077582 | -0.4697813881 |
| Chml | 0.99014965 | 3.512799219 | 4.259367668 | 0.000943535034 | 0.02775077582 | -0.5751604872 |
| Wdfy2 | 0.9030004198 | 4.380845904 | 4.255207471 | 0.000950827519 | 0.02789406008 | -0.7050699963 |
| LOC123464113 | -3.455296494 | 7.37455125 | -4.251928723 | 0.000956615847 | 0.02789911149 | -0.7611553537 |
| Zhx1 | 0.5970292815 | 5.646804435 | 4.252407347 | 0.000955768619 | 0.02789911149 | -0.8397798074 |
| Atxn1 | 0.7772144338 | 2.849619697 | 4.253702544 | 0.000953479830 | 0.02789911149 | -0.5235135622 |
| Def8 | 0.3351750023 | 7.31759439 | 4.246591665 | 0.000966115771 | 0.02812113935 | -0.8435075568 |
| Ccdc107 | -0.5374611039 | 6.052446664 | -4.242615977 | 0.000973255660 | 0.02821873367 | -0.855432752 |
| Elmod2 | 0.7483908346 | 4.04512748 | 4.243446813 | 0.000971759088 | 0.02821873367 | -0.6917426662 |
| Stxbp1 | -0.7087861631 | 3.823396025 | -4.234144506 | 0.000988651533 | 0.02860946371 | -0.6313649828 |
| Shisa2 | -1.580222123 | 2.936386792 | -4.229185823 | 0.000997779633 | 0.02865107877 | -0.5282442014 |
| Acap3 | -0.6424887877 | 3.823231903 | -4.229896588 | 0.000996465918 | 0.02865107877 | -0.614485613 |
| Ubtd1 | -0.4303154284 | 5.051947097 | -4.230117801 | 0.000996057412 | 0.02865107877 | -0.796351795 |
| Mipol1 | 0.5167658196 | 3.578210466 | 4.230419408 | 0.000995500723 | 0.02865107877 | -0.6109037315 |
| Ptprn | -1.79567192 | 1.936815019 | -4.219953595 | 0.001015006785 | 0.02886764525 | -0.5285736989 |
| Virma | 0.3000166919 | 5.208361837 | 4.220058791 | 0.001014808773 | 0.02886764525 | -0.8616887853 |
| Chd2 | 0.4507739862 | 5.21984719 | 4.220042096 | 0.001014840196 | 0.02886764525 | -0.8693796618 |
| Smim34 | 1.35541972 | 3.067048811 | 4.220208158 | 0.001014527688 | 0.02886764525 | -0.5319364136 |
| Amy2b | 1.394423884 | 5.196878756 | 4.222618698 | 0.001010002507 | 0.02886764525 | -0.7921700999 |
| Pias3 | -0.4803942181 | 4.871888358 | -4.216585584 | 0.001021367492 | 0.02899321854 | -0.7849519913 |
| Fgf12 | -1.126346361 | 4.336379487 | -4.214483628 | 0.001025357981 | 0.02901127158 | -0.7274486524 |
| Fmnl3 | -0.3870527915 | 5.536787697 | -4.214200452 | 0.001025896808 | 0.02901127158 | -0.8843383707 |
| Hid1 | -0.9921718163 | 4.06714403 | -4.210294294 | 0.001033359277 | 0.02909526979 | -0.6428008463 |
| LOC101597245 | -0.7503632345 | 4.198701663 | -4.210755334 | 0.001032475585 | 0.02909526979 | -0.6922367123 |
| St14 | -0.3820631524 | 7.069668971 | -4.208874832 | 0.001036084909 | 0.02909526979 | -0.926556248 |
| Slc39a13 | -0.3091067211 | 6.298712129 | -4.208567306 | 0.001036676391 | 0.02909526979 | -0.9370364526 |
| Maob | -1.230787681 | 2.940671955 | -4.205263338 | 0.001043053103 | 0.02910977603 | -0.5608653449 |
| C2cd4d | -0.7081180592 | 3.013492076 | -4.205786896 | 0.001042039939 | 0.02910977603 | -0.5609886539 |
| Pxylp1 | -0.7019566538 | 2.939313901 | -4.205696872 | 0.001042214077 | 0.02910977603 | -0.5729287084 |
| Jag2 | -0.5840745507 | 3.884148691 | -4.202638873 | 0.001048147159 | 0.02913601639 | -0.6619847666 |
| LOC123455359 | -0.5615296488 | 4.333128632 | -4.202873026 | 0.001047691631 | 0.02913601639 | -0.7139624452 |
| Lrba | -0.285016032 | 7.481964685 | -4.197914015 | 0.001057382765 | 0.02913601639 | -0.9244502391 |

| | | | | | | |
|---|---|---|---|---|---|---|
| Far1 | 0.3498688898 | 4.845393105 | 4.200132257 | 0.001053036404 | 0.02913601639 | -0.8636632959 |
| Zbtb18 | 0.3573188477 | 5.551767153 | 4.198157986 | 0.001056903832 | 0.02913601639 | -0.924979636 |
| Msn | 0.4100264455 | 8.04406189 | 4.200076771 | 0.001053144897 | 0.02913601639 | -0.8814388024 |
| LOC123464458 | 1.229477192 | 2.914227618 | 4.197763348 | 0.001057678646 | 0.02913601639 | -0.5965331474 |
| C12H9orf64 | 0.4027606212 | 3.675721664 | 4.196251915 | 0.001060651546 | 0.02914681633 | -0.6797336875 |
| Hsp90aa1 | 0.97630673 | 9.658722654 | 4.1955768 | 0.001061982236 | 0.02914681633 | -0.7903080509 |
| Cfd | -1.871911934 | 4.355652185 | -4.186460449 | 0.001080120455 | 0.02918302115 | -0.7408109445 |
| Nupr1 | -1.064638519 | 7.462145434 | -4.186920586 | 0.001079197346 | 0.02918302115 | -0.9487786741 |
| Mtcl1 | -0.9069551024 | 2.413724326 | -4.178317985 | 0.001096590743 | 0.02918302115 | -0.5955762646 |
| Lypd6 | -0.9012598287 | 3.469342113 | -4.179753178 | 0.001093669017 | 0.02918302115 | -0.6722414284 |
| Tagln | -0.8002790181 | 6.628903215 | -4.181134494 | 0.001090864543 | 0.02918302115 | -0.9857444428 |
| Mitf | -0.3701532402 | 6.074696752 | -4.178480818 | 0.001096258848 | 0.02918302115 | -0.9888016163 |
| Ddx51 | -0.3632843452 | 4.709239961 | -4.185912396 | 0.001081220997 | 0.02918302115 | -0.8089864804 |
| Lrp10 | -0.2788455007 | 8.149641409 | -4.193343022 | 0.001066397418 | 0.02918302115 | -0.8934789311 |
| Zbtb41 | 0.2818268814 | 5.388953723 | 4.186240092 | 0.001080562812 | 0.02918302115 | -0.9356060019 |
| Ralgapa1 | 0.3090630411 | 6.142789331 | 4.183479589 | 0.001086120266 | 0.02918302115 | -0.9849842947 |
| Map3k2 | 0.378474921 | 6.234364652 | 4.187357398 | 0.001078321783 | 0.02918302115 | -0.9779034748 |
| Aak1 | 0.5719223391 | 5.534020336 | 4.180590948 | 0.001091967214 | 0.02918302115 | -0.9750735991 |
| LOC101596333 | 0.5822814448 | 4.218342672 | 4.187075033 | 0.001078887683 | 0.02918302115 | -0.8092972736 |
| Hmbox1 | 0.7377245253 | 2.993901634 | 4.182051164 | 0.001089007524 | 0.02918302115 | -0.6718293758 |
| Esrrg | 0.936976959 | 4.520834691 | 4.186726149 | 0.001079587319 | 0.02918302115 | -0.8640291596 |
| LOC101617174 | 1.30130273 | 1.478658457 | 4.180253455 | 0.001092652451 | 0.02918302115 | -0.6071967002 |
| Steap1 | 1.89713209 | 2.489575923 | 4.188461963 | 0.001076111015 | 0.02918302115 | -0.5787360326 |
| LOC123459515 | 3.17353801 | 1.056767722 | 4.176351832 | 0.001100606439 | 0.02923767873 | -0.7743211847 |
| Klhl32 | -0.4664028453 | 5.014940337 | -4.174550993 | 0.001104297782 | 0.02928354065 | -0.9016178266 |
| LOC101607323 | -0.6021419078 | 3.199153034 | -4.173424058 | 0.001106614247 | 0.02929284568 | -0.6373776773 |
| L1cam | -0.5263774147 | 6.365798745 | -4.165463336 | 0.001123121028 | 0.02955581103 | -1.015739664 |
| Ptp4a1 | -0.5172616215 | 4.022431893 | -4.163300689 | 0.001127648981 | 0.02955581103 | -0.8044657462 |
| Slc22a15 | -0.5101852804 | 4.139736569 | -4.162971008 | 0.001128340887 | 0.02955581103 | -0.8080387986 |
| Ctps2 | -0.4816222391 | 3.972042565 | -4.161794002 | 0.001130814651 | 0.02955581103 | -0.7739090565 |
| Arglu1 | -0.4350139103 | 7.351063603 | -4.161827389 | 0.001130744402 | 0.02955581103 | -1.010144166 |
| Ap1m1 | -0.374247436 | 6.12339399 | -4.166605542 | 0.001120737139 | 0.02955581103 | -1.011184318 |
| Med11 | -0.3174077299 | 4.33933878 | -4.161034377 | 0.001132414151 | 0.02955581103 | -0.8241279526 |
| Man1a1 | 0.5871778198 | 7.960384032 | 4.167375083 | 0.001119133975 | 0.02955581103 | -0.945876757 |
| Fem1b | 0.5783247562 | 4.031639101 | 4.159746402 | 0.001135131492 | 0.02957493816 | -0.8080772265 |
| Bmp6 | -0.6936761685 | 4.742489963 | -4.155513018 | 0.001144110384 | 0.02975685349 | -0.8982721841 |
| Dr1 | 0.4038254029 | 6.162156628 | 4.154290823 | 0.001146716202 | 0.02977266822 | -1.035908059 |
| Igsf10 | -1.440253075 | 1.016774042 | -4.15282181 | 0.001149856344 | 0.02980227668 | -0.7458272571 |
| Ramac | 0.3959584529 | 5.349181545 | 4.151373602 | 0.001152960678 | 0.02983085588 | -0.974937799 |
| Tmem52b | -0.7277533114 | 6.801509962 | -4.149523024 | 0.001156940078 | 0.02988193758 | -1.043379652 |
| Gjb5 | -1.40740047 | 1.070959218 | -4.146392321 | 0.001163704421 | 0.02989296255 | -0.7474436321 |
| Adora1 | -1.043877728 | 3.222570449 | -4.145614983 | 0.001165390273 | 0.02989296255 | -0.7355148598 |
| Pde2a | -0.6022256035 | 5.37754284 | -4.146880242 | 0.001162647524 | 0.02989296255 | -0.9733587013 |
| Hmgn2 | 0.384154303 | 5.866958532 | 4.145736958 | 0.001165125572 | 0.02989296255 | -1.031429072 |
| Slc34a2 | -0.9439494389 | 5.023641017 | -4.135665716 | 0.001187191061 | 0.0299011528 | -0.9685286083 |
| Fxyd1 | -0.7753419829 | 1.355335274 | -4.144104608 | 0.001168673089 | 0.0299011528 | -0.7327471801 |
| Schip1 | -0.6035317783 | 5.081753498 | -4.138672582 | 0.001180558531 | 0.0299011528 | -0.966913708 |
| Ccpg1 | 0.2823743712 | 7.347677564 | 4.139791647 | 0.001178099849 | 0.0299011528 | -1.035718518 |
| Tmem168 | 0.321755886 | 5.795190597 | 4.136303606 | 0.001185780809 | 0.0299011528 | -1.058981258 |
| Otud4 | 0.3466628814 | 5.89479907 | 4.143154465 | 0.001170743111 | 0.0299011528 | -1.051121377 |
| C11H16orf72 | 0.3517180336 | 6.114683969 | 4.136585686 | 0.001185157735 | 0.0299011528 | -1.069624211 |
| Rab11fip4 | 0.4656538221 | 6.307981577 | 4.135399644 | 0.001187779807 | 0.0299011528 | -1.074247302 |
| Fndc3a | 0.4776371767 | 7.614664604 | 4.138722396 | 0.001180448974 | 0.0299011528 | -1.01548477 |
| Cdon | 0.4792841221 | 5.46787814 | 4.141990708 | 0.001173283663 | 0.0299011528 | -1.017202043 |
| Chtf8 | 0.6591258658 | 2.6198534 | 4.137872562 | 0.00118231948 | 0.0299011528 | -0.6948861742 |
| LOC101608894 | 0.5044840408 | 3.532847796 | 4.134443127 | 0.001189898797 | 0.02990398275 | -0.7792819938 |
| Popdc2 | -1.326796221 | 1.315258519 | -4.131676857 | 0.001196048886 | 0.0299575068 | -0.7185080218 |
| Ormdl2 | 0.3817896098 | 4.238006336 | 4.131879609 | 0.00119559701 | 0.0299575068 | -0.8669992978 |
| LOC101610073 | -1.364027056 | 1.50672824 | -4.127875229 | 0.00120455423 | 0.02996906794 | -0.7627593957 |
| Cited2 | -0.4850217731 | 6.737726281 | -4.128356337 | 0.001203474422 | 0.02996906794 | -1.083107331 |
| Btbd7 | 0.284257663 | 5.839741253 | 4.130439506 | 0.001198810395 | 0.02996906794 | -1.073751434 |
| Mob3b | 0.8744626472 | 6.098421553 | 4.128405673 | 0.001203363747 | 0.02996906794 | -1.080822298 |
| Porcn | -0.4106539384 | 3.478640358 | -4.12650452 | 0.001207636146 | 0.02997693678 | -0.7392982394 |
| Lhx1 | 0.8475229185 | 6.41615183 | 4.125946359 | 0.001208893445 | 0.02997693678 | -1.086081808 |
| Znf629 | -0.2868166722 | 5.569637533 | -4.123972186 | 0.001213351228 | 0.03001938042 | -1.051202617 |
| Edem3 | 0.5382400858 | 4.526256443 | 4.123405602 | 0.001214633724 | 0.03001938042 | -0.9959589077 |
| Dusp6 | -0.5930085686 | 7.581324403 | -4.121729067 | 0.001218436832 | 0.03006351673 | -1.058494622 |
| Tinf2 | -0.2820766236 | 5.333715343 | -4.120722719 | 0.001220725548 | 0.03007020305 | -1.024000345 |
| Tepsin | -0.4808790113 | 4.638476222 | -4.117802251 | 0.001227392575 | 0.0301348131 | -0.9117577985 |
| Rrm1 | 0.7346423839 | 5.315581898 | 4.118283732 | 0.001226290849 | 0.0301348131 | -1.054957433 |
| Vsig2 | -1.165132814 | 2.656977961 | -4.116113738 | 0.001231264297 | 0.03018015103 | -0.6977996336 |
| Lypd3 | -1.881319515 | -0.5285468744 | -4.113953747 | 0.001236235425 | 0.03021223708 | -1.34547974 |
| Diaph2 | 0.4467322509 | 4.155133047 | 4.112905614 | 0.001238655093 | 0.03021223708 | -0.9352364246 |
| Dusp14 | 0.5844002007 | 5.320501518 | 4.113297203 | 0.001237750522 | 0.03021223708 | -1.032164381 |
| Srgap3 | -0.4647430405 | 5.587130209 | -4.111797368 | 0.001241218835 | 0.03022530115 | -1.072672594 |
| G0s2 | -1.327311212 | 5.477293598 | -4.108969029 | 0.001247786487 | 0.03028625737 | -0.9975562472 |
| Golim4 | 0.7325573567 | 5.830603656 | 4.109608892 | 0.001246297548 | 0.03028625737 | -1.113822564 |
| Hspb6 | -0.8319479961 | 3.375827186 | -4.106456668 | 0.001253650383 | 0.030342692 | -0.769778597 |
| Ankrd23 | -0.6773038638 | 4.224453684 | -4.106228811 | 0.001254183606 | 0.030342692 | -0.8693063849 |
| Esam | -0.3025528916 | 5.724224577 | -4.103485583 | 0.001260621539 | 0.03043135337 | -1.103086176 |
| Cdr2 | 0.5876862114 | 6.58643934 | 4.102928848 | 0.001261932254 | 0.03043135337 | -1.132866117 |
| Svep1 | -1.307991818 | 3.690437649 | -4.101876323 | 0.001264414028 | 0.03044194227 | -0.8251056406 |
| Helz | 0.7521849098 | 4.344221655 | 4.100622739 | 0.001267376438 | 0.03046405009 | -0.9762877891 |
| Zeb2 | -0.5803239521 | 6.638770408 | -4.098817688 | 0.001271654589 | 0.03046859861 | -1.140781823 |
| Ankrd13c | 0.303142394 | 6.336255869 | 4.099273877 | 0.001270571975 | 0.03046859861 | -1.143489636 |
| Pradc1 | 0.2888772949 | 5.269362052 | 4.089657379 | 0.001293595432 | 0.03094454692 | -1.084958567 |

| | | | | | | |
|---|---|---|---|---|---|---|
| Sart3 | -0.2453303818 | 5.431493259 | -4.086099389 | 0.001302222162 | 0.03101420376 | -1.114362626 |
| Rhobtb3 | 0.492483917 | 5.351923112 | 4.085882247 | 0.001302750557 | 0.03101420376 | -1.131515198 |
| LOC101593666 | 0.5090047595 | 4.86272978 | 4.087205507 | 0.001299533943 | 0.03101420376 | -1.050666867 |
| Lmo4 | -0.4926314881 | 6.327529912 | -4.081116687 | 0.001314402766 | 0.03119194971 | -1.174749759 |
| Hoxa9 | -0.4397263262 | 5.615358634 | -4.081867837 | 0.001312559052 | 0.03119194971 | -1.114291889 |
| Trim2 | 0.9387754514 | 3.606504835 | 4.077557435 | 0.00132317529 | 0.03135020882 | -0.9205329851 |
| Rerg | -0.6418315975 | 5.170591579 | -4.075579358 | 0.001328076668 | 0.03141639141 | -1.079143953 |
| Boc | -0.4013951933 | 4.763832204 | -4.068236052 | 0.001346436075 | 0.0317974763 | -1.049797736 |
| Elp1 | 0.4506121346 | 5.036271894 | 4.067435348 | 0.001348453668 | 0.0317974763 | -1.143565048 |
| Fam3d | -1.00178539 | 2.921937186 | -4.06531792 | 0.00135380409 | 0.03187321066 | -0.8141344014 |
| Nfatc4 | -0.6886713829 | 3.830716436 | -4.063668486 | 0.001357987069 | 0.03192126387 | -0.8897457006 |
| Sptssb | -0.9679337284 | 3.578750511 | -4.060058475 | 0.001367188496 | 0.03208694513 | -0.8608040654 |
| Trim29 | -1.029515962 | 1.131004374 | -4.054088089 | 0.001382547001 | 0.03211559815 | -0.8846622373 |
| Ca8 | -0.9362462042 | 3.990409739 | -4.049576977 | 0.001394269074 | 0.03211559815 | -0.9532514983 |
| Ddx42 | -0.2767160217 | 6.577937669 | -4.05060618 | 0.001391585744 | 0.03211559815 | -1.235187257 |
| Fbh1 | 0.2814813919 | 6.028412292 | 4.05566239 | 0.001378480068 | 0.03211559815 | -1.21934688 |
| Atrx | 0.3689924435 | 6.760029419 | 4.050349846 | 0.001392253561 | 0.03211559815 | -1.224420435 |
| Nck1 | 0.3744266852 | 5.449132401 | 4.054534001 | 0.001381393817 | 0.03211559815 | -1.186452176 |
| Noc3l | 0.3791900177 | 4.638822713 | 4.05122566 | 0.001389973199 | 0.03211559815 | -1.094298509 |
| LOC101614042 | 0.4521009284 | 5.401528325 | 4.050805775 | 0.001391065977 | 0.03211559815 | -1.20557984 |
| Mknk2 | 0.5239098352 | 7.512103438 | 4.058509847 | 0.001371155346 | 0.03211559815 | -1.190958687 |
| Cdhr2 | 0.7288665935 | 7.081164659 | 4.051490009 | 0.001389285665 | 0.03211559815 | -1.212555724 |
| Slc5a2 | 0.9828848687 | 8.825323523 | 4.049830454 | 0.001393607719 | 0.03211559815 | -1.117437656 |
| LOC123454100 | 1.025775818 | 2.095165846 | 4.05343442 | 0.001384239263 | 0.03211559815 | -0.8029133781 |
| Wnt2b | -1.064650005 | 1.408946072 | -4.046236673 | 0.00140301456 | 0.03213823941 | -0.8395578565 |
| Nsun6 | -0.6520798828 | 4.267405775 | -4.045546033 | 0.001404829791 | 0.03213823941 | -1.018119336 |
| Naa60 | -0.2889540733 | 6.12860316 | -4.047858825 | 0.001398760473 | 0.03213823941 | -1.231830892 |
| Ube2m | -0.2684526163 | 6.098881997 | -4.045088186 | 0.001406034496 | 0.03213823941 | -1.237388164 |
| Abca5 | 0.2940057673 | 7.425729805 | 4.045158279 | 0.001405849996 | 0.03213823941 | -1.207899342 |
| Mapk8 | 0.698180594 | 3.006861925 | 4.042624776 | 0.001412534544 | 0.03223736954 | -0.9072108018 |
| Myoc | -2.18612738 | -0.2491313462 | -4.03923484 | 0.001421529852 | 0.03226728402 | -1.140251179 |
| Lgr6 | -0.9615100115 | 2.395202274 | -4.039530707 | 0.001420742421 | 0.03226728402 | -0.8289019053 |
| Rassf7 | -0.4733247537 | 5.427865004 | -4.039151603 | 0.00142175146 | 0.03226728402 | -1.169636118 |
| Lix1l | 0.5768002235 | 3.283824147 | 4.038868331 | 0.001422505912 | 0.03226728402 | -0.9030919677 |
| Nbl1 | -1.59591627 | 4.003675253 | -4.033737609 | 0.00143624217 | 0.03243088962 | -0.9906688128 |
| Lpl | -0.8653900656 | 9.401101722 | -4.033267779 | 0.001437506813 | 0.03243088962 | -1.113504118 |
| Polm | -0.8373422109 | 2.145178218 | -4.032927668 | 0.001438423005 | 0.03243088962 | -0.8389523705 |
| Elk4 | 0.5081043296 | 4.765011458 | 4.033078605 | 0.001438016337 | 0.03243088962 | -1.173396134 |
| Slc11a2 | 0.4306376958 | 4.645045173 | 4.02843166 | 0.001450590863 | 0.03265582423 | -1.135094362 |
| Fam241a | -0.4972108219 | 4.968512879 | -4.027326161 | 0.001453598903 | 0.0326741847 | -1.158787473 |
| Tulp4 | 0.4483886811 | 5.842853215 | 4.025324567 | 0.001459061511 | 0.03274758087 | -1.268829283 |
| S100a6 | -1.043304961 | 5.182418868 | -4.019236489 | 0.001475806544 | 0.03307360139 | -1.188939312 |
| Ier3 | -0.5388870708 | 6.178210169 | -4.018417789 | 0.001478073347 | 0.0330746653 | -1.284202561 |
| Il13ra1 | 0.4318132457 | 5.778124834 | 4.017386292 | 0.00148093443 | 0.03308900422 | -1.280865183 |
| Rassf10 | -0.7689531227 | 3.540991268 | -4.01192284 | 0.001496183577 | 0.03337967641 | -0.9800005429 |
| Rarres1 | -0.5518946719 | 4.420182314 | -4.00908527 | 0.001504167066 | 0.03349799679 | -1.104674531 |
| Nufip2 | 0.5200110779 | 5.545161659 | 4.008442152 | 0.001505982544 | 0.03349799679 | -1.277956607 |
| Cant1 | -0.491075986 | 4.161984991 | -4.005863199 | 0.001513285368 | 0.03361027101 | -1.080063983 |
| Dusp4 | -1.322443823 | 3.394110307 | -4.001565864 | 0.001525534968 | 0.03378164581 | -0.9970935449 |
| Gpm6a | 0.7085151537 | 5.470593247 | 4.001960369 | 0.001524406198 | 0.03378164581 | -1.298967874 |
| Nptxr | -1.156669137 | 2.734228683 | -3.999147454 | 0.001532473365 | 0.03385673172 | -0.9044435832 |
| Kdm5a | 0.3587521357 | 5.69161627 | 3.998801217 | 0.001533469363 | 0.03385673172 | -1.314012444 |
| LOC123463614 | -0.9819547984 | 3.365022489 | -3.994491597 | 0.001545922289 | 0.03393060365 | -0.9448123722 |
| Hacd1 | -0.6856242628 | 4.226915258 | -3.994969319 | 0.001544536784 | 0.03393060365 | -1.072598858 |
| Tesk1 | -0.4294431616 | 5.247877231 | -3.996826507 | 0.001539162591 | 0.03393060365 | -1.240012055 |
| Sec31a | 0.2712934949 | 6.948071811 | 3.994534951 | 0.0015457965 | 0.03393060365 | -1.329332576 |
| Pspc1 | -0.2654254225 | 5.837929504 | -3.990152387 | 0.001558565562 | 0.03415779789 | -1.332066187 |
| Arc | -1.46609348 | 1.621083781 | -3.986249196 | 0.001570029005 | 0.03430812648 | -0.9211728837 |
| Mfsd10 | -0.4398157553 | 4.029023523 | -3.986779462 | 0.001568466583 | 0.03430812648 | -1.084005744 |
| Spag1 | -0.8715440664 | 3.817774803 | -3.984018908 | 0.001576618022 | 0.03440166673 | -1.060795611 |
| Rasa2 | 0.4211813744 | 5.130373373 | 3.980948836 | 0.001585734465 | 0.03455000106 | -1.291296578 |
| Pear1 | -0.641850139 | 3.788027223 | -3.977098729 | 0.001597243647 | 0.03469930331 | -1.074632312 |
| Mafg | 0.5197908013 | 5.679572429 | 3.977246956 | 0.001596798968 | 0.03469930331 | -1.343243196 |
| Btnl9 | -2.070329568 | 0.7924488921 | -3.976100458 | 0.001600241754 | 0.03470549154 | -1.045438393 |
| Fus | -0.4857993184 | 7.268523314 | -3.975454108 | 0.001602186015 | 0.03470549154 | -1.358137426 |
| Slco2a1 | 1.199003272 | 2.799771718 | 3.973085043 | 0.001609333046 | 0.03480971029 | -0.9682677362 |
| Fjx1 | -0.9706772843 | 2.600064456 | -3.970792827 | 0.001616279354 | 0.03485877166 | -0.9504523141 |
| Ostc | 0.3098290128 | 5.928180372 | 3.971128171 | 0.001615261216 | 0.03485877166 | -1.367848006 |
| Fasn | -1.120328089 | 5.387556819 | -3.967219204 | 0.001627170213 | 0.0348688654 | -1.305413932 |
| Actn1 | -0.4050736209 | 5.80463406 | -3.965275477 | 0.001633125414 | 0.0348688654 | -1.36698118 |
| Setx | 0.3437363897 | 6.043742363 | 3.969402842 | 0.001620506521 | 0.0348688654 | -1.381890075 |
| Rb1cc1 | 0.3760356957 | 6.756378904 | 3.966670165 | 0.0016288501 | 0.0348688654 | -1.385466972 |
| Rtn4 | 0.39011248 | 9.151654275 | 3.968244934 | 0.001624036555 | 0.0348688654 | -1.244601465 |
| Creb1 | 0.4453770393 | 5.432773722 | 3.965382477 | 0.001632797006 | 0.0348688654 | -1.351193964 |
| Cd47 | 0.4996605751 | 5.667956113 | 3.967354365 | 0.001626756934 | 0.0348688654 | -1.358498337 |
| Wdfy1 | 0.4971085834 | 4.242705744 | 3.964327274 | 0.001636038628 | 0.03488109253 | -1.217993366 |
| Pfkl | -0.4660462056 | 6.455010987 | -3.960975191 | 0.001646380189 | 0.03500143218 | -1.401331263 |
| Zscan26 | 0.5145828918 | 5.353826499 | 3.961220545 | 0.001645620975 | 0.03500143218 | -1.336127794 |
| Lamc2 | -0.5190972989 | 6.354830346 | -3.959612 | 0.001650604936 | 0.03502744061 | -1.40419025 |
| Invs | 0.4223919834 | 4.519488822 | 3.958308641 | 0.001654654646 | 0.03502744061 | -1.239650865 |
| Avl9 | 0.4562606513 | 5.115904009 | 3.958613475 | 0.001653706574 | 0.03502744061 | -1.356588595 |
| Nudt21 | 0.2640868904 | 5.651334955 | 3.957443968 | 0.001657346918 | 0.03503466826 | -1.377165355 |
| Fos | -1.268912666 | 2.22793295 | -3.949199851 | 0.001683242646 | 0.03533150022 | -0.9814777493 |
| Spaca6 | -0.7127430692 | 3.082964926 | -3.950240849 | 0.001679950006 | 0.03533150022 | -0.9985816685 |
| Prkag2 | -0.4350667273 | 6.261512411 | -3.949554919 | 0.001682118835 | 0.03533150022 | -1.420005117 |
| Ripor1 | -0.3061181902 | 7.595438959 | -3.949827902 | 0.001681255351 | 0.03533150022 | -1.392214544 |

| | | | | | | |
|---|---|---|---|---|---|---|
| Npl | 0.7394597393 | 5.88602497 | 3.951364896 | 0.001676402093 | 0.03533150022 | -1.399800894 |
| LOC101598203 | -1.39259973 | -0.119191561 | -3.948408151 | 0.001685751194 | 0.03533438825 | -1.211044685 |
| Trmt2a | -0.3618215111 | 4.925288639 | -3.946369468 | 0.00169222853 | 0.03542033959 | -1.285832903 |
| Sesn3 | -0.6462359457 | 6.426073657 | -3.943060369 | 0.001702796601 | 0.03554908739 | -1.433256871 |
| Chordc1 | 0.9929049578 | 6.473273426 | 3.942949993 | 0.001703150265 | 0.03554908739 | -1.42747939 |
| Vav3 | 0.3108442943 | 7.075890286 | 3.940802817 | 0.001710045195 | 0.03564308188 | -1.422508157 |
| C2cd2l | -0.2936359145 | 5.035989778 | -3.939531231 | 0.001714141942 | 0.03567857173 | -1.331509345 |
| Mroh6 | -0.7182712553 | 4.206764485 | -3.935873869 | 0.001725981236 | 0.03587489311 | -1.146291839 |
| Syn2 | 1.897589794 | 2.335217133 | 3.934469326 | 0.00173055014 | 0.03591976147 | -1.00236319 |
| Fam13c | -0.5839133172 | 2.783548855 | -3.93274914 | 0.001736162693 | 0.03598613715 | -1.039986436 |
| Chkb | -0.4541843233 | 5.266194086 | -3.930662077 | 0.001742997328 | 0.036030447 | -1.355210632 |
| Rfx3 | 0.543024586 | 1.932483613 | 3.93061989 | 0.001743135764 | 0.036030447 | -1.020627907 |
| Rundc3b | 0.3813118324 | 4.458552657 | 3.929127417 | 0.001748040554 | 0.03608178446 | -1.290837907 |
| LOC101593520 | -0.3110626211 | 6.132534001 | -3.925649481 | 0.001759525201 | 0.03626860868 | -1.458708927 |
| Gramd4 | 0.4108239392 | 5.417046733 | 3.92293629 | 0.001768538198 | 0.03640403973 | -1.427595154 |
| Ercc5 | -0.2938875173 | 5.217733999 | -3.915076058 | 0.001794917103 | 0.03664335559 | -1.390835322 |
| Nup155 | 0.3151073925 | 4.911034546 | 3.915403895 | 0.00179380887 | 0.03664335559 | -1.391324598 |
| Stk4 | 0.460101077 | 5.174428028 | 3.916225355 | 0.001791035054 | 0.03664335559 | -1.432849693 |
| Tet3 | 0.6793150724 | 3.770601208 | 3.915712717 | 0.001792765559 | 0.03664335559 | -1.232469152 |
| Mapk6 | 1.178661769 | 5.892235775 | 3.915744433 | 0.001792658446 | 0.03664335559 | -1.464182774 |
| Rora | 1.339625615 | 2.7105533 | 3.916597147 | 0.001789781073 | 0.03664335559 | -1.100670707 |
| Ireb2 | 0.6349413885 | 5.995836507 | 3.914127552 | 0.001798127415 | 0.03665867698 | -1.487618809 |
| LOC123457702 | -0.8719453572 | 1.353876006 | -3.912574068 | 0.001803398044 | 0.03666581317 | -1.060742973 |
| LOC123456050 | -0.7989633814 | 3.306433941 | -3.913182729 | 0.001801331109 | 0.03666581317 | -1.124519687 |
| Synj1 | 0.2980999292 | 6.044885741 | 3.908835231 | 0.001816148042 | 0.03687473333 | -1.494359068 |
| LOC123463457 | 2.201269641 | 1.017248948 | 3.907620964 | 0.001820308699 | 0.03690892591 | -1.137921852 |
| Adamts1 | -0.5402897246 | 6.311383838 | -3.906082869 | 0.001825592964 | 0.03692682629 | -1.504123138 |
| Rhbdf1 | -0.3799509697 | 5.456827529 | -3.905921825 | 0.00182614715 | 0.03692682629 | -1.432322557 |
| Sgpp2 | -0.8254578524 | 3.06467732 | -3.905196313 | 0.001828645939 | 0.0369272499 | -1.131816844 |
| Ampd3 | -0.7255328018 | 5.795314066 | -3.904250877 | 0.001831907434 | 0.03694305343 | -1.463842038 |
| LOC101617303 | 1.233887156 | -0.3076270711 | 3.902353034 | 0.001838472461 | 0.03702534472 | -1.415905553 |
| Flna | -0.4373342738 | 7.45412058 | -3.90139925 | 0.001841780884 | 0.03704191703 | -1.483890782 |
| Rpia | -0.4935286261 | 4.941851624 | -3.900447471 | 0.00184508843 | 0.03705842705 | -1.400709543 |
| LOC101603506 | -0.6272798186 | 1.758092323 | -3.892876379 | 0.001871616378 | 0.03743986427 | -1.082882011 |
| LOC123463360 | -0.4969451278 | 5.717694283 | -3.893514648 | 0.001869364988 | 0.03743986427 | -1.471542545 |
| Fbxo10 | 1.209948063 | 4.774779818 | 3.893968816 | 0.001867764671 | 0.03743986427 | -1.360531279 |
| Krt20 | -1.005519101 | 1.362794668 | -3.890578317 | 0.001879745423 | 0.03750180192 | -1.106472006 |
| LOC101598985 | 0.8318704348 | 2.618209376 | 3.890735615 | 0.001879187855 | 0.03750180192 | -1.116658326 |
| Pknox2 | -0.6401071272 | 2.379362753 | -3.882368833 | 0.001909081511 | 0.03783383214 | -1.098264491 |
| Hmbs | -0.3516767061 | 5.174653643 | -3.883237852 | 0.001905954077 | 0.03783383214 | -1.449846338 |
| Laptm4a | 0.3326698808 | 8.151075853 | 3.883742506 | 0.00190414033 | 0.03783383214 | -1.471543822 |
| Tmem150a | 0.7833415172 | 6.277179833 | 3.882559341 | 0.001908395458 | 0.03783383214 | -1.538924224 |
| Klhl31 | 2.203835117 | 3.387119372 | 3.883075468 | 0.001906538067 | 0.03783383214 | -1.198131588 |
| Trmt9b | 0.7558415772 | 3.699172177 | 3.881345017 | 0.001912772766 | 0.03785664347 | -1.284235959 |
| Hnrnpd | -0.4026303642 | 6.487743343 | -3.878415927 | 0.001923373668 | 0.03796561295 | -1.55733419 |
| Ube3a | 0.3984373231 | 5.441816921 | 3.878555096 | 0.001922868631 | 0.03796561295 | -1.521957353 |
| Slc17a3 | -0.4732257292 | 7.57717247 | -3.874956543 | 0.001935971253 | 0.03811331516 | -1.522932928 |
| Nktr | -0.3667474195 | 6.096977026 | -3.875486643 | 0.001934035396 | 0.03811331516 | -1.548086281 |
| LOC123463346 | 0.6040455855 | 5.266579413 | 3.87355658 | 0.00194109328 | 0.03816373767 | -1.498835399 |
| Ren | -1.431908072 | 5.354755312 | -3.871222454 | 0.001949663963 | 0.03826875396 | -1.428900838 |
| Ltbp3 | -0.4502745014 | 6.248018028 | -3.870704698 | 0.001951570355 | 0.03826875396 | -1.5658496 |
| Sft2d2 | 0.4494305114 | 5.573458588 | 3.869838067 | 0.001954765578 | 0.03828103996 | -1.551517266 |
| Abcg1 | -0.97068478 | 4.147792542 | -3.868850506 | 0.001958413191 | 0.03830214145 | -1.293569004 |
| Ccdc40 | -1.193805207 | 0.810070445 | -3.867525949 | 0.00196331645 | 0.03834771304 | -1.20782813 |
| Cavin2 | 0.480245313 | 6.191252873 | 3.865868677 | 0.001969469045 | 0.03841753557 | -1.578742529 |
| Mphosph9 | 0.5217222993 | 2.561806049 | 3.864743427 | 0.001973657754 | 0.038448917 | -1.161770551 |
| Cavin1 | -0.5594940266 | 6.348424561 | -3.862997008 | 0.001980176804 | 0.03847532583 | -1.580727367 |
| Tra2b | 0.4147312164 | 6.376741724 | 3.863437619 | 0.001978530011 | 0.03847532583 | -1.585299667 |
| LOC101606252 | 1.632778649 | 4.018318994 | 3.862110771 | 0.001983493376 | 0.03848958566 | -1.30814121 |
| Alkbh7 | -0.5702163208 | 5.042630468 | -3.860875334 | 0.001988126243 | 0.03852931782 | -1.472555631 |
| Dusp1 | -0.545578022 | 4.249861369 | -3.858731721 | 0.001996191035 | 0.03863537013 | -1.373865 |
| Znf608 | -0.5495753459 | 4.080681686 | -3.852935175 | 0.002018167014 | 0.03898974104 | -1.35148429 |
| Ap1b1 | -0.34244589 | 6.860006553 | -3.851839369 | 0.00202234918 | 0.03898974104 | -1.602981401 |
| Hif1an | 0.3341212741 | 6.572588592 | 3.852130542 | 0.002021237047 | 0.03898974104 | -1.605697979 |
| Susd2 | -0.9342006914 | 7.094732061 | -3.844743457 | 0.002049646202 | 0.03926205315 | -1.603417564 |
| Fam83f | -0.7358446203 | 3.566636779 | -3.845204015 | 0.002047863117 | 0.03926205315 | -1.265556186 |
| Dusp8 | -0.6905080106 | 3.131439562 | -3.846314942 | 0.002043568611 | 0.03926205315 | -1.21494433 |
| Gtf2e1 | 0.3860834946 | 4.893839723 | 3.845851473 | 0.002045359121 | 0.03926205315 | -1.499476345 |
| Snx8 | 0.4295978898 | 5.234372639 | 3.845666089 | 0.002046075758 | 0.03926205315 | -1.548494102 |
| Iqub | -1.327053236 | -0.1628173974 | -3.843567417 | 0.002054206526 | 0.03926771881 | -1.380122112 |
| Mapkap1 | 0.2342877281 | 6.006538244 | 3.843308545 | 0.002055211747 | 0.03926771881 | -1.618306532 |
| C8H15orf48 | -1.838139225 | 1.019929306 | -3.841252297 | 0.00206321419 | 0.03937014222 | -1.271163275 |
| Tinagl1 | -0.5090821358 | 6.903461325 | -3.839178793 | 0.002071316046 | 0.03947419825 | -1.630494773 |
| LOC123461566 | -0.9227427709 | 1.842401081 | -3.834623613 | 0.00208922901 | 0.03951241109 | -1.173326193 |
| Sh3yl1 | -0.3472052058 | 5.379592487 | -3.836518506 | 0.002081758303 | 0.03951241109 | -1.565848662 |
| Sart1 | -0.2463606324 | 6.323068122 | -3.835829219 | 0.002084472687 | 0.03951241109 | -1.636300369 |
| Med1 | 0.3612143425 | 5.129528164 | 3.83549948 | 0.002085772461 | 0.03951241109 | -1.57497029 |
| Zbtb6 | 0.4293240176 | 4.584642191 | 3.837150918 | 0.002079271074 | 0.03951241109 | -1.470640195 |
| Mob3a | 0.5574404565 | 3.155460601 | 3.834977964 | 0.002087829881 | 0.03951241109 | -1.287569202 |
| Loxl4 | -1.671035009 | 5.524026157 | -3.832424483 | 0.002097933546 | 0.0395700764 | -1.552283091 |
| Ncoa5 | 0.2387346771 | 5.020915643 | 3.832842973 | 0.002096274238 | 0.0395700764 | -1.541036988 |
| Mlec | 0.4595102834 | 8.310203081 | 3.831842431 | 0.002100243604 | 0.0395700764 | -1.550782596 |
| Mss51 | -0.9708465419 | 2.050605603 | -3.830523967 | 0.002105485971 | 0.03958193742 | -1.178462582 |
| Pdzd8 | 0.4268498345 | 5.071489231 | 3.830348391 | 0.002106185088 | 0.03958193742 | -1.584457188 |
| Prpf18 | 0.4270576218 | 5.318215466 | 3.829355669 | 0.002110142431 | 0.03960636354 | -1.606180945 |
| Epha2 | -0.7497765474 | 3.945153956 | -3.825947034 | 0.00212378839 | 0.03976233464 | -1.359286278 |

| | | | | | | |
|---|---|---|---|---|---|---|
| Hif1a | -0.4125423 | 7.430764059 | -3.826126239 | 0.002123068729 | 0.03976233464 | -1.621542193 |
| LOC101610854 | -1.433245214 | 1.256605602 | -3.818431629 | 0.002154194672 | 0.04023699226 | -1.228786409 |
| Klhl24 | 0.6183383199 | 6.521026567 | 3.818346731 | 0.002154540685 | 0.04023699226 | -1.663986747 |
| Ap2a1 | -0.293472387 | 5.670191031 | -3.816293788 | 0.002162925068 | 0.04034301914 | -1.633126666 |
| Sgk1 | 0.9316521996 | 8.702649196 | 3.81523243 | 0.002167272803 | 0.04037358323 | -1.582823539 |
| Upk1b | -1.22700797 | 2.447117775 | -3.812984076 | 0.002176512444 | 0.04039422784 | -1.209741211 |
| Josd1 | -0.4282725442 | 6.439561485 | -3.81313607 | 0.002175886557 | 0.04039422784 | -1.678888783 |
| Hspa4l | 0.6164753419 | 7.299777756 | 3.814197763 | 0.002171519794 | 0.04039422784 | -1.644998495 |
| LOC123462327 | -0.9626491611 | 2.213286715 | -3.811295378 | 0.002183478627 | 0.04047311192 | -1.208135228 |
| Tab1 | -0.5001317979 | 4.095581098 | -3.807503614 | 0.002199203434 | 0.0406130468 | -1.422896716 |
| Hectd3 | -0.3080343091 | 6.464468063 | -3.808189875 | 0.002196348895 | 0.0406130468 | -1.691028225 |
| Clstn1 | -0.2691570335 | 7.275819225 | -3.807989591 | 0.002197181594 | 0.0406130468 | -1.67275516 |
| LOC123462202 | -0.9217671247 | 0.4916444274 | -3.805820156 | 0.002206221924 | 0.04065862799 | -1.319928097 |
| Mdm4 | -0.4242157952 | 6.148883761 | -3.805603199 | 0.002207128098 | 0.04065862799 | -1.679233451 |
| Syt7 | -1.3558861 | 2.136300876 | -3.802025379 | 0.002222126718 | 0.04088438824 | -1.224295449 |
| Tanc2 | 0.9389692425 | 8.050382306 | 3.801010987 | 0.002226398073 | 0.04091246669 | -1.620672262 |
| Slc36a4 | -0.4519436015 | 4.065161042 | -3.798697801 | 0.002236169718 | 0.04104142525 | -1.498378258 |
| Ampd2 | -0.5200966018 | 5.954293664 | -3.796099136 | 0.002247199563 | 0.04104868923 | -1.698227941 |
| Slc45a3 | -0.5120705186 | 5.914245953 | -3.796470707 | 0.002245619058 | 0.04104868923 | -1.700416511 |
| Asl | -0.4123771258 | 7.750876314 | -3.795449472 | 0.002249965682 | 0.04104868923 | -1.667593308 |
| Esyt2 | 0.3358585402 | 7.073897532 | 3.796089189 | 0.002247241888 | 0.04104868923 | -1.699780323 |
| Ipmk | 0.5339926201 | 4.23938538 | 3.795362218 | 0.002250337455 | 0.04104868923 | -1.551778599 |
| LOC123460879 | 0.6397926655 | 1.418162132 | 3.794554165 | 0.002253783393 | 0.04106128838 | -1.242312179 |
| Isyna1 | -0.515307021 | 5.72689593 | -3.793494196 | 0.002258311797 | 0.04109355398 | -1.678660047 |
| Lrch4 | -0.4767348944 | 6.79450354 | -3.792293751 | 0.002263451566 | 0.04113685205 | -1.719422847 |
| Tlr4 | -1.049076512 | 4.85285819 | -3.791409697 | 0.002267244321 | 0.04115559332 | -1.575788978 |
| Fkbpl | -0.4197728968 | 3.73616582 | -3.790735537 | 0.002270140953 | 0.04115804212 | -1.394823164 |
| Hcn3 | -1.481847442 | 4.746161297 | -3.789132908 | 0.002277042074 | 0.04118295877 | -1.474906668 |
| LOC101610855 | 0.4926143309 | 3.72195139 | 3.789499053 | 0.002275463523 | 0.04118295877 | -1.437187417 |
| LOC123458855 | -0.4577623498 | 8.701911894 | -3.787481637 | 0.002284175054 | 0.04126189192 | -1.612655327 |
| Ufl1 | 0.260561937 | 5.15014862 | 3.78519207 | 0.002294103036 | 0.04134101275 | -1.669817562 |
| Sost | 1.468824831 | 0.6473083032 | 3.785672348 | 0.002292016818 | 0.04134101275 | -1.341675781 |
| Mettl1 | 0.6261914846 | 3.528700743 | 3.783424922 | 0.002301795826 | 0.04142954492 | -1.403161477 |
| Syndig1 | -1.56487365 | 0.2129302921 | -3.781485035 | 0.002310270905 | 0.04148188832 | -1.490813819 |
| Slc25a44 | 0.4791718634 | 4.781152611 | 3.782022177 | 0.00230792103 | 0.04148188832 | -1.642820672 |
| LOC123462992 | -0.7250614202 | 2.300014135 | -3.777436906 | 0.002328059319 | 0.04165540847 | -1.272423829 |
| Tspyl2 | -0.5383708099 | 4.452809126 | -3.778436661 | 0.002323653207 | 0.04165540847 | -1.51254384 |
| LOC101606776 | 0.4503404178 | 3.603261328 | 3.777377783 | 0.002328320154 | 0.04165540847 | -1.457915876 |
| Nyap1 | -0.7637535026 | 3.396202182 | -3.776498159 | 0.002332204318 | 0.04167486925 | -1.353310284 |
| Akap5 | -1.097440967 | -0.2704990265 | -3.772642307 | 0.002349308831 | 0.04193023893 | -1.584566245 |
| Srpx2 | -1.590987717 | 1.06795328 | -3.764370923 | 0.002386433732 | 0.04254189224 | -1.344356361 |
| LOC101598267 | 0.4407072706 | 4.00695362 | 3.762843212 | 0.002393355856 | 0.04261431579 | -1.54082502 |
| Mia2 | 0.3417274892 | 8.095151036 | 3.761429333 | 0.002399780454 | 0.0426777185 | -1.696941318 |
| LOC123455744 | -2.265576669 | -0.03816295013 | -3.753961447 | 0.0024340072 | 0.04309598594 | -1.439922709 |
| Selenoo | -0.285679901 | 5.909746296 | -3.755535275 | 0.002426752871 | 0.04309598594 | -1.775123649 |
| Klhl15 | 0.6249255653 | 1.730747943 | 3.754935726 | 0.002429513798 | 0.04309598594 | -1.315161312 |
| Kiaa0930 | 0.6481357139 | 8.242889329 | 3.75377527 | 0.002434866816 | 0.04309598594 | -1.706247794 |
| Lats2 | 0.3942031915 | 5.998696647 | 3.752698835 | 0.002439843013 | 0.04309860455 | -1.79165715 |
| Ubr1 | 0.4677770271 | 5.205266841 | 3.752492372 | 0.002440798647 | 0.04309860455 | -1.763932869 |
| Angpt2 | -0.7055246518 | 5.493814937 | -3.7514657 | 0.002445556361 | 0.0431315106 | -1.735980137 |
| Atp8b1 | -0.3564039997 | 6.191634431 | -3.746228174 | 0.002469975256 | 0.04340806749 | -1.802386229 |
| Hoxb9 | -0.3459747593 | 5.367627134 | -3.747345018 | 0.002464747425 | 0.04340806749 | -1.724189927 |
| Pwwp3b | 0.5223550331 | 4.562700719 | 3.746734571 | 0.002467603465 | 0.04340806749 | -1.62516414 |
| LOC101610537 | -1.344802696 | 4.541686484 | -3.745386934 | 0.00247392048 | 0.04342619189 | -1.681526788 |
| Klf15 | 0.9244675461 | 7.032215575 | 3.744633461 | 0.002477459551 | 0.04343715257 | -1.781027849 |
| Msantd3 | -0.7404270651 | 2.603462422 | -3.741948872 | 0.002490111117 | 0.04355648588 | -1.361343934 |
| Lmtk2 | 0.3468066325 | 7.48400457 | 3.742300472 | 0.002488450402 | 0.04355648588 | -1.763869664 |
| Hsd17b2 | 0.4842209548 | 7.251249027 | 3.73984933 | 0.00250005144 | 0.04367909334 | -1.796050783 |
| Lbr | 0.958743801 | 5.049416626 | 3.739095393 | 0.002503630836 | 0.04369041025 | -1.715231266 |
| Mob1b | 0.3979525071 | 6.385241952 | 3.738262772 | 0.002507589863 | 0.04370831781 | -1.821569701 |
| Mgam | 1.775938443 | 4.554588225 | 3.736516287 | 0.002515914973 | 0.04380219725 | -1.687842228 |
| Coro6 | -1.150007263 | 4.818597567 | -3.735101392 | 0.002522680119 | 0.04381760119 | -1.638119917 |
| Myh10 | -0.3520715332 | 7.70419438 | -3.735273133 | 0.002521857974 | 0.04381760119 | -1.780118375 |
| LOC123453637 | -4.318046664 | -1.693165757 | -3.728908657 | 0.002552508824 | 0.04423260349 | -1.788630735 |
| Gins3 | -1.375045677 | 2.453248508 | -3.729469199 | 0.002549794115 | 0.04423260349 | -1.361387209 |
| C5H1orf198 | -0.5583912951 | 5.002778412 | -3.726947686 | 0.002562029039 | 0.04424324307 | -1.72081341 |
| Spag17 | 0.7852756948 | 4.488419578 | 3.728169572 | 0.002556092731 | 0.04424324307 | -1.609787867 |
| Enpp2 | 0.8036807907 | 7.383634441 | 3.727307184 | 0.002560281026 | 0.04424324307 | -1.794841932 |
| Tmx1 | 0.2998304225 | 6.496583132 | 3.72516216 | 0.00257072897 | 0.04434209935 | -1.846187398 |
| Phf1 | -0.3331457455 | 5.509242358 | -3.72379033 | 0.002577433626 | 0.04435507313 | -1.781685904 |
| Gdap2 | 0.3347353113 | 5.299994399 | 3.724249213 | 0.002575188908 | 0.04435507313 | -1.791331578 |
| LOC101611160 | 0.5470224511 | 5.234369473 | 3.720673684 | 0.002592732178 | 0.0445668831 | -1.833987712 |
| Herpud1 | 0.9195576247 | 7.46432808 | 3.718714212 | 0.002602397905 | 0.04463007592 | -1.81222373 |
| LOC101608141 | 4.219606646 | 6.847508038 | 3.719266459 | 0.002599670061 | 0.04463007592 | -1.669879133 |
| LOC101610994 | -0.3735939036 | 6.960667885 | -3.715152975 | 0.002620058968 | 0.04488130896 | -1.860734628 |
| Errfi1 | 1.4041368 | 5.994950946 | 3.711743624 | 0.002637081211 | 0.04512103477 | -1.845656407 |
| Loxl3 | -0.4576949439 | 6.04069642 | -3.709597216 | 0.002647855573 | 0.04525343073 | -1.848603932 |
| Cdh22 | -3.519542509 | -0.4956775638 | -3.707868879 | 0.002656563924 | 0.04526712291 | -1.830312932 |
| St3gal3 | -0.3908663092 | 3.669921195 | -3.707028944 | 0.002660806527 | 0.04526712291 | -1.546079487 |
| Poldip3 | -0.2830806313 | 7.070282017 | -3.70745655 | 0.00265864578 | 0.04526712291 | -1.872101722 |
| Egln3 | -0.2626135447 | 7.356160039 | -3.707865553 | 0.002656580709 | 0.04526712291 | -1.854079384 |
| LOC101607370 | -0.5362684738 | 4.959169175 | -3.703800654 | 0.002677177318 | 0.04539018608 | -1.74313923 |
| Anxa11 | -0.3363804929 | 8.353491117 | -3.704835164 | 0.002671920136 | 0.04539018608 | -1.801349183 |
| Entpd6 | -0.3141075888 | 5.049680091 | -3.704206379 | 0.002675114244 | 0.04539018608 | -1.78687677 |
| Uros | -0.3401317744 | 6.431633344 | -3.702237294 | 0.002685142016 | 0.0454734903 | -1.89027611 |
| Ltv1 | -0.3666833974 | 4.476505431 | -3.697438536 | 0.002709740846 | 0.04583798845 | -1.691482558 |

| | | | | | | |
|---|---|---|---|---|---|---|
| Ppp1r1a | -0.4359542438 | 8.054435146 | -3.696285501 | 0.00271568552 | 0.04584615235 | -1.839798901 |
| LOC101599550 | 2.409449501 | -0.0458150651 | 3.696151728 | 0.002716376067 | 0.04584615235 | -1.663450664 |
| Vcl | -0.3292060644 | 6.466445394 | -3.695280857 | 0.002720875962 | 0.04587015211 | -1.904282555 |
| Acot12 | 1.051206611 | 5.650858092 | 3.692966494 | 0.002732871487 | 0.04602032064 | -1.881242024 |
| Pitpnm2 | -0.3774183551 | 5.728617592 | -3.691152668 | 0.002742310315 | 0.04602325521 | -1.881957765 |
| Ints14 | 0.2671895417 | 5.307180421 | 3.691686834 | 0.002739527167 | 0.04602325521 | -1.852320729 |
| Bdp1 | 0.3445588865 | 5.433139167 | 3.691995126 | 0.002737922189 | 0.04602325521 | -1.863119494 |
| Akap11 | 0.488446851 | 7.769010101 | 3.6873933 | 0.002761979241 | 0.04630121106 | -1.866172971 |
| Dnase1 | -2.16437597 | 9.881315297 | -3.685234138 | 0.002773340804 | 0.04643943595 | -1.694595285 |
| LOC123463453 | -1.377354401 | 0.4961193741 | -3.682803677 | 0.002786186888 | 0.04654993631 | -1.526454045 |
| Inpp1 | -0.3880843429 | 4.323509561 | -3.683044183 | 0.002784913007 | 0.04654993631 | -1.683341997 |
| Kcna1 | 1.485633682 | 1.016960886 | 3.682078466 | 0.002790031674 | 0.04656197317 | -1.456038948 |
| Pbx2 | -0.3165433454 | 5.31822634 | -3.680449718 | 0.002798686366 | 0.04665416433 | -1.850002267 |
| Atxn1l | 0.354077794 | 5.525489748 | 3.674280825 | 0.002831714575 | 0.04709937758 | -1.904912486 |
| Mael | 1.953010763 | -0.8699910074 | 3.67445886 | 0.002830755839 | 0.04709937758 | -1.794402346 |
| Tsku | 1.017848521 | 5.282980487 | 3.670884075 | 0.002850069832 | 0.04735182911 | -1.846192588 |
| LOC123460593 | 1.068921709 | 1.64626157 | 3.670105294 | 0.002854295218 | 0.04736922231 | -1.452040774 |
| Pygb | -0.4068862796 | 6.271024677 | -3.665199628 | 0.002881058572 | 0.04770712878 | -1.959318494 |
| Upf2 | 0.3338576015 | 5.796458269 | 3.665471532 | 0.002879568511 | 0.04770712878 | -1.942217509 |
| Cmtr2 | 0.3506901189 | 4.69886972 | 3.66415447 | 0.002886793454 | 0.04774903756 | -1.826182983 |
| Gcat | -0.7684942243 | 5.224457996 | -3.657484807 | 0.002923665108 | 0.04780755131 | -1.860307094 |
| Cbx4 | -0.743215281 | 3.800564241 | -3.660158503 | 0.002908827017 | 0.04780755131 | -1.630554318 |
| Cd82 | -0.4412246626 | 8.503478836 | -3.659755062 | 0.002911061059 | 0.04780755131 | -1.871538898 |
| Tns2 | -0.4124570096 | 8.108316197 | -3.658903547 | 0.002915782026 | 0.04780755131 | -1.915655712 |
| Bnip2 | -0.3284408752 | 6.091783172 | -3.656884322 | 0.002927008165 | 0.04780755131 | -1.967173824 |
| Ldb1 | -0.2594206306 | 6.199738096 | -3.662335468 | 0.002896802209 | 0.04780755131 | -1.956160808 |
| Spin1 | 0.268663432 | 6.795427456 | 3.656617923 | 0.002928492523 | 0.04780755131 | -1.970775389 |
| Xpo1 | 0.3388663557 | 6.512211441 | 3.66027689 | 0.002908171785 | 0.04780755131 | -1.969598279 |
| Stradb | 0.3541312894 | 5.87233576 | 3.655982896 | 0.002932033946 | 0.04780755131 | -1.966968969 |
| Zfyve26 | 0.3735664284 | 5.543667336 | 3.658580557 | 0.002917574779 | 0.04780755131 | -1.934650134 |
| Tmf1 | 0.4154369389 | 5.754742791 | 3.661778356 | 0.002899874682 | 0.04780755131 | -1.956633351 |
| Ankrd12 | 0.4244921679 | 5.725891751 | 3.656442295 | 0.00292947153 | 0.04780755131 | -1.960151811 |
| Lrrc19 | 0.6975608216 | 7.004337053 | 3.656936442 | 0.002926717846 | 0.04780755131 | -1.95378436 |
| Fat3 | 0.9619823601 | -0.03175999496 | 3.655295719 | 0.002935871105 | 0.04781780009 | -1.545852643 |
| Sox17 | -0.6512844942 | 2.793756647 | -3.65400714 | 0.002943080238 | 0.04788288733 | -1.523207892 |
| Neto2 | -1.218360106 | -0.4024609601 | -3.653170684 | 0.002947769538 | 0.04788858851 | -1.721654491 |
| Plod1 | -0.5518362963 | 6.77451781 | -3.652798704 | 0.002949857361 | 0.04788858851 | -1.97841417 |
| Wfdc1 | -0.7310184789 | 1.855436874 | -3.651743158 | 0.002955790035 | 0.04793268649 | -1.474035692 |
| Wasf1 | -0.803776647 | 1.82880639 | -3.65027142 | 0.002964082172 | 0.04797472688 | -1.476572172 |
| Tmem50a | 0.3593818985 | 5.924367968 | 3.650140538 | 0.002964820738 | 0.04797472688 | -1.977147936 |
| Vwa5b2 | -0.7447561962 | 1.535185315 | -3.648094118 | 0.002976393045 | 0.04805762248 | -1.485201026 |
| Fdxr | -0.517180226 | 3.816866932 | -3.64824999 | 0.002975509989 | 0.04805762248 | -1.690947779 |
| Myo3b | 0.9694864931 | 4.460632272 | 3.647070793 | 0.002982197065 | 0.04809922387 | -1.740225331 |
| Bpgm | -0.5276951923 | 5.520469995 | -3.645848722 | 0.002989143403 | 0.0481550444 | -1.94442559 |
| Ssbp2 | -0.2741692079 | 4.684492982 | -3.643818271 | 0.003000721033 | 0.0481550444 | -1.833995207 |
| Senp6 | 0.4153461408 | 6.347056976 | 3.6436491 | 0.003001687701 | 0.0481550444 | -2.000346138 |
| Rbm15 | 0.4680770161 | 4.135556635 | 3.645024439 | 0.002993837977 | 0.0481550444 | -1.760820186 |
| Slc39a5 | 0.5455462047 | 6.215647371 | 3.643061923 | 0.00300504538 | 0.0481550444 | -1.995904663 |
| LOC101593466 | 1.69543338 | -0.2161069386 | 3.643135484 | 0.003004624523 | 0.0481550444 | -1.734490034 |
| Sumo1 | 0.2464339524 | 5.669657084 | 3.642086431 | 0.003010632013 | 0.04817641054 | -1.971652355 |
| Usp37 | 0.4101462825 | 5.501643918 | 3.641666631 | 0.003013039455 | 0.04817641054 | -1.966487045 |
| Gramd1b | 0.5275948538 | 5.281965326 | 3.641137495 | 0.003016076698 | 0.04817641054 | -1.96659766 |
| Podn | -0.8418030143 | 3.547942371 | -3.639479428 | 0.003025614183 | 0.04822537772 | -1.662130036 |
| Lmod1 | -0.5079051822 | 2.732910328 | -3.639949964 | 0.003022904465 | 0.04822537772 | -1.543644219 |
| Enpp5 | 0.3670520125 | 7.847298598 | 3.637828425 | 0.003035141508 | 0.04832554903 | -1.949190625 |
| Rtn4rl1 | -0.4287504978 | 4.74499853 | -3.637050041 | 0.003039643844 | 0.04834558399 | -1.834152255 |
| LOC123464079 | -1.050760679 | 4.210679458 | -3.636400922 | 0.003043403682 | 0.0483537794 | -1.748028644 |
| Nepro | -0.3677758066 | 3.547836387 | -3.634201325 | 0.003056179406 | 0.04843112738 | -1.647424877 |
| Znf366 | 0.75085487 | 2.219341268 | 3.634333195 | 0.003055411942 | 0.04843112738 | -1.523872335 |
| Gk | 0.8698387979 | 8.284257612 | 3.633884984 | 0.003058021262 | 0.04843112738 | -1.915554188 |
| Cldn12 | 0.3198537252 | 5.349180845 | 3.633144745 | 0.00306233562 | 0.04844226812 | -1.960761444 |
| Bcl2l11 | 0.547652418 | 4.019468084 | 3.632649467 | 0.003065225715 | 0.04844226812 | -1.772472649 |
| Npr1 | -0.4122093829 | 5.353707359 | -3.631884094 | 0.003069697356 | 0.04846154629 | -1.947236806 |
| Gfus | -0.2607964745 | 5.722067322 | -3.631155239 | 0.003073961808 | 0.04847751621 | -1.98807234 |
| Tacstd2 | -0.5957142582 | 7.173503948 | -3.630545607 | 0.003077533317 | 0.04848253597 | -2.016980838 |
| Bzw1 | 0.3019069889 | 7.263283609 | 3.628623317 | 0.00308882265 | 0.04855593191 | -1.997178882 |
| Gfm1 | 0.3167264589 | 7.021740339 | 3.628580055 | 0.003089077207 | 0.04855593191 | -2.008906422 |
| Dnajc5 | 0.357960274 | 5.961847135 | 3.628089239 | 0.003091966676 | 0.04855593191 | -2.027465317 |
| Irs2 | 0.9884199597 | 5.453469497 | 3.627457939 | 0.003095687233 | 0.04856318614 | -2.001129032 |
| Rab31 | -0.5668962256 | 5.221293161 | -3.625893741 | 0.003104925418 | 0.04864330257 | -1.951771008 |
| Lrfn3 | -0.5258629026 | 3.870787832 | -3.625488679 | 0.003107322287 | 0.04864330257 | -1.72836115 |
| Mfsd5 | -0.3513389154 | 4.874081455 | -3.624771671 | 0.003111569644 | 0.04865868038 | -1.885694159 |
| Elp5 | -0.3094475153 | 4.759436331 | -3.623821946 | 0.003117204634 | 0.04869570299 | -1.860109046 |
| LOC123462838 | -0.6646924288 | 1.476294217 | -3.622339047 | 0.003126023849 | 0.04873610345 | -1.523746226 |
| Zfp3 | 0.5355608978 | 2.385628095 | 3.622287433 | 0.003126331269 | 0.04873610345 | -1.562629208 |
| Chp2 | -4.054141919 | -1.219778871 | -3.621065845 | 0.003133616161 | 0.04879784017 | -1.960711317 |
| Reln | -1.036657456 | 2.274840561 | -3.620526119 | 0.003136840293 | 0.04879784017 | -1.532610916 |
| Ctc1 | -0.4571355322 | 4.333046774 | -3.618011692 | 0.003151905071 | 0.04898106494 | -1.805927459 |
| Sat1 | -0.4420849028 | 7.569466488 | -3.61686899 | 0.00315877566 | 0.04899062481 | -2.019873251 |
| Niban2 | -0.3835857186 | 6.971222965 | -3.616518119 | 0.003160888351 | 0.04899062481 | -2.044914367 |
| Nelfe | -0.3691013407 | 6.080579496 | -3.616270177 | 0.003162382143 | 0.04899062481 | -2.040020431 |
| Agap3 | -0.2233164843 | 5.978111598 | -3.614825792 | 0.003171098507 | 0.04907464283 | -2.037091811 |
| Rimoc1 | 0.8010163381 | 4.656911603 | 3.611942214 | 0.00318857298 | 0.0492938829 | -1.972992765 |
| Acox3 | -0.8446617368 | 7.725310802 | -3.610441498 | 0.00319770599 | 0.04938384702 | -2.013103275 |
| LOC123458055 | -1.173862152 | 0.1295952789 | -3.609618672 | 0.00320272479 | 0.04941015275 | -1.703164399 |
| Tfap2a | -0.6561977765 | 4.357214269 | -3.605931795 | 0.003225311231 | 0.04948445667 | -1.812382259 |

| | | | | | | |
|---|---|---|---|---|---|---|
| Adamtsl5 | -0.6317297446 | 6.024734726 | -3.605042374 | 0.00323078416 | 0.04948445667 | -2.048476296 |
| Smtnl2 | -0.5595691208 | 4.218400978 | -3.606308382 | 0.003222996796 | 0.04948445667 | -1.826028585 |
| Znf500 | -0.3928345943 | 4.98048139 | -3.605539579 | 0.003227723518 | 0.04948445667 | -1.931044582 |
| Asb3 | 0.4134313266 | 5.408859676 | 3.6061564 | 0.003223930649 | 0.04948445667 | -2.019924298 |
| Lmbrd2 | 0.4180878292 | 6.989393881 | 3.605868308 | 0.003225701582 | 0.04948445667 | -2.049373364 |
| Rap2a | 0.4554472813 | 4.584270651 | 3.606289852 | 0.003223110639 | 0.04948445667 | -1.914448882 |
| Stap2 | -0.8220950058 | 4.958988505 | -3.603956869 | 0.003237476449 | 0.04953604879 | -1.948854451 |
| Nln | 0.7662067714 | 6.160732556 | 3.602227955 | 0.00324816453 | 0.04964861128 | -2.070245024 |
| Osbp2 | -0.9067237946 | 5.534675392 | -3.600745376 | 0.003257358318 | 0.049687217 | -2.032601517 |
| Ovol1 | 0.8371738022 | 4.726288075 | 3.601005192 | 0.003255745233 | 0.049687217 | -1.915893473 |
